# Engineering cell fate with adaptive feedback control

**DOI:** 10.1101/2024.11.29.625974

**Authors:** Frank Britto Bisso, Giulia Giordano, Christian Cuba Samaniego

## Abstract

Engineering cell fate is fundamental for optimizing stem cell-based therapies aimed at replacing cells in patients suffering from trauma or disease. By timely administering molecular regulators—such as transcription factors, RNAs, or small molecules—in a process that mimics *in vivo* embryonic development, stem cell differentiation can be guided toward a specific cell fate. A significant challenge in scaling up these therapies is that such differentiation strategies often result in mixed cellular populations. While synthetic biology approaches have been proposed to increase the yield of desired cell types, designing gene circuits that effectively redirect cell fate decisions requires mechanistic insight into the dynamics of endogenous regulatory networks that govern decision-making. In this work, we present a biomolecular adaptive controller based on an Incoherent Feedforward Loop (IFFL)-like topology designed to favor a specific cell fate. This controller requires minimal knowledge of the endogenous network as it exhibits adaptive, non-reference-based behavior. The synthetic circuit operates through a sequestration mechanism and a delay introduced by an intermediate species, producing an output that asymptotically approximates a discrete temporal derivative of its input, provided there is a sufficiently fast sequestration rate. By allowing the controller to actuate over a target species involved in the decision-making process, a tunable, synthetic bias is created that favors the production of the desired species with minimal alteration to the overall equilibrium landscape of the endogenous network. Through theoretical and computational analysis, we provide design guidelines for the controller’s optimal operation, evaluate its performance under parametric perturbations, and extend its applicability to various examples of common multistable systems in biology.

## 1 Introduction

In multicellular organisms, phenotypical diversity is a direct consequence of cell decision-making, where cells in a transient state (e.g., undifferentiated) integrate and process information from their environment, and respond by committing to a specific fate [1, 2]. For example, during embryonic development, pluripotent stem cells are primarily responsible for generating cellular diversity. They give rise to the primordial germ layers—mesoderm, endoderm, and ectoderm—which then further specialize into all the tissues within our body [3]. Historically, the theoretical framework used to understand cell decision-making is the Waddington’s epigenetic landscape (Fig. 1-A). In this model, each decision is depicted as a bifurcation point, with branching pathways cascading until a stable equilibrium is reached, representing the cell’s ultimate fate (see Box 1). This visual representation provides insights for controlling cell fate: the use of ectopic molecular regulators (e.g., transcription factors, RNAs, and small molecules) can induce a cell to follow the path toward a specific equilibrium.

**Figure 1:**
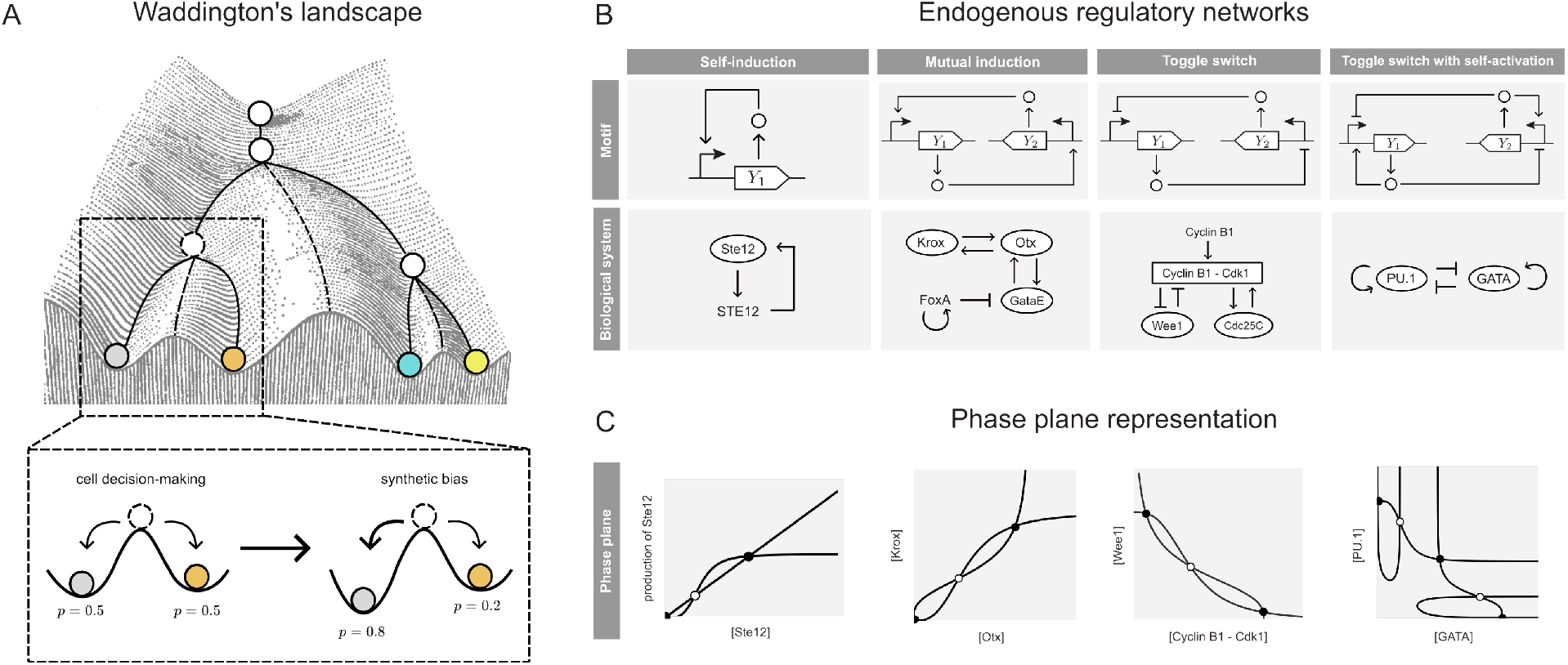
Multistability in endogenous biological systems. (A) Waddington’s epigenetic landscape, adapted from [4], illustrates how a pluripotent cell undergoes a cascade of decisions (bifurcations) that ultimately determine its fate (stable equilibrium). We consider the case of an unstable equilibrium from which a perturbation can lead to either of two stable equilibria with the same probability, and design an adaptive feedback controller that introduces a synthetic bias to favor one equilibrium (cell fate) over the other. (B) Well-documented networks that exhibit multistability: (I) self-induced upregulation of the Ste12 gene upon exposure to mating pheromones [5]; (II) mutual activation between Kros, Otx and GataE genes during endomesoderm specification [6]; (III) mutual inhibition in the “mitotic trigger” [7]; and (IV) mutual inhibition with positive self-activation in a gene regulatory network of hematopoietic differentiation [8]. (C) Phase plane representation of the endogenous regulatory networks detailed in (B), where the curves represent the nullclines of the associated dynamical system and their intersections correspond to the system equilibria, denoted as black dots when stable and as white dots when unstable

In the context of cell-based therapeutics for tissue replacement, developmental pathways are considered a discrete sequence of decision-making processes [9]. Stem cell differentiation can then be guided towards a desired fate by appropriately timing the activation or inhibition of key signaling pathways to mimic *in vivo* development. This approach led to the development of pancreatic islet replacement therapies for restoring normal glycemic control in diabetic patients, with currently an FDA-approved treatment for type 1 diabetes [10], and a promising case report for diabetes type 2 [11]; as well as stem cell-derived dopaminergic neuron replacement as an alternative treatment to Parkinson’s disease, with ongoing clinical trials [12, 13]. A current challenge in scaling up the manufacturing of these therapies is that guided differentiation strategies often lead to mixed populations, producing both desired cell types and off-target cells due to diverging cell fates. This issue can be traced back to specific progenitor (undifferentiated) cells that can adopt multiple fates (i.e., multiple stable equilibria) with an inherent bias towards certain fates, resulting in the heterogeneity reported in the literature. For instance, in the later stages of the differentiation protocol for pancreatic islet production, endocrine progenitors can differentiate into *β* cells (the desired cell fate) but also into *α* and *δ* cells [14]. Similarly, during an early stage of the differentiation protocol for dopaminergic neuron production, stem cells can differentiate into ventral midbrain progenitors (which can subsequently be induced into dopaminergic neurons) as well as into ventral hindbrain progenitors and midbrain-hindbrain boundary cells [15]. Additionally, in later stages of differentiation for both examples, single-cell analyses have identified more off-target cell types, which are often the result of a previous, non-optimal differentiation step, leading to proliferative cells and various undesired types, such as non-endocrine cells (e.g., duct-like and liver-like cells [16]) and non-midbrain neurons (e.g., serotonin and GABAergic neurons [15, 17]).

The optimization of these protocols to increase the yield of the desired cell fate (e.g., *β* cells or midbrain dopaminergic neurons) can potentially be addressed using biomolecular feedback controllers. Rather than titrating the concentrations of molecular regulators within a certain range, suitably designed genetic circuits can bias cell decision-making by dynamically adjusting the concentrations of these regulators within the appropriate range and at the right time, given the correct reference [18, 19]. Conveniently, the mathematical framework used for designing these circuits is the same as that applied to study cell decision-making (see Box 1). Thus, the challenge is reduced to an engineering problem: for a given initial condition, how can we design a controller to drive the system to a desired stable equilibria? Although this problem has been extensively explored in control theory [20], the main limitation when translating it to a biological context is the need for mechanistic insight into the regulatory network dynamics that govern cell fate (Fig. 2–A). The stoichiometry of cell fate drivers alone is insufficient for efficient, guided differentiation: both the relative expression levels and the duration of expression must also be considered [21]. Furthermore, quantifying the impact of introduced exogenous perturbations on the equilibrium remains a challenge. Minimal perturbations may not induce a bias towards the desired cell fate, whereas larger perturbations could alter the number of equilibrium points, potentially making some cell fates inaccessible [22, 23].

**Figure 2:**
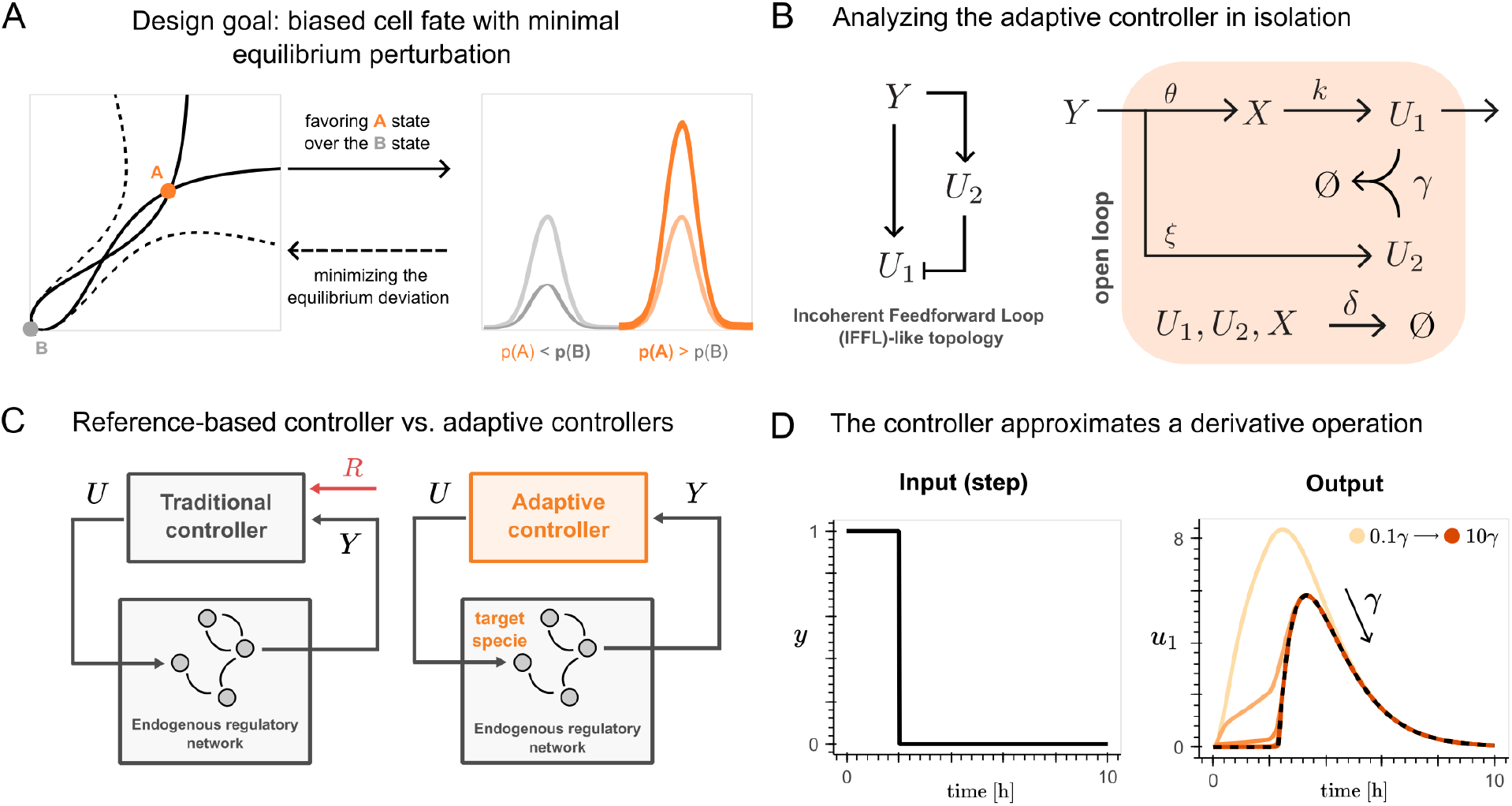
Building an adaptive controller. (A) The objective of this work is to design a control strategy that favors one stable equilibrium point (orange) with respect of another (grey), corresponding to a skewed (asymmetric) probability distribution; and that minimizes the perturbation to the equilibrium (black lines), thus avoiding the lost of the multistable behavior (dashed lines) (B) The chemical reactions network that implements the adaptive controller has a topology akin to that of an Incoherent Feedforward Loop (IFFL). (C) Feedback controllers generate a control signal U based on an error metric estimated by comparing the current output of the process being controlled, Y , and a desired set-point value. The feedback controller aims to minimize this error. Typically, controllers require a reference species (R) as the set-point, which, however, is often unknown for endogenous networks. In contrast, the proposed adaptive controller can directly estimate the reference from the dynamics of the endogenous regulatory network. (D) If we analyze the controller in isolation, following Eq. (4), by tuning the adaptive metric 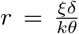 so that r = 1, the adaptive controller computes a low-pass filter that introduces a delay, followed by a derivative operation. For increasing values of the sequestration rate γ, the time evolution of u_1_ obtained by numerically solving Eqs. (1)-(3) quickly converge to the approximation in Eq. (4), represented as black dashed lines in the “Output” plot.

In this work, given a gene regulatory network that exhibits multistability, where the stable equilibria are associated with different possible cell fates, we propose an adaptive biomolecular feedback controller that introduces a synthetic bias into the decision-making process and thus favors a specific cell fate over the others, while minimally altering the corresponding equilibrium value (see the schematic in Fig. 2-A). Through theoretical analysis, we delineate the requirements needed for generating a biased outcome that favours the desired cell fate, and provide design guidelines for the correct operation of the controller. Additionally, we provide computational simulations to address the performance of the controller under parameter variations within the endogenous regulatory network; and extend the application of the controller to the motifs shown in Fig. 1–B.

### Box 1: Decision-making and cell fate: insights from dynamical systems theory

Dynamical systems theory provides a powerful mathematical framework to formalize Waddington’s idea of the epigenetic landscape. Within this framework, the state of a cell—characterized by its unique gene expression profile—can be visualized in a coordinate system known as the *phase plane*, where each axis represents the expression level of a specific gene that defines the cell’s state (Fig. 1–C). Changes in gene expression are represented as the movement of a point along a trajectory in this plane. However, these trajectories are not arbitrary: they are constrained by the underlying gene regulatory networks [2, 24, 25]. Some of these networks exhibit a property called *multistability*, which allows for the emergence of multiple stable equilibrium points (also known as attractors), towards which trajectories eventually converge. In the context of Waddington’s landscape, each “valley” corresponds to one of these equilibrium points, representing a distinct cell fate. Namely, bistable and tristable systems are among the most studied multistable networks in synthetic biology (Fig. 1–B). In this context, cell decision-making can be conceptualized as the process by which a cell reaches one of these equilibrium points.

Recent advancements in algorithms for estimating dynamic trajectories from single-cell RNA sequencing snapshots have significantly improved our ability to identify key genes responsible for specific cell fates within high-dimensional gene regulatory networks [23, 26]. However, the intricate nature of these networks often makes it difficult to understand the mechanisms by which gene interactions lead to particular cell fates [27]. As an alternative, a bottom-up approach examines simpler gene regulatory motifs, such as self-activation, mutual induction, and mutual inhibition, and builds up more elaborate dynamics from these fundamental building blocks (Fig. 1–B). These motifs have been extensively studied both theoretically and experimentally, primarily in the context of differentiation [6, 8, 28]; and although initially proposed for studying developmental processes, this framework can be applied to virtually any biological process involving multistability, including cell cycle entry [7], quiescence [29], macrophage activation [30], and yeast mating [5].

Cell decision-making is inherently a single-cell event and thus exhibits stochasticity [31]. Consequently, for a given initial cellular state (i.e., a coordinate in the *phase plane*), there is a probability distribution corresponding to each potential cell fate. During development, these distributions are often skewed, favoring one cell fate over others. While multistability can explain the emergence of diverse cell types from a single progenitor, the varying proportions of cell types within a tissue reflect this inherent bias [32, 1, 33]. In our work, we designed a synthetic gene circuit to replicate this effect by introducing a “synthetic bias” in cell decision-making, as illustrated in Fig. 1–A.

## 2 Results

Throughout the manuscript, we denote species with uppercase letters (e.g. *Y* ) and their concentrations with the corresponding lowercase letters (e.g. *y*). The time derivative of *y* is denoted by 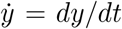, while the steady-state concentration of a chemical species *Y* is denoted by 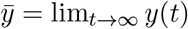.

### 2.1 An adaptive controller enables a biased cell fate

#### 2.1.1 The controller and its adaptive nature

To generate a biased cell fate, defined as a probability distribution that is skewed towards the production of the target species, with minimal perturbation to the equilibrium landscape, we propose a controller whose topology, similar to an Incoherent Feedforward Loop (IFFL) and building upon previous work [34, 35, 36], is shown in Fig. 2-B. An input *Y* produces at rate constant *θ* the species *X*, which in turn produces *U*_1_ at rate constant *k*. At the same time, *Y* produces the species *U*_2_ at rate constant *ξ*, which further downregulates the expression of *U*_1_ through sequestration at rate constant *γ*. Ultimately, every species decays at rate *δ*. This leads to the following chemical reactions:

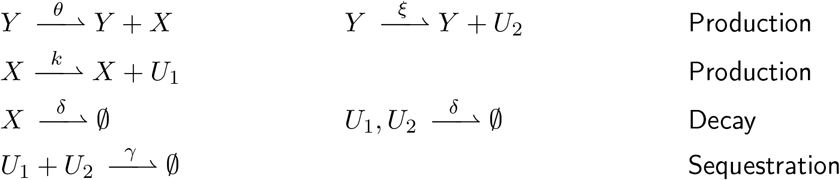

We use the law of mass action to derive a set of Ordinary Differential Equations (ODEs) that describe the dynamics of the species concentrations in the controller.

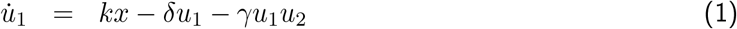

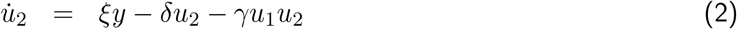

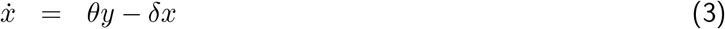

Unlike reference-based feedback controllers, the proposed network does not require a reference species, since it can adaptively estimate the signal to be tracked directly from the dynamics of the process (as illustrated in Fig. 2-C, see also [36]). To understand its steady-state behavior, we can analyze the controller in isolation (Fig. 2-B) and derive an approximation of its dynamics in the fast sequestration regime (namely, when *γ*→ ∞ ), as detailed in the Supplementary Material (cf. also [34]). Then, through a suitable coordinate transformation, we can approximate the dynamics of *U*_1_ as

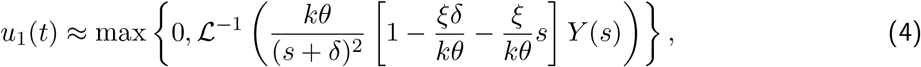

where *Y* (*s*) is the Laplace transform of the input to the controller and *ℒ*^−1^ denotes the inverse Laplace operator. Let 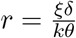 denote the *adaptive metric*, proposed as a design parameter. Substituting *r* = 1 into Eq. (4), we observe that the controller asymptotically includes a term resembling a low-pass filter, 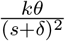, and a term resembling a temporal derivative, 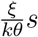, and whose overall effect is illustrated in Fig. 2-D. This behavior was previously demonstrated to function effectively as an adaptive controller [36]. Applied to a multistable endogenous regulatory network system, the controller will steer the system trajectories towards a desired fate (i.e., a specific equilibrium point) at the price of small perturbations in the expression of the target species. Furthermore, due to the derivative-like behavior in the fast sequestration regime, the equilibrium is minimally perturbed, since the effect of the controller converges to zero asymptotically, as the system reaches the desired steady state (proof available in the Supplementary Material).

#### 2.1.2 Toggle switch: model description

As a first example of biological multistable system, we consider the canonical *toggle switch* [37], a motif consisting of the mutual inhibition of two genes that respectively produce transcriptional repressors *Y*_1_ and *Y*_2_, as shown in Fig. 3-A1. The chemical reactions

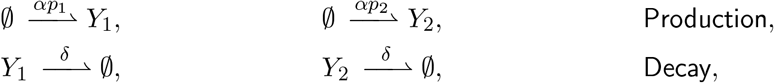

with 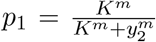 and 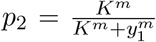 , allow for two possible stable equilibria to which the system variables *y*_1_ and *y*_2_ could converge (Fig. 3-A2): either *Y*_1_ is maximally expressed and *Y*_2_ is minimally expressed, or vice versa, resulting in a bimodal probability distribution (Fig. 3-A3).

**Figure 3:**
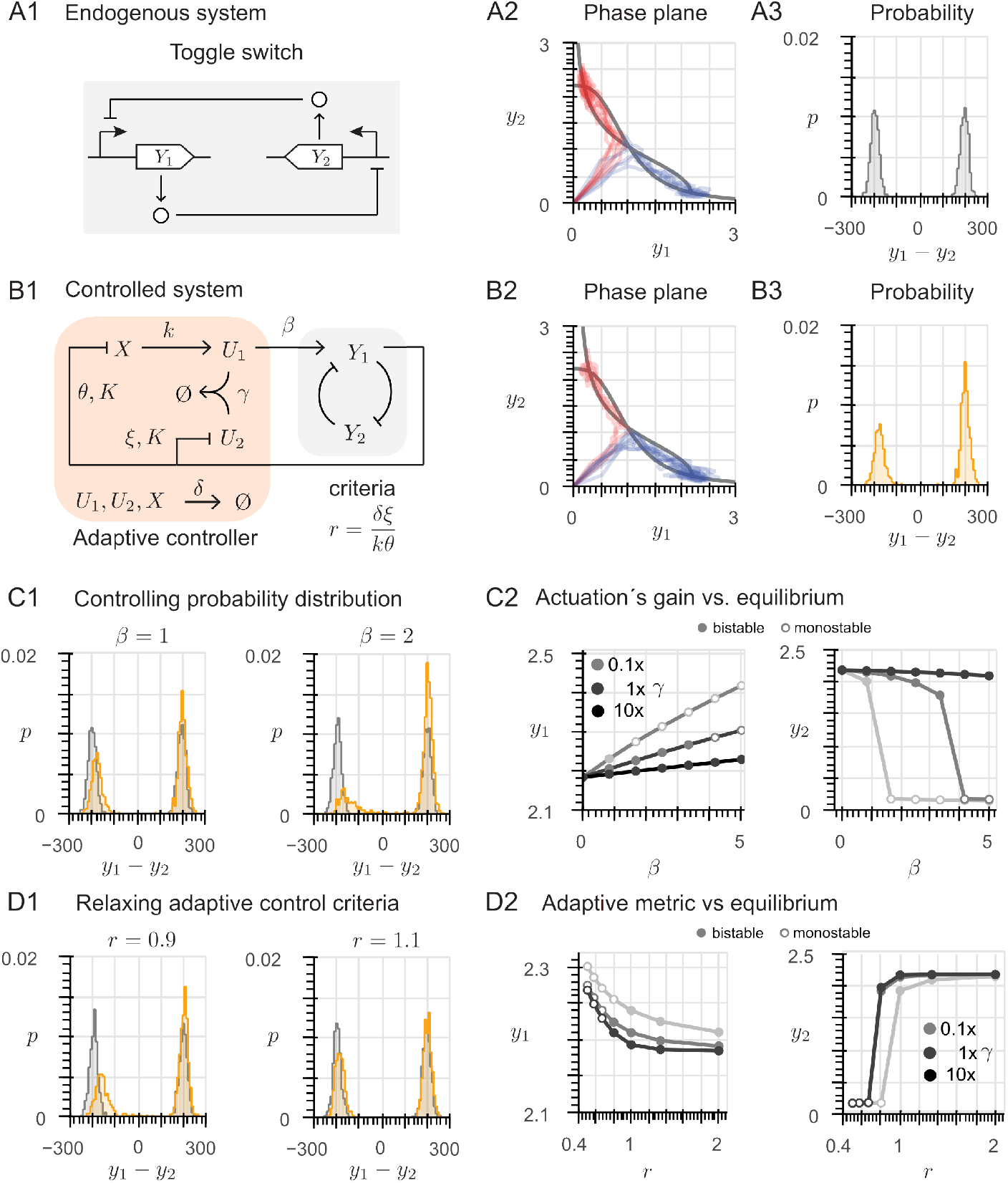
The adaptive controller enables a biased cell fate. (A1) Architecture of the toggle switch, formed by two mutually inhibiting transcription factors Y_1_ and Y_2_. (A2) For the toggle switch, in the phase plane, the nullclines (black lines) are shown along with 1000 trajectories starting from different initial conditions; each trajectory converges either to the stable equilibrium point with high expression of Y_1_ and low expression of Y_2_ (blue trajectories) or to the stable equilibrium point with high expression of Y_2_ and low expression of Y_1_ (red trajectories). (A3) The trajectories in (A2) converge essentially with equal probability to either of the stable equilibria, resulting in an unbiased bimodal probability distribution. (B1) Architecture of the toggle switch with the proposed adaptive controller. (B2) For the controlled system, in the phase plane, the nullclines (black lines) are shown along with 1000 trajectories starting from different initial conditions; more trajectories converge to the stable equilibrium point with high expression of Y_1_ (blue) and less to the stable equilibrium point with high expression of Y_2_ (red). (B3) The controller yields a biased cell fate: the trajectories in (B2) converge with higher probability to the equilibrium where the production of Y_1_ is favored over that of Y_2_. (C1) Increasing the control gain β increases the bias in the cell fate, leading to a larger imbalance in the probability distribution. (C2) Equilibrium values for increasing control gain β, for different values of the sequestration rate γ. (D1) Effect of a 10% deviation of the adaptive metric 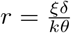 from its nominal value 1: a biased cell fate is still generated at the price of a small alteration of the equilibrium values. (D2) Equilibrium values for varying adaptive metric 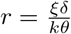 , for different values of the sequestration rate γ.

To model the controlled system (see schematics in Fig. 3-B1), we can update the chemical reactions describing the controller as follows:

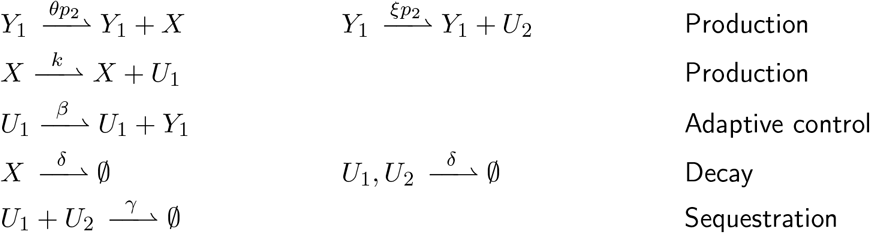

We use the law of mass actions to derive the ODEs of the controlled system.

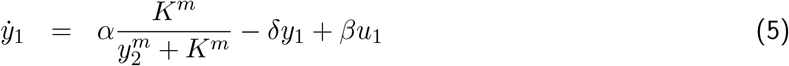

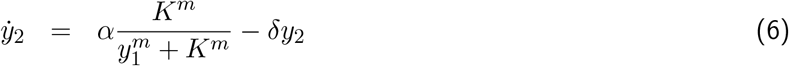

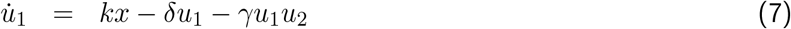

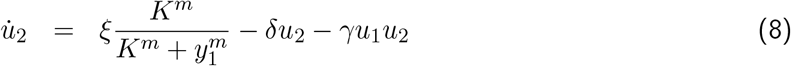

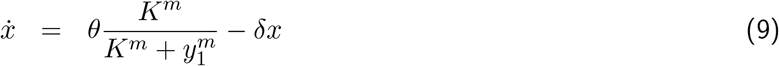

where Eq.(5) and (6) model the controlled toggle switch, with the control enforced by the additive term *βu*_1_ in Eq.(5), while Eqs.(7)-(9) describe the controller dynamics.

#### 2.1.3 Dynamic properties: biased cell fate and minimum equilibrium alteration

To validate the statements derived from the theoretical analysis of Eq.(4), we solved Eqs.(5)-(9) numerically. Enforcing the controller preserves the bistability of the toggle switch system, as the nullclines associated with 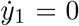 and 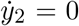 still have three intersections, corresponding to two stable equilibria (one with low *y*_1_ and high *y*_2_, and the other with high *y*_1_ and low *y*_2_) and one intermediate unstable equilibrium. Thus, also the controlled system can converge to either of the two stable equilibria, as shown by the blue and red trajectories in Fig. 3-B2. However, the symmetry of the probability distribution of the toggle switch in isolation is disrupted: in the controlled system, the production of *Y*_1_ is favoured with respect to that of *Y*_2_, as shown in Fig. 3-B3. The bias in the outcome can be further increased with higher values of the control gain *β*, as shown in Fig.3-C1, at the expense of a higher alteration of the equilibrium value. Above a certain threshold (in this case, *β >* 4 for the parameters detailed in the Methods section), the system transitions to monostability, as illustrated by the white circles in Fig.3-C2. Conversely, increasing the sequestration rate *γ* reduces deviations from the equilibrium value and prevents the transition to monostability, as indicated by the black circles in Fig.3-C2 for a ten-fold increase in *γ*. However, this adjustment also results in a decreased probability of producing the species *Y*_1_, highlighting the trade-off between maintaining system stability and achieving desired output levels (see Fig. S3. Further details are available in Supplementary Material 2.1).

As mentioned before, the adaptive behavior of the controller depends on the relationship between its rate constants, which must be such that 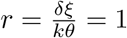. As shown in Fig. S4, if *r >* 1, due to increasing the degradation rate *δ* and/or the maximum production rate *ξ* of *U*_2_, the biasing capacity is lost, while the equilibria remain almost unperturbed. Conversely, if *r <* 1, due to increasing the maximum production rate *θ* of *X* and/or the production rate *k* of *U*_1_, the equilibrium landscape is significantly altered, and bistability is eventually lost, but the probability of the biased output increases. Since fine-tuning the kinetic parameters *δ, ξ, k* and *θ* so that *r* is exactly 1 is difficult in practice in an experimental setting, we provide numerical simulations to assess the effect of an error margin of *±* 10% (Fig. 3-D1). Within this range, for a fixed value of the control gain (*β* = 1), bistability is preserved and the maximum alteration in the equilibrium value is about 8%. Further details are available in Section 2.2 of the Supplementary Material.

#### 2.1.4 The adaptive controller is effective in both negative and positive feedback architectures

Enforcing the adaptive controller in the toggle switch, as described in Fig. 3-B1, results in an overall increase in the concentration of *Y*_1_. Therefore, species *X* will be subject to an increased inhibition, since *Y*_1_ acts as a repressor; hence, this is a negative feedback control architecture. However, as we now show, the effectiveness of the proposed adaptive controller is independent of the type of feedback provided by the endogenous gene regulatory network. Let us extend our analysis to a positive feedback control architecture. For the toggle switch, this implies that the feedback to the controller originates from the non-target species *Y*_2_ (see Fig. S5-B1), and the chemical reactions that describe the feedback interaction between the endogenous regulatory network and the input species *X* and *U*_2_ of the adaptive controller become

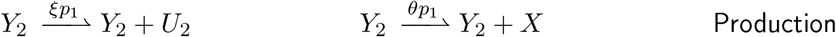

with 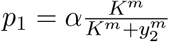 . Consequently, in the ODE system that models the kinetics of the controlled toggle switch, Eqs. (8) and (9) become

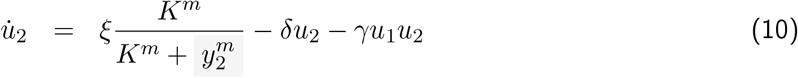

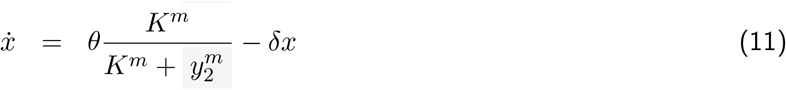

This alternative control strategy also yields a biased cell fate, favoring the production of species *Y*_1_ (Fig. S5-B2 and B3). Analogously to the negative feedback architecture, higher values of the control gain *β* increase the bias in the cell fate, further favoring higher production of the target species *Y*_1_ (Fig. S5-C1), while a larger sequestration rate reduces the alteration in the equilibrium values (Fig. S5-C2). However, the positive feedback architecture requires larger values of the control gain to achieve a probability distribution similar to that obtained with the negative feedback. On the other hand, the positive feedback architecture reduces the alteration in the equilibrium values, and exhibits a wider range of values of the adaptive metric *r*, from − 20% to +10% of the nominal value 1, for which the performance remains satisfactory (Fig. S5-D1 and D2, respectively).

In general, once we identify a target species for our control action, as long as the endogenous system exhibits multistability, in the fast sequestration regime and with adaptive metric *r* ≈ 1 (within the aforementioned error margin), the proposed controller can yield a biased cell fate, regardless of the feedback architecture (Fig. S6-A). However, by comparing both architectures, we can observe some specific features of each controller (Fig. S6-B). The negative feedback architecture allows to reach with higher probability the equilibrium that favors the production of the target species even with relatively small values of the control gain, but causes a higher alteration of the equilibrium values, even leading to loss of the system’s bistability. Conversely, the positive feedback architecture preserves the system’s bistability for a wider range of values of the control gain, and leads to a smaller alteration of the equilibrium values, but the ability to induce a biased cell fate is significantly reduced, for the same control gain value, compared to negative feedback. As a rule of thumb, the negative feedback architecture typically yields 80%-20% distributions, whereas the positive feedback architecture yields 60%-40% distributions.

#### 2.1.5 Changing the input species

To assess how the choice of the input species (so far, *U*_1_) affects the controller’s ability to generate a biased cell fate, we changed the input species to *U*_2_. As it can be shown [38], the negative (respectively, positive) feedback control architecture with input *U*_2_ has a similar steady-state behavior to the positive (respectively, negative) feedback control architecture with input *U*_1_. Numerical simulations for both architectures further validate this statement (see Fig. S7 for the negative feedback and Fig. S8 for the positive feedback architecture), wherein the dynamic properties are conserved.

### 2.2 The adaptive controller exhibits robustness under kinetic mutations

Are the aforementioned dynamic properties of our proposed controller robust to parameter variations? To study the robustness of the controller, we simulated genetic mutations that impact the kinetics of the endogenous regulatory network by affecting the binding between the repressors, *Y*_1_ and *Y*_2_, and their corresponding DNA binding sequences [39]. For the toggle switch, this amounts to modifying the maximum production rate (*α*) for each species in the chemical reactions associated with the production of the repressors, resulting in the modified ODEs

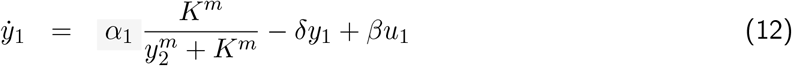

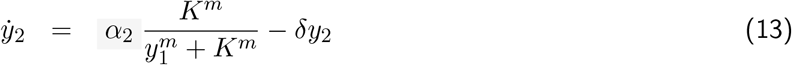

while the ODEs that describe the adaptive controller remain the same as Eqs. (7), (8) and (9). For a more systematic analysis, we consider the *ratio α*_2_*/α*_1_ and study four different cases: a *ratio* larger than 1, either due to increasing *α*_2_ or to decreasing *α*_1_, and a *ratio* smaller than 1, either due to decreasing *α*_2_ or to increasing *α*_1_. We consider *robust* a network motif that maintains the same number of equilibria as the original multistable system for a given range of *α*_2_*/α*_1_ values, and we wonder which is the maximum range of parameter variations for the bistability of the toggle switch is preserved. Since the most straightforward approach to compensate for an imbalance in the maximum expression rate between the two species is the introduction of a positive feedback, the robustness of the adaptive controller will be compared to that of a self-activation motif with chemical reactions

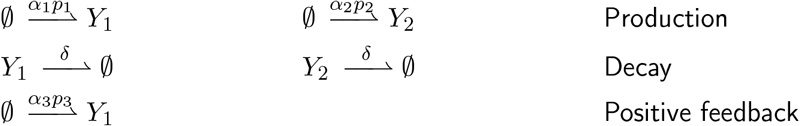

with 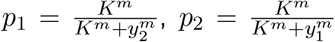 and 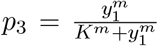 . Under the law of mass action, these reactions correspond to the ODEs

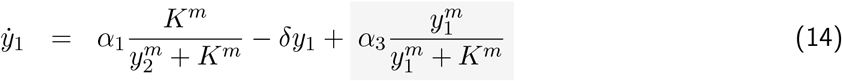

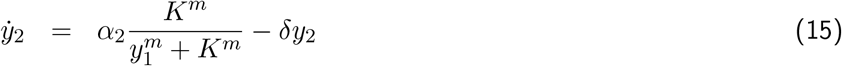

#### 2.2.1 The positive feedback architecture is more robust to parameter variations

Simulations of the behavior of the toggle switch under parameter variations are shown in Fig. 4-B1 to B4. Inherently, this motif is robust for *α*_2_*/α*_1_ ∈ [0.8, 1.8] when altering *α*_2_ (see Fig. 4-B1 and B2), and for *α*_2_*/α*_1_ ∈ [0.6, 1.2] when altering *α*_1_ (see Fig. 4-B3 and B4). For the toggle switch with the self-activation motif, since the addition of the positive feedback is done to *Y*_1_, the effect on the equilibrium is more apparent when *α*_2_*/α*_1_ *>* 1, both due to decreasing values of *α*_1_ and due to increasing values of *α*_2_. Increasing production of *Y*_1_ compensates for the higher abundance of *Y*_2_, thus preserving bistability for much larger values of the ratio with respect to the toggle switch in isolation (which, in contrast, loses bistability either due to increasing values of *α*_2_, as shown in Fig. 4-C1 and C2, in red and grey, or to decreasing values of *α*_1_, as shown in Fig. 4-C3 and C4, in red and grey). Conversely, when *α*_2_*/α*_1_ *<* 1, the higher abundance of *Y*_1_ is amplified by the positive feedback, resulting in a smaller robustness margin with respect to the toggle switch in isolation. For instance, with the positive feeback, bistability is lost for *α*_2_*/α*_1_ *<* 1 for decreasing values of *α*_2_ (see Fig. 4-C1 and C2, in red), and increasing values of *α*_1_ (see Fig. 4-C3 and C4, in red).

**Figure 4:**
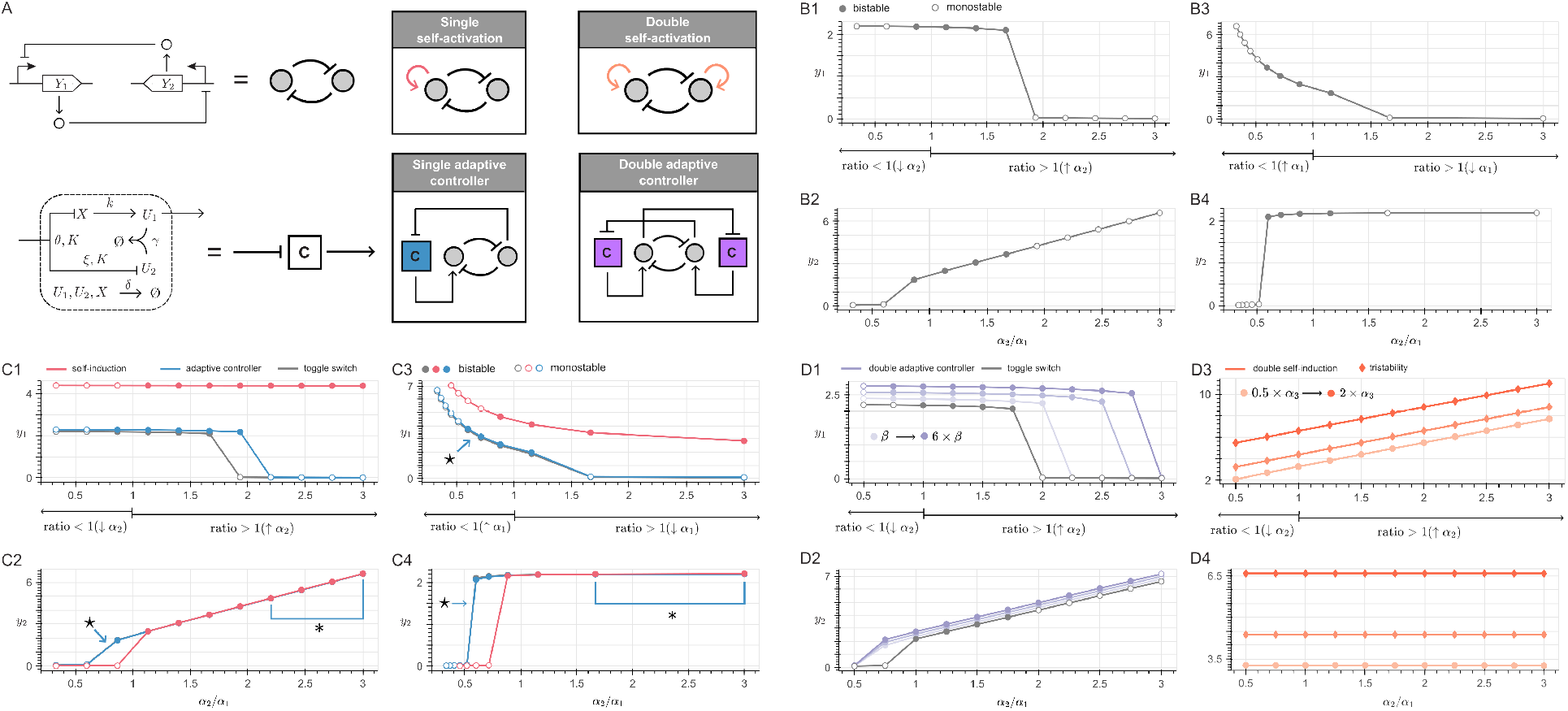
The adaptive controller with the positive feedback architecture is more robust to parameter variations. (A) Schematics of the circuits considered for robustness analysis: the adaptive controller with the positive feedback architecture is compared to the toggle switch with a self-activation motif (in the same species that is the control target), to a double adaptive controller with the positive feedback architecture, acting on both species Y_1_ and Y_2_, and to a toggle switch with a self-activation motif on each species. (B1) and (B2): effect on the alteration of the equilibrium value of the isolated toggle switch’s Y_1_ species (B1, for the equilibrium associated with high expression of Y_1_) and of the equilibrium value of Y_2_ species (B2, for the equilibrium associated with high expression of Y_2_) of parameter variations that alter the ratio α_2_/α_1_ due to changes in α_2_. (B3) and (B4): same analysis as in (B1) and (B2), due to changes in α_1_. (C1) to (C4): same analysis as in (B1) to (B4), considering the toggle switch in isolation (grey), with the adaptive controller with positive feedback architecture (blue), and the toggle switch with self-activation motif (red). For (C2) and (C4), the points highlighted with the (*) symbol correspond to monostability of the blue line, overlapped by the red line, since deviations to the equilibrium are within a similar magnitude. Likewise, for (C2) to (C3), the blue lines pointed with the (⋆) symbol overlap the grey lines shown in (B1) to (B4), and exhibit similar stability. (D1) and (D2): increasing the gain of the adaptive controller (β) enlarges the range of values of the ratio for which bistability is obtained. For comparison, (D3) and (D4) characterize the alteration of the equilibrium values for a toggle switch with a self-activation motif on each species. Simulations were performed by evaluating the equations of the nullclines for the toggle switch with a simple adaptive controller (Eqs.(13) and (13), the toggle switch with a single, self-activation motif (Eqs. (14) and (15)), the toggle switch with the pair of controllers (Eqs. (16) and (17)), and the toggle switch with a self-activation motif on each species (Eqs. (18) and 19). The same nominal parameters as in Fig. 3 were used, except for the maximum production rates α_1_ and α_2_, which were varied accordingly.

We considered the positive feedback architecture for the present analysis since it proved to be more robust than the negative feedback architecture (Fig. S9). Similarly to the self-activation motif, the adaptive controller also increases the abundance of species *Y*_1_, due to the additive terms in Eqs. (5), respectively. However, when considering parameter variations, the adaptive controller can preserve the system’s bistable behaviour within a smaller parameter range, due to the derivative effect (and the corresponding magnitude of *u*_1_). For instance, see the blue and red curves in Fig. 4-C1 to C4 for *α*_2_*/α*_1_ *>* 1 for increasing values of *α*_2_ and decreasing values of *α*_1_. Additionally, when comparing the effects of the self-activation motif and the adaptive controller on the equilibrium values as the ratio of *α*_2_*/α*_1_ increases, in all cases we observe negligible alterations when considering a fixed control gain *β* = 1, within a deviation *<* 5% with respect to the isolated toggle switch (the corresponding effect on the nullclines is shown in Fig. S13). In comparison, the self-activation motif preserves the equilibrium value of the species not involved in the positive feedback (*Y*_2_), while the auto-induced species (*Y*_1_) has a significantly different new equilibrium value, as suggested in Fig. 3-C1 and C3, and validated through nullclines computations in Fig. S16.

In particular, when *α*_2_*/α*_1_ *>* 1 due to changes in *α*_2_ and when *α*_2_*/α*_1_ *<* 1 due to changes in *α*_1_, the range of values of the ratio for which bistability is preserved can be enlarged by increasing the control gain of the adaptive controller, at the expense of a larger alteration of the equilibrium values, which increases with the magnitude of the control gain (Fig. S10-A). Similarly, increased robustness when *α*_2_*/α*_1_ *>* 1 due to changes in *α*_1_ and *α*_2_*/α*_1_ *<* 1 due to changes in *α*_2_ can be achieved by drastically increasing the sequestration rate *γ* (ten-fold for a noticeable change, as shown in Fig. S11-A). However, as an intrinsic limitation, high values of the sequestration rate *γ* reduce the magnitude of the control action. Consequently, the adaptive controller exhibits the same robustness margin as the isolated toggle switch (Fig. 3-C1 and C2, as the grey and blue curves overlap). The same conclusions hold for the adaptive controller with a negative feedback configuration under parameter variations, although a narrower range of viable ratios *α*_2_*/α*_1_ is observed in this case (Fig. S10-B and Fig. 11-B).

#### 2.2.2 An additional adaptive control loop improves overall robustness

Either the control gain *β* or the sequestration rate *γ* can be fine-tuned to increase robustness, but in a way that depends on the type of mutation (i.e., on which of the system parameters is varied). A single controller with a positive feedback architecture can only minimize the alteration of the equilibrium values corresponding to the species not affected by the mutation (namely, *Y*_1_ for changes in *α*_2_ and *Y*_2_ for changes in *α*_1_). Simulating how the nullclines are altered by increasing the positive feedback strength *β* in the adaptive controller and *α*_3_ in the self-activation motif (as shown in Fig. S12 - S16) suggested that the limitations previously discussed can be mitigated by adding to each of the two motifs a second, symmetric feedback loop that targets the other species (see the schematics in Fig. 4-A). Therefore, we simulated an additional adaptive controller with a double positive feedback architecture, which applies another control input to *Y*_2_, driven by species 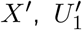and 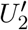 that share the same production rate constants as *X, U*_1_ and *U*_2_. The corresponding chemical reactions are

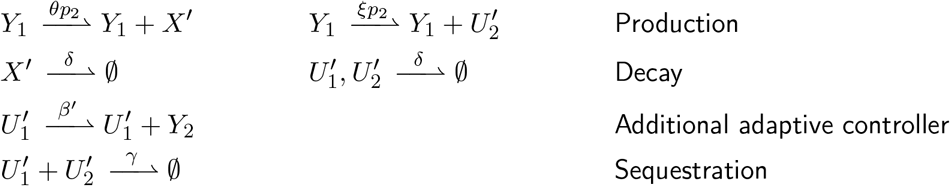

with 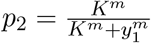 , and the system can be modeled as

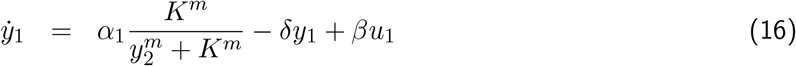

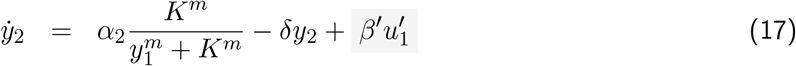

along with ODEs that describe each controller, which are similar to Eqs. (7)-(9), with 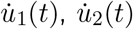 and 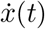 depending on *y*_2_, and 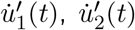 and 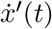 depending on *y*_1_, as detailed in the Supplementary Material.

Analogously, we can model the addition of another self-activation motif acting on the non-target species, with the same maximum expression rate *α*_3_, described by the chemical reactions

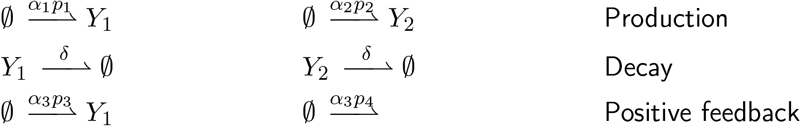

with 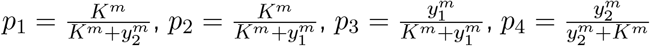 . Then, the dynamics of the toggle switch with a self-activation motif on each species is described by the ODEs

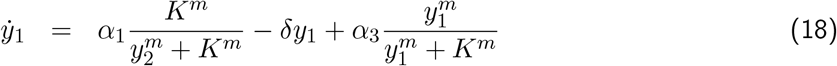

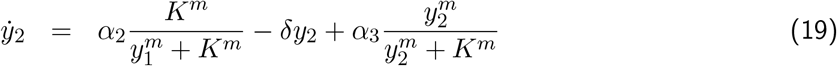

Comparative simulations reveal that the toggle switch with self-activation motifs on both species is so robust that bistable behavior becomes independent of the ratio *α*_2_*/α*_1_ (see Figs. 3-E1 and E2). However, this strategy is only effective when *α*_3_ ≪ *α*_1,2_; with higher values of *α*_3_, new intermediate equilibria appear, leading to loss of bistability in favor of tristability (see the curves with diamonds in Fig. 3-E1 and E2, as well as Fig. S17). On the other hand, the addition of another adaptive controller with a positive feedback architecture enables robust performance for an extended *α*_2_*/α*_1_ range, regardless of the parameter that is being varied. As observed before, increasing the gain of both controllers (assuming *β* = *β*^*′*^ for simplicity) can maintain a bistable behaviour for an increased range of values of *α*_2_*/α*_1_: the configuration with two controllers in a fast sequestration (large *γ*) and high gain (large *β*) regime exhibits bistability for values of *α*_2_*/α*_1_ from 0.7 to 2.7, with a maximum 25% alteration of the equilibrium value, and without the generation of new, intermediate equilibrium points (see Fig. 3-D1 and D2, as well as the nullcline analysis in Fig. S19).

### 2.3 Adaptive controller: general applicability

To outline a general design principle for the application of the adaptive controller to any type of regulation and to multistable networks with more than two stable equilibria, we proposed the mutual activation motif and the toggle switch with self-activation as study cases, respectively.

#### 2.3.1 Applying the adaptive control principle to a different system

The mutual activation (double activation) motif in Fig. 4-A1 has a similar architecture to the toggle switch (double repression); in both cases, we have an overall positive loop, which is known to be associated with bistability [40, 41]. Its bistable behavior arises as well from the mutual interaction between species *Y*_1_ and *Y*_2_, but with an opposite effect, since both species are inducers. Consequently, two stable, equilibrium points emerge: either both species are highly expressed, or neither of the two (Fig. 4-A2). Favoring high expression for both species is trivial, since any increase in the abundance of one species is amplified by the mutual activation loop. We modified the controller to favor minimal expression for both species, by enforcing the chemical reactions

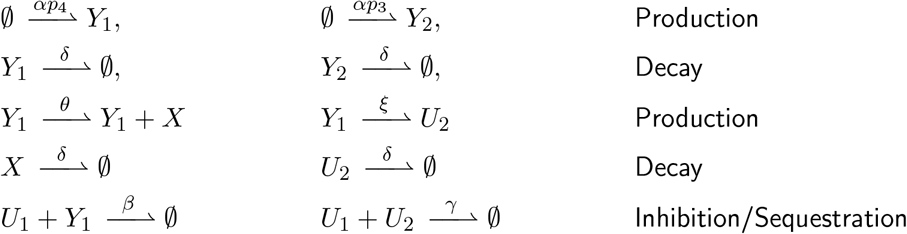

with 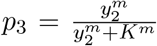 and 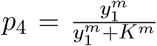 . The first two reactions are associated with the mutual activation motif, all the others with the adaptive controller in a negative feedback architecture. Adopting a Michaelis-Menten approximation to model *U*_1_-mediated degradation of species *Y*_1_ [42] leads to the ODEs

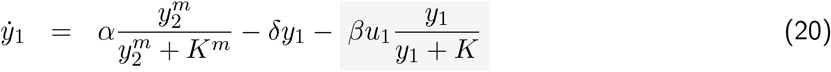

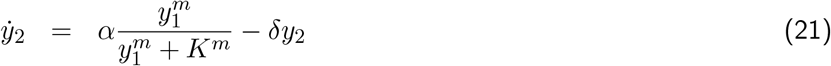

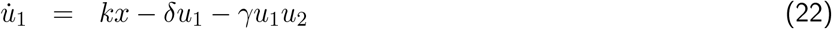

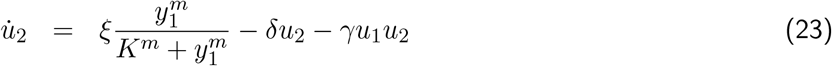

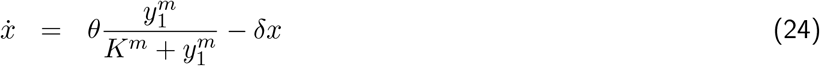

The resulting architecture of the adaptive controller is depicted in Fig. 5-B1. As in the toggle switch case, the bistability of the mutual activation motif is preserved when enforcing the controller (compare Fig. 5-A2 and B2). Hence, the system trajectories can still converge to either of the two stable equilibria (high production of both species vs. no production of both species), resulting in a bimodal distribution, which is however not symmetric (Fig. 4-A3). Enforcing of the controller favors the equilibrium at which zero molecules of both species are produced (Fig. 4-B3). The same properties hold for the positive feedback architecture, as well as by selecting *U*_2_ as the input specie (see Fig. S20-A1 to A4, and B1 to B4). In general, high control gains (*β >* 3) were needed to achieve a noticeable level of inhibition, which resulted in a moderate alteration of the equilibrium value associated with the high expression of both species (Fig. 5-B2 and B4). By adding another adaptive controller that targets *Y*_2_, a similar level of inhibition can be achieved with lower gain values, resulting in a smaller alteration of the equilibrium value (Fig. S20-C1 to C4).

**Figure 5:**
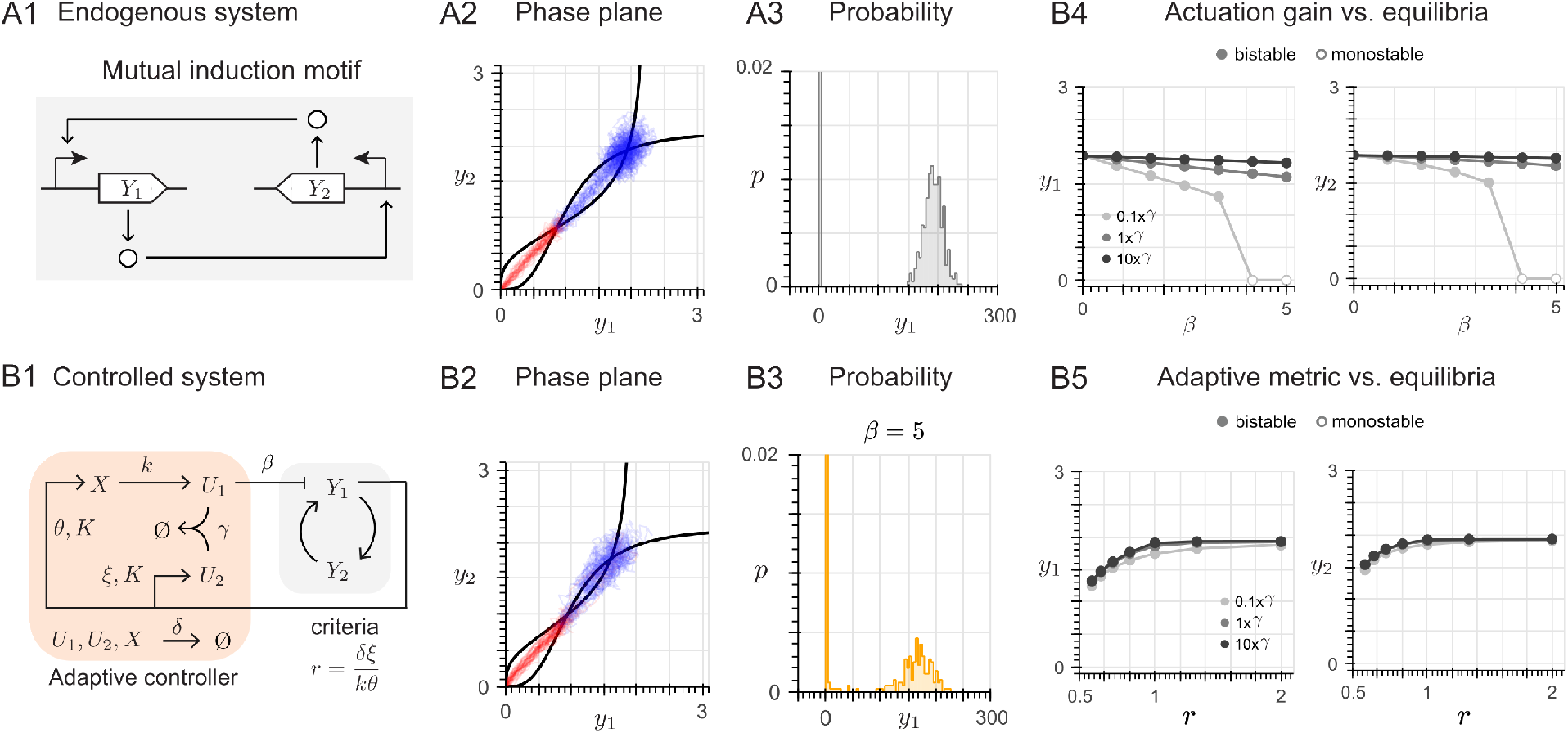
Applying the adaptive controller principle to a different system. (A1) Mutual activation motif, composed of two mutually inducing transcription factors Y_1_ and Y_2_. (A2) For the mutual activation motif, nullclines (black lines) and trajectories in the phase plane, converging either to the stable equilibrium with high expression of both Y_1_ and Y_2_ (blue) or to the equilibrium with zero expression (red). (A3) The trajectories in (A2) yield an asymmetric probability distribution for each species. The probability distribution for Y_2_ is the same as that for Y_1_. (B1) Architecture of the proposed adaptive controller with an inhibiting effect through species U_1_. (B2) For the controlled system, nullclines (black lines) and trajectories in the phase plane. (B3) Generation of a biased cell fate, which favors the equilibrium with no expression: the probability of converging to the equilibrium with high expression significantly decreases for both species. (B4) Characterization of the effect on the equilibrium value of increasing inhibition strength β, for different choices of the sequestration rate γ. (B5) Characterization of the effect on the equilibrium value for varying adaptive metric 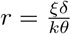 for different values of the sequestration rate γ

#### 2.3.2 Applying the adaptive control principle to multistable systems

To gain insight into the application of the adaptive controller to control and intermediate equilibrium point, we evaluated the steady-state behavior of the toggle switch with a self-activation motif in both species, whose architecture is depicted in Fig. 6-A1. In this case, in light of the high density of trajectories converging to the intermediate equilibrium and the corresponding peak in the probability distribution (Fig. 6-A2 and A3, respectively), we wish to design an strategy to reduce the probability of the system to converge to the intermediate equilibrium. To this aim, we implement an inhibition strategy using an adaptive controller for each species (Fig. 6-B1), based on the results obtained from the analysis of the mutual activation motif. Considering the same chemical reactions used to describe the kinetics of both the toggle switch with a double self-activation motif and the inhibitory strategy, the same equations as Eq. (18) and (19) can be considered, with *α*_1_ = *α*_2_ = *α*_3_ = *α* and a control input acting on each species, resulting in the ODEs

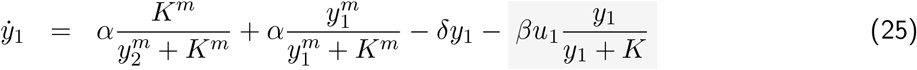

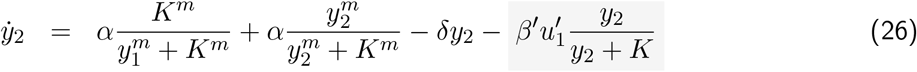

where the controller equations have a similar form as Eq. (7), (8) and (9). With a small alteration of the equilibrium value (Fig. 6-B2), the proposed control strategy reduces the probability of converging to the equilibrium where both species are equally produced, and induces a higher production of either *Y*_1_ or *Y*_2_ (Fig. 6-B3).

**Figure 6:**
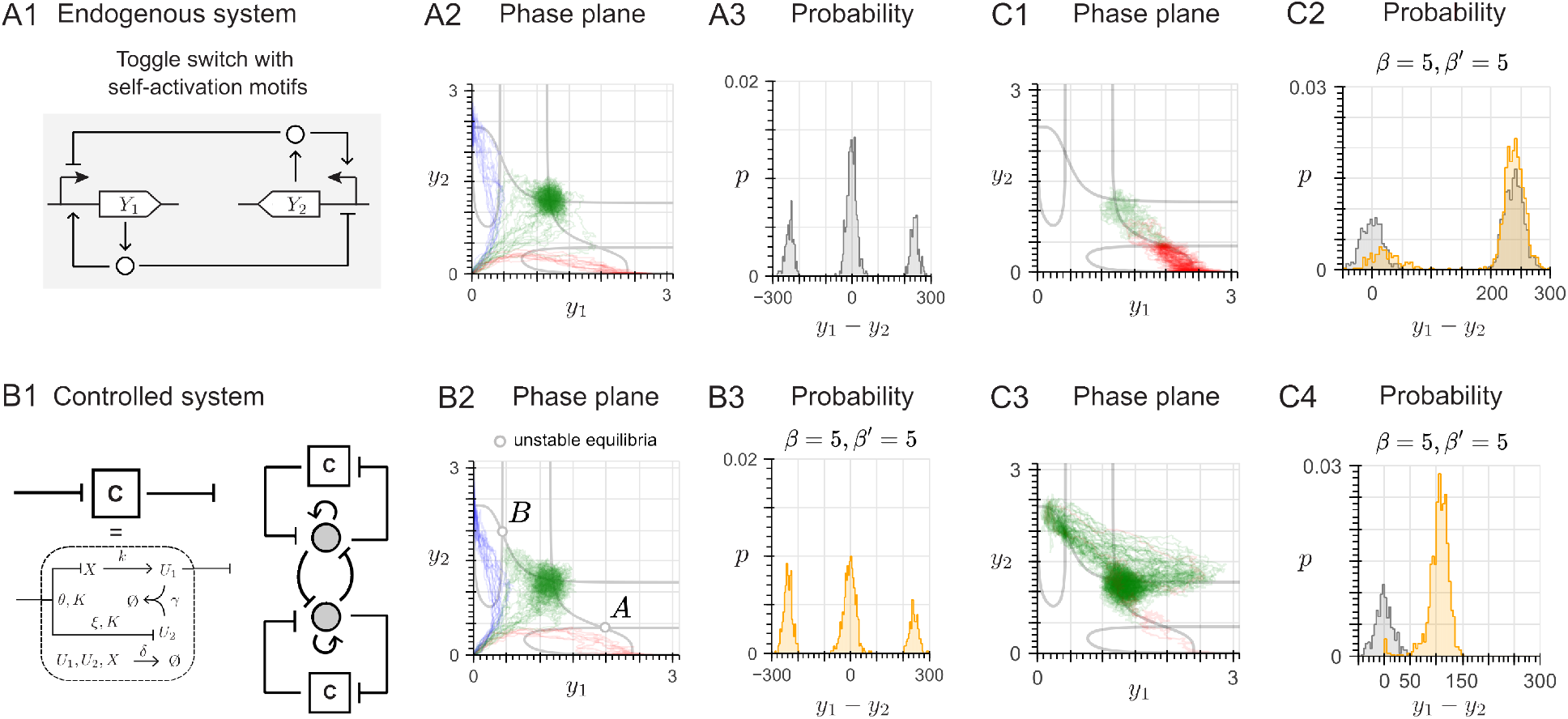
Controlling an intermediate equilibrium point. (A1) Architecture of the toggle switch with a self-activation loop acting on both species. (A2) For the toggle switch with a double self-activation, nullclines (black lines) and trajectories in the phase plane, exhibiting three stable equilibrium points: high expression of Y_2_ and low expression of Y_1_ (blue), low expression of Y_2_ and high expression of Y_1_ (red), and an intermediate state with the same level of expression for both Y_1_ and Y_2_ (green); and two unstable equilibrium points. (A3) The trajectories in (A2) yield a multimodal probability distribution, where convergence to the intermediate equilibrium is favored. (B1) Architecture of the proposed design strategy to control minimize the probability corresponding to the intermediate equilibrium of (A1): two adaptive controllers with the negative feedback architecture are implemented. (B2) For the system with both controllers, nullclines and trajectories in the phase plane. (B3) Generation of the desired biased cell fate: the probability of converging to the intermediate equilibrium is decreased. (C1) For the system with both controller, considering the Eqs.(27) and (28), and setting the initial conditions at the unstable equilibrium A, nullclines and phase plane (C2) For the same system as (C1), setting the initial conditions at the unstable equilibrium B, nullclines and phase plane (C3) The trajectories in (C1) generate the desired cell fate, favoring the equilibrium of high expression of Y_1_ and low expression of Y_2_ (red) over the intermediate equilibrium (green) (C4) The adaptive control strategy fails to generate the desired cell fate for the conditions of (C2)

Until now, we have considered the null expression of the species involved in the studied networks (*Y*_1_ and *Y*_2_) as our initial condition, as this enables the trajectories to converge to the equilibrium points with equal probability. Extending the analysis to other initial conditions (mainly, unstable equilibria), we defined the working limits of the adaptive controller. Particularly, to illustrate the conditions in which the controller fails to generate a biased cell fate, we set as our goal to design an strategy to favor the high production of the *Y*_1_ specie, associated with the equilibria associated with the high expression of *Y*_1_ and the low expression of *Y*_2_ (see Fig. 6–A2 and B2, in red). As for the control strategy, we consider a similar network as detailed in Fig. 6–B1, and consider a positive (inducing) effect over the *Y*_1_ specie rather than a negative (inhibitory) one. Hence, the controller will simultaneously increase the production of the target specie *Y*_1_ while decreasing the production of the non-target specie *Y*_2_. Considering the same chemical reactions used to describe the previous example and including a positive action as the one applied for the toggle switch with a single controller with the negative feedback architecture, we can modify Eqs. (25) and (26), resulting in the following ODEs:

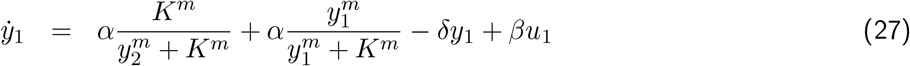

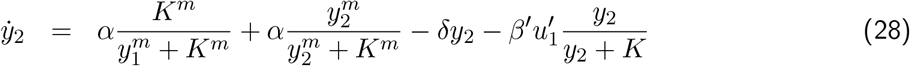

where the controller equations have a similar form as Eq. (7), (8) and (9). As shown in Fig. 6–C1, if we set the initial conditions in the unstable equilibrium named as “A” in the phase plane (see Fig. 6– B2), and apply the controller, we noticed that a biased cell fate is achievable, favoring the adoption of the equilibria associated with the high expression of *Y*_1_ and the low expression of *Y*_2_, rather than the intermediate equilibria. However, following the same control goal, if we set the initial conditions in the unstable equilibrium named “B” in the phase plane (see Fig. 6–B2), the control strategy fails to achieve the desired cell fate, even for increasing control gain values either inducing the expression of the target specie *Y*_1_ (by increasing *β*) or inhibiting the expression of the non-target specie *Y*_2_ (by increasing *β*^*′*^) (see Fig. S21).

## 3 Discussion

Synthetic biology offers promising tools to engineer cell fate, and improve the production of cell types that are phenotypically and functionally equivalent to their *in vivo* counterparts, thus enabling the development of novel cell-based therapeutics and disease models [9]. In this work, we propose a synthetic gene circuit based on an Incoherent Feedforward Loop motif and molecular sequestration to generate an output proportional to an approximated, discrete temporal derivative of its input. By enabling the controller to act on a target species involved in a decision-making process within a multistable gene regulatory network, we achieve a tunable synthetic bias in cell fate with minimal alteration of the equilibrium value.

Throughout this work, we outlined the parametric conditions that enable a biased cell fate through theoretical analysis, resulting in two principal design requirements: the circuit needs to operate in a fast sequestration regime (*γ*→ ∞ ) and satisfy the ideal adaptive metric *r* = *ξδ/kθ* = 1. When both conditions are met, our computational simulations demonstrate the generation of a synthetic bias, which can be further tuned through the control gain parameter (*β*). A limitation of the proposed design is the need for kinetic parameter fine-tuning to achieve the ideal adaptive metric during experimental implementation. Although biological differentiators have not yet been implemented experimentally, previous theoretical work regarding approximated differentiators and derivative motifs for biomolecular PID controllers (corresponding to an adaptive metric *r >* 1) already suggests a synthetic implementations [43, 44, 45, 46, 47]. Further advances in RNA technology and protein engineering offer even more choices to achieve the desired tunability [48, 49]. Overall, to assess the robustness with respect to variations in key parameters and metrics, we provide complementary simulations, which show that a relatively wide variation of the adaptive metric ( *±* 10% with respect to the equilibrium without the controller) does not alter the control performance.

Furthermore, we consider various design architectures built by varying both the feedback species – resulting in negative and positive feedback architectures – and the input species to the endogenous network, either through the *U*_1_ or *U*_2_ species. Our simulations demonstrate that all architectures can generate a synthetic bias with minimum alteration of the equilibrium value, and validate that the effectiveness of the adaptive controller is independent of the positive/negative feedback architecture and of the input to the controlled species. For control gains below a threshold value (*β* = 1 for the toggle switch case study), the performance is essentially the same regardless of the architecture. On the other hand, for control gains above that threshold, the behavior of the controller is architecture-specific. Negative feedback more strongly biases cell fate, but alters the equilibrium value more significantly. Hence, this architecture is well suited for increasing the yield of differentiation protocols and maximizing a desired population, such as *β*-like cells during stem cell-derived pancreatic islet manufacturing [10], or midbrain dopaminergic neurons [13, 12], while minimizing the number of cells of undesired, off-target types [15, 14]. On the other hand, positive feedback alters the equilibrium values less, but also induces less bias in the cell fate, making it better suited for subpopulation tuning; for example, balancing the cell ratio between *β* and *α* within an islet-like cluster [14], or adjusting the proportion between parenchymal and supporting cells, often associated with increased functionality and maturation [50, 51].

As endogenous regulatory networks grow in complexity (for instance, in terms of the number of genes involved in the decision-making process increases), quantify kinetic parameters for further mathematical modeling is harder. Hence, it is fundamental to address parameter uncertainty, due to parameter variations or even fluctuations over time [52]. When the system is subject to parameter variations (in particular, in the case of an unbalanced toggle switch), we show that increasing the value of the control gain in the positive feedback architecture confers robustness to changes in the kinetic parameters. However, only the equilibrium values corresponding to the controlled species can be maintained. An additional controller can be implemented to extend the range of parameter variation in which the multistable behavior (bistable in our case study) is maintained, but, even for higher gain values, we can only ensure minimal alteration of the equilibrium value for the species unaffected by the kinetic mutation. Conversely, the appearance of new, undesired equilibria can be avoided, compared to the addition of a self-activation motif.

In our last simulations, we outline general principles for applying the controller to different endogenous networks, demonstrating that both a positive (activating) and a negative (inhibiting) effect can be enforced over the endogenous network. Overall, by considering the mutual inhibition (Section 2.1.3) and mutual activation (Section 2.3.1) examples, we show that a synthetic bias can be generated by either favoring the production of the desired species (positive, activation-based control) or decreasing the production of the other species (negative, repression-based control). Moreover, we illustrate a preliminary strategy for translating the adaptive controller to higher-order multistable systems, considering the toggle switch with a self-activating motif on each specie as an example. Given the tristable behavior of this network, control over the probability of the intermediate equilibrium, associated with the equal expression of both species, was achieved by considering two adaptive controller, each inhibiting one specie. Based on these results, we conjecture that multistable system where intermediate equilibrium values are present will need one adaptive controller per specie associated with each equilibria. Finally, we identified the conditions were the adaptive controller fails to generate a biased cell fate. Within this context, considering that the transfer function shown in Eq. (4) resembles a proportional-derivative (PD) controller [34], future work will aim at implementing a strategy to generate an adaptive mechanism for the metric *r* = *ξδ/kθ* itself, yielding a two-step control strategy where the proportional component bypasses the basin of attraction of the intermediate equilibrium (e.g. based on the high-gain feedback controller proposed in [53]), and eventually shuts down (*r* = 1), leaving the derivative component working as described in this work.

A few studies have previously addressed cell fate control through the experimental implementation of synthetic gene circuits. Haellman et al. [54] optimized a vanillic-acid responsive induction system to enhance control over gene expression in human stem cells: the dose-response curve was fine-tuned into a switch, and applied to successfully commit progenitor cells to a pancreatic fate by regulating the expression of the TGIF2 and HHEX genes, but with a low yield (*<* 20% of the total population). In a more recent, proof-of-concept study, Prochazka et al. [55] designed a Boolean classifier able to discriminate between stem cells and differentiated cells, and actuate accordingly. The circuit was used to adjust the proportion between the three primordial germlines by controlling the expression of the endogenous morphogen BMP4; and through the action of the circuit in human pluripotent stem cells, the relative cell composition switch from a predominant ectoderm-like fate to relatively balanced subpopulations, including endoderm-like and mesoderm-like fates, although each represent *<* 30% and ∼ 10% of the total population, respectively. Moreover, both strategies implement an open-loop control, and hence require a pre-defined input (e.g., the optimal concentration of TGIF2 for commitment into a pancreatic fate [54], or the microRNA sequence that encodes the desired BMP4 secretion level [55]), whose knowledge is hard to achieve, also because concentrations vary in time. Galloway et al. [19] addressed this drawback by adding a feedback loop through a promoter responsive to the signaling pathway (MAPK) that controls the decision between mating and non-mating fate in yeast by combining positive and negative actions. Upon optimization, the controller was able to increase or decrease mating efficiency to a 70%. This last strategy resembles traditional controller design (see Fig. 2–A and [56]), which is still constrained by the knowledge of the dynamics of the reference species, which can be empirically determined, but could be difficult to extrapolate to higher organisms [19]. In this context, a key technical advantage of the proposed adaptive controller lies in the fact that it requires only partial knowledge of the species involved in the decision-making event, and it can thus exploit current computational algorithms that are not yet able to map the complete signaling pathways regulating cell fate but provide enough information for design [57, 58, 33].

An interesting avenue for future work on the control of multistable systems involves exploring switching between cell states (e.g, in Fig. 2-A2, switching from the “red” to the “blue” state) with minimum alteration of the equilibrium value, often referred to as “reprogramming”, with potential applications to stem cell-based therapeutics [59] and regenerative medicine [60]. Similarly, novel engineered circuits that exhibit multistability, such as the MultiFate-2 [61], can be implemented for *in-vitro* characterization, and further improvements to the present work can address the limitations when targeting intermediate equilibria in high-dimensional networks. Ultimately, our understanding of the non-local behavior and transient response of the adaptive controller can be enhanced by adopting more advanced mathematical methods to address the nonlinearities of the proposed chemical networks.

## 4 Methods

### 4.1 Numerical simulations

For deterministic simulations, the ODE models described in this work were integrated using Python’s *scipy* package with the parameters listed in Table 1. A comprehensive list of the models simulated, along with their corresponding nullcline equations, is available in the Supplementary Material. These were used to generate phase plane plots and systematically characterize deviations from the equilibrium, as detailed in each figure’s captions. When the nullclines did not have closed-form algebraic expressions, numerical methods were used to approximate them using the *scipy* and *numpy* packages. For stochastic simulations, the Gillespie algorithm was implemented through the *biocircuits* package. Probability distributions were computed as histograms generated from over 1000 trajectories for *t* = 200 h, with a bin number of 8.

**Table 1.** Nominal simulation parameters of each simulated endogenous regulatory network and adaptive controller, recovered from [34, 62, 38].

| Parameter | Description | Value |
| --- | --- | --- |
| Toggle switch and mutual activation motif |  |  |
| $\alpha$ ($\mu\text{M}/\text{h}$) | Production | 2.2 |
| $K$ ($\mu\text{M}$) | Hill activation constant | 1 |
| $m$ | Cooperativity | 3 |
| $\delta$ ($/\text{h}$) | Decay | 1 |
| Toggle switch with self-activation |  |  |
| $\alpha$ ($\mu\text{M}/\text{h}$) | Production | 1.2 |
| $K$ ($\mu\text{M}$) | Hill activation constant | 0.5 |
| $m$ | Cooperativity | 4 |
| $\delta$ ($/\text{h}$) | Decay | 1 |
| Adaptive controller |  |  |
| $\xi$ ($\mu\text{M}/\text{h}$) | Production | 1 |
| $\theta$ ($\mu\text{M}/\text{h}$) | Production | 1 |
| $k$ ($/\text{h}$) | Production | 1 |
| $\beta$ ($/\text{h}$) | Control gain | 1 |
| $\gamma$ ($/\mu\text{M}/\text{h}$) | Sequestration rate | 100 |

### 4.2 Data and code availability

All data presented in this article were generated exclusively using computational models, which can be found in our Github repository (https://github.com/frank-britto/engineering_cell_fate), along with the original code for generating both the figures in the main text and the Supplementary Figures.

## 5 Acknowledgments

We thank Jose Vargas, Enoch Antwi, Cindy Ren, Jérémie Marlhens and Rongrong Du for useful comments and discussions.

