## Supplementary_Figures_and_Text for "Engineering cell fate with adaptive feedback control"

### Engineering cell fate with adaptive feedback control (Supplementary Material)

November 29, 2024

**3 Robustness analysis under kinetic mutations** **15**

**4 Extending the applications of the adaptive controller** **26**

**5 Models** **28**

5.5.1 Single inhibiting controller in a negative feedback with actuation through $U_1$ specie 33

5.5.2 Single inhibiting controller in a positive feedback with actuation through $U_1$ specie 34

5.5.3 Single inhibiting controller in a negative feedback with actuation through $U_2$ specie 35

### 1 Analysis of the adaptive controller

The results obtained in the main text that demonstrate the dynamic properties of the proposed adaptive controller in a closed-loop system can be further complemented with theoretical analysis and numerical simulations. In this section, we will address the following properties:

1. The controller introduces minimum perturbation to the equilibrium values in the high sequestration regimen ($\gamma \to \infty$), evaluated through the analysis of the nullclines equations corresponding to the toggle switch (mutual inhibition)

2. The ideal adaptive metric ($r = 1$) admits an error margin where the dynamic properties are maintained

#### 1.1 Approximated dynamics in the fast sequestration regime

In previous work, we analyzed the combination of an Incoherent FeedForward Loop (IFFL) and a negative feedback network (NF) to generate a band-pass filter with tunable cut-off frequencies [1]. For our current analysis, we followed the same analytical approach for evaluating the dynamics of the proposed controller. Starting with the chemical reactions that describe the open-loop system, for an arbitrary input specie $Y$,

| | | |
|---|---|---|
| $Y \xrightarrow{\theta} Y + X$ | $X \xrightarrow{\delta} \emptyset$ | Production/Decay |
| $Y \xrightarrow{\xi} Y + U_2$ | $U_2 \xrightarrow{\delta} \emptyset$ | Production/Decay |
| $X \xrightarrow{k} X + U_1$ | $U_1 \xrightarrow{\delta} \emptyset$ | Production/Decay |
| $U_1 + U_2 \xrightarrow{\gamma} C$ | $C \xrightarrow{\delta} \emptyset$ | Sequestration/Decay |

Note that we are considering the specie $C$ that corresponds to the complex formation between the $U_1$ and $U_2$ species as a result of the sequestration reaction, which also decays at rate $\delta$. Under the law of mass action, we can modeled these chemical reactions using Ordinary Differential Equations (ODEs). Following the coordinate transformation described in [1], so that $u_1^T(t) = u_1(t) + c(t)$ and $u_2^T(t) = u_2(t) + c(t)$, we can avoid the nonlinearities and describe the controller´s dynamics as follows:

$$
\begin{aligned}
\dot{u_1}^T &= kx(t) - \delta u_1^T(t) \\
\dot{u_2}^T &= \xi y(t) - \delta u_2^T(t) \\
\dot{x} &= \theta y(t) - \delta x(t) \\
\frac{1}{\gamma}\dot{c} &= [u_1^T(t) - c(t)][u_2^T(t) - c(t)] - \frac{\delta}{\gamma}c(t)
\end{aligned}
$$

Applying the timescale separation argument [2], it can be shown that, in the fast sequestration regime ($\gamma \to \infty$), the asymptotic value for $u_1(t)$ can be computed as

$$\lim_{\gamma \to \infty} \bar{u_1} = \max\{0, \bar{u_1}^T - \bar{u_2}^T\} \quad (1)$$

Furthermore, since $\dot{u_1}^T$, $\dot{u_2}^T$ and $\dot{x}$ are described by linear ODEs, we can take the Laplace transform.

$$X(s) = \frac{\theta}{s+\delta} Y(s) \tag{2}$$

$$U_1^T(s) - U_2^T(s) = \frac{k}{s+\delta} X(s) - \frac{\xi}{s+\delta} Y(s) \tag{3}$$

Replacing Eq. (2) in Eq. (3), we obtain the following expression for the difference between $U_1^T(s)$ and $U_2^T(s)$ in the Laplace domain,

$$U_1^T(s) - U_2^T(s) = \frac{k\theta - \xi\delta - \xi s}{s^2 + 2\delta s + \delta^2} = \frac{k\theta}{(s+\delta)^2}[1 - \frac{\xi\delta}{k\theta} - \frac{\xi}{k\theta} s] Y(s) \tag{4}$$

Finally, by taking the inverse Laplace transform of Eq.(4) and considering Eq.(1), we obtained the expression shown in the main text.

$$u_1(t) \approx \max\left\{0, \mathcal{L}^{-1}\left(\frac{k\theta}{(s+\delta)^2}\left[1 - \frac{\xi\delta}{k\theta} - \frac{\xi}{k\theta} s\right] Y(s)\right)\right\} \tag{5}$$

Retrieving the adaptive metric from the main text: $r = \xi\delta/k\theta$ and setting $r = 1$, we obtained

$$u_1(t) \approx \max\left\{0, \mathcal{L}^{-1}\left(\underbrace{\frac{k\theta}{(s+\delta)^2}}_{Low-pass filter} \underbrace{\left[-\frac{\xi}{k\theta} s\right]}_{Derivative} Y(s)\right)\right\} \tag{6}$$

#### 1.2 The adaptive controller barely perturbs the equilibrium in the fast sequestration regime

##### 1.2.1 Non-dimensionalization of the ODEs

We define new variables to obtain a non-dimensionalized representation of the closed-loop system, considering the toggle switch´s study case. Let $\tau = \delta t, \hat{y}_1 = \frac{y_1}{K}, \hat{y}_2 = \frac{y_2}{K}, \hat{u}_1 = \frac{u_1}{K}, \hat{u}_2 = \frac{u_2}{K}, \hat{x} = \frac{x}{K}$ and parameters $\hat{\alpha} = \frac{\alpha}{\delta K}, \hat{\beta} = \frac{\beta}{\delta}, \hat{k} = \frac{k}{\delta}.\hat{\gamma} = \frac{\gamma K}{\delta}, \hat{\xi} = \frac{\xi}{\delta K}, \hat{\theta} = \frac{\theta}{\delta K}$, resulting in the following ODEs:

$$\frac{d}{d\tau}\hat{y}_1 = \hat{\alpha}\frac{1}{1+\hat{y}_2^m} - \hat{y}_1 + \hat{\beta}\hat{u}_1 \tag{7}$$

$$\frac{d}{d\tau}\hat{y}_2 = \hat{\alpha}\frac{1}{1+\hat{y}_1^m} - \hat{y}_2 \tag{8}$$

$$\frac{d}{d\tau}\hat{u}_1 = \hat{k}\hat{x} - \hat{u}_1 - \hat{\gamma}\hat{u}_1\hat{u}_2 \tag{9}$$

$$\frac{d}{d\tau}\hat{u}_2 = \hat{\xi}\frac{1}{1+\hat{y}_1^m} - \hat{u}_2 - \hat{\gamma}\hat{u}_1\hat{u}_2 \tag{10}$$

$$\frac{d}{d\tau}\hat{x} = \hat{\theta}\frac{1}{1+\hat{y}_1^m} - \hat{x} \tag{11}$$

Note that we can rewrite the approximated dynamics of $U_1$, described in Eq. (5), considering the non-dimensionalized ODEs. Therefore, we can retrieve as well the adaptive metric $r = \frac{\hat{\xi}}{\hat{k}\hat{\theta}}$.

##### 1.2.2 Boundedness analysis

Following a similar approach as [1], we can guarantee boundness of the state variables $\hat{y}_1$, $\hat{y}_2$, $\hat{u}_1$, $\hat{u}_2$ and $\hat{x}$ since

$$\begin{aligned}
\frac{d}{d\tau}\hat{x} &< \hat{\theta} - \hat{x} && (12)\\
\frac{d}{d\tau}\hat{u}_2 &< \hat{\xi} - \hat{u}_2 && (13)\\
\frac{d}{d\tau}\hat{u}_1 &< \hat{k}\hat{\theta} - \hat{u}_1 && (14)\\
\frac{d}{d\tau}\hat{y}_2 &< \hat{\alpha} - \hat{y}_2 && (15)\\
\frac{d}{d\tau}\hat{y}_1 &< \hat{\alpha} - \hat{y}_1 + \hat{\beta}\hat{k}\hat{\theta} && (16)
\end{aligned}$$

##### 1.2.3 Analysis of the controller´s perturbation to the equilibrium in the fast sequestration regime

We can find the nullclines equations for the $Y_1$ and $Y_2$ repressors by equationg $d\hat{y}_1/d\tau = 0$ and $d\hat{y}_2/d\tau = 0$ from Eqs. (7) and (8). As a result, we obtain

$$\begin{aligned}
\bar{y}_1 &= \hat{\alpha}\frac{1}{1+\hat{y}_2^m} + \hat{\beta}\bar{u}_1\\
\bar{y}_2 &= \hat{\alpha}\frac{1}{1+\hat{y}_1^m}
\end{aligned}$$

When $\hat{\beta} = 0$, it´s known that the system admits either one or three steady–state equilibrium points [3]. Since $\hat{y}_2 = f(\hat{y}_1)$, we can compute the steady–state equilibrium points by finding the real roots of the following 5-th order polynomial, constructed by replacing the nullclines equation of $\hat{y}_2$ in the expression for $\hat{y}_1$ and considering $m = 2$:

$$P_p(\hat{y}_1) = a_5\hat{y}_1^5 + a_4\hat{y}_1^4 + a_3\hat{y}_1^3 + a_2\hat{y}_1^2 + a_1\hat{y}_1 + a_0 = 0$$

where $a_5 = 1, a_4 = -\hat{\alpha}$, $a_3 = 2$, $a_2 = -2\hat{\alpha}$, $a_1 = 1 + \hat{\alpha}^2$ and $a_0 = \hat{\alpha}$. Then, to analyze the effect of the adaptive controller´s actuation (through $\hat{u}_1$) in the steady–state equilibrium of the toggle switch, we can study how the coefficients of the aforementioned polynomial change when $\hat{\beta} \neq 0$. Hence,

$$P(\bar{y}_1) = P_p(\bar{y}_1) \underbrace{-\hat{\beta}\hat{u}_1(\bar{y}_1^4 + 2\bar{y}_1^2 + 1 + \hat{\alpha}^2)}_{actuation} = 0 \qquad (17)$$

We can further find $\bar{u}_1$ as a function of $\bar{y}_1$ by equating $d\hat{u}_1/d\tau = 0$, $d\hat{u}_2/d\tau = 0$ and $d\hat{x}/d\tau = 0$.

$$\begin{aligned}
\bar{u}_2 &= \frac{k\bar{x} - \bar{u}_1}{\hat{\gamma}\bar{u}_1} = \frac{\hat{\xi}/(1+\hat{y}_1^m)}{\hat{\gamma}\bar{u}_1 + 1}\\
\bar{x} &= \hat{\theta}\frac{1}{1+\bar{y}_1^m}
\end{aligned}$$

Let $f_1 = \hat{k}\bar{x}$ and $f_2 = \hat{\xi}\frac{1}{1+\bar{y}_1^m}$. We can write the nullclines equation of $\bar{u}_1$ as a second order polynomial given by

$$\bar{u}_1^2 + \bar{u}_1(f_2 - f_1 + \frac{1}{\hat{\gamma}}) - \frac{f_1}{\hat{\gamma}} = 0 \tag{18}$$

Since molecule´s concentrations can´t be negative, the only feasible solution will be

$$\bar{u}_1 = \frac{1}{2}(f_1 - f_2 - \frac{1}{\hat{\gamma}} + \sqrt{(f_2 - f_1 + \frac{1}{\hat{\gamma}})^2 + \frac{4f_1}{\hat{\gamma}}}) \tag{19}$$

Now, recalling the adaptive metric $r = \hat{\xi}/\hat{k}\hat{\theta}$, by subtracting $f_1 - f_2 = \hat{k}\theta(1-r)\bar{y}^*$, where $y^* = \frac{1}{1+\bar{y}_1^m}$, we can rewrite Eq.(19) as follows:

$$\bar{u}_1 = \frac{1}{2}(y^*(1-r)\hat{k}\hat{\theta} - \frac{1}{\hat{\gamma}} + \sqrt{(-y^*(1-r)\hat{k}\hat{\theta} + \frac{1}{\hat{\gamma}})^2 + \frac{4f_1}{\hat{\gamma}}} \tag{20}$$

Then, we can find the limit of the asymptotic value of $\bar{u}_1$ in the vicinity of the ideal adaptive metric ($r = 1$), which can be computed as

$$\lim_{\hat{r}\to 1} \bar{u}_1 = \frac{1}{2}\left(-\frac{1}{\hat{\gamma}} + \sqrt{(\frac{1}{\hat{\gamma}})^2 + \frac{4f_1}{\hat{\gamma}}}\right) \leq \sqrt{\frac{f_1}{\hat{\gamma}}} \tag{21}$$

From Eq. (12), we notice that

$$\lim_{\hat{r}\to 1} \bar{u}_1 = \frac{1}{2}\left(-\frac{1}{\hat{\gamma}} + \sqrt{(\frac{1}{\hat{\gamma}})^2 + \frac{4f_1}{\hat{\gamma}}}\right) \leq \sqrt{\frac{f_1}{\hat{\gamma}}} \leq \sqrt{\frac{\hat{k}\hat{\theta}}{\hat{\gamma}}} \tag{22}$$

Considering this last upper bound, which describes the maximum value admissible for $\hat{u}_1$, we analyze the polynomial describing the nullclines equation of $\hat{y}_1$ in Eq. (17),

$$P(\bar{y}_1) = P_p(\bar{y}_1) - \hat{\beta}\sqrt{\frac{\hat{k}\hat{\theta}}{\hat{\gamma}}}[\hat{y}_1^4 + \hat{y}_1^2 + (1 + \hat{\alpha}^2)]$$

We notice that in the high sequestration regime, the asymptotic value of $P(\bar{y}_1)$ approaches $P_p(\bar{y}_1)$ since the effect of the controller´s actuation on the coefficients of the polynomial approach 0 as $\hat{\gamma} \to \infty$. Hence, the roots of $P(\hat{y}_1)$ will be closely similar to the one of $P_p(\hat{y}_1)$ since the contribution of the controller´s actuation is negligible.

### 2 Design guidelines for the adaptive controller

#### 2.1 The adaptive controller barely perturbs the equilibrium

We can crosscheck numerically the statements made for stating minimum perturbation, from the adaptive controller to the equilibria of the toggle switch from the analysis of the open-loop system, by computing the dynamics of $\hat{u}_1(\tau)$ obtained by solving Eqs. (7) to (11). For increasing values of sequestration rate

($\hat{\gamma}$), Fig. 1–A shows how the steady–state response of the adaptive controller approaches to zero, in a similar fashion as the dynamics of $u_1(t)$ shown in Fig. 2–C of the main text. Nevertheless, since setting the sequestration rate to a very high value is difficult experimentally, then $\hat{u}_1(\tau)$ will converge to a small, non-zero value that perturbs the equilibrium (although minimally, as already proven). Fig. 1–B characterizes the normalized error between the equilibrium point of the toggle switch in isolation and when the adaptive controller actuates ($\Delta y_1$, same for the $Y_2$ specie, without loss of generality in the analysis), as a function of the sequestration rate. For each value, we notice how the actuation gain ($\hat{\beta}$) determines the maximum deviation admitted to the equilibria. We can further determine empirically the stability of the closed-loop system as a function of both $\hat{\gamma}$ and $\hat{\beta}$, as shown in Fig. 1–C. From a designer perspective, once the high sequestration regime is achieved, the modulation of the actuation gain has to be within a range such that the bistability is preserved. Fig. 3 illustrates this conclusion with a concrete example, too.

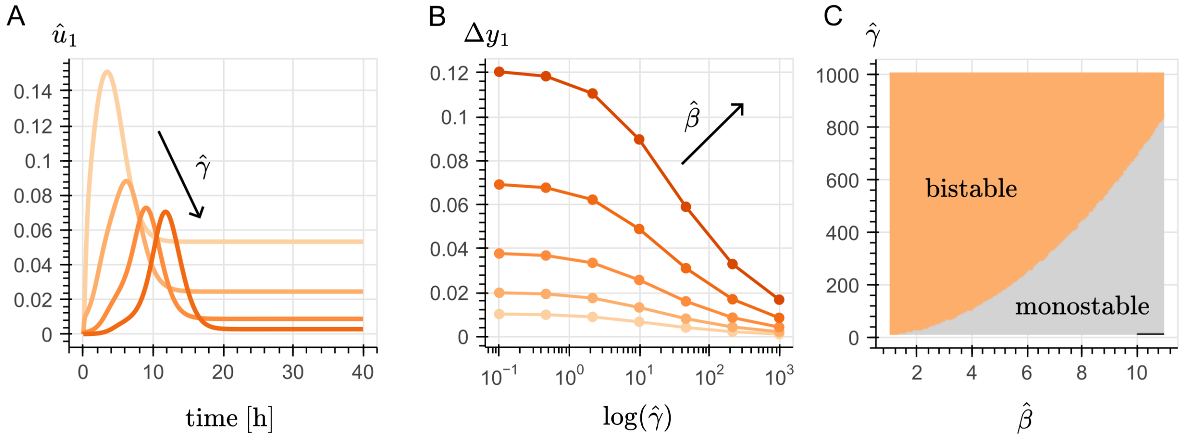

***Figure 1: The adaptive controller barely perturbs the steady–state equilibrium.*** *(A) In the closed loop system described by Eqs. (7)–(11), by increasing values of sequestration rate ($\hat{\gamma}$, as darker orange colors), the steady–state value of $\hat{u}_1(\tau)$ approaches zero. (B) Let $\Delta y_1$ be the difference between the equilibrium point associated with the high expression of $Y_1$ and low expression of $Y_2$ calculated from the toggle switch in isolation (Eqs. (7) and (11) for $\hat{\beta} = 0$), and the same equilibrium point, but calculated considering the actuation of the adaptive controller ($\hat{\beta} \neq 0$); divided by the former. (C) The relationship between $\hat{\beta}$ and $\hat{\gamma}$ suggested by Panel B is explored in terms of stability, by numerically determining the number of intersections between the nullclines equations derived by equating Eqs. 7–(11) to zero. Bistable is defined as three intersections between the nullclines curves, corresponding to one unstable equilibrium point, and two stable equilibrium points; while monostable is defined as a single intersection between the nullclines curves, corresponding to one stable equilibrium point.*

#### 2.2 Analyzing deviations from the adaptive metric

With a similar approach as the analysis presented in Section 1.1, we can determine an expression for the approximated dynamics of the adaptive controller in isolation, as the one provided in Eq. (6), using the non-dimensionalized model described in Eqs. (7)–(11). Considering the adaptive metric $r = \hat{\xi}/\hat{k}\hat{\theta}$, we can rewrite the expression as follows:

$$\hat{u}_1(\tau) \approx \max\left\{0, \mathcal{L}^{-1}\left(\frac{\hat{k}\hat{\theta}}{(s+1)^2}\left[1 - r - \frac{\hat{\xi}}{\hat{k}\hat{\theta}}s\right]Y(s)\right)\right\} \tag{23}$$

For a fixed gain value $\hat{\beta}$, the capacity of generating a biased output of the adaptive controller depends on the value of actuation specie $\hat{U}_1$. According to Eq. (23), for $r = 1$, that is true when the gradient is negative, since there is a negative sign before to the $s$ operator. A deviation from the metric introduces an offset, as depicted in Fig. 2–A. When $r > 1$, a greater gradient is needed for $u_1(t) > 0$. Thus, for the same range of values for the input $Y(s)$, $u_1(t)$ will take smaller values with respect to when $r = 1$, so the probability of a biased output will decrease. Inversely, when $r < 1$, a lesser gradient is needed for $u_1(t) > 0$. Similarly, for the same input, $u_1(t)$ will result in greater values with respect to when $r = 1$, so an increased probability of a biased output is expected.

As a reference, Fig. 2–B shows the input/output map of the adaptive controller in isolation, satisfying the ideal adaptive metric ($r = 1$). For comparison, Fig. 2–C replicates the same input/output map in the fast sequestration regime, varying the $r$ metric within a $\pm 20\%$ margin of its ideal value. As mentioned before, taking the magnitude of $\hat{u}_1(\tau)$ as a baseline, achieving $r < 1$ results in a higher magnitude, while $r > 1$ results in a reduced magnitude (see Fig. 4 for an illustrative example).

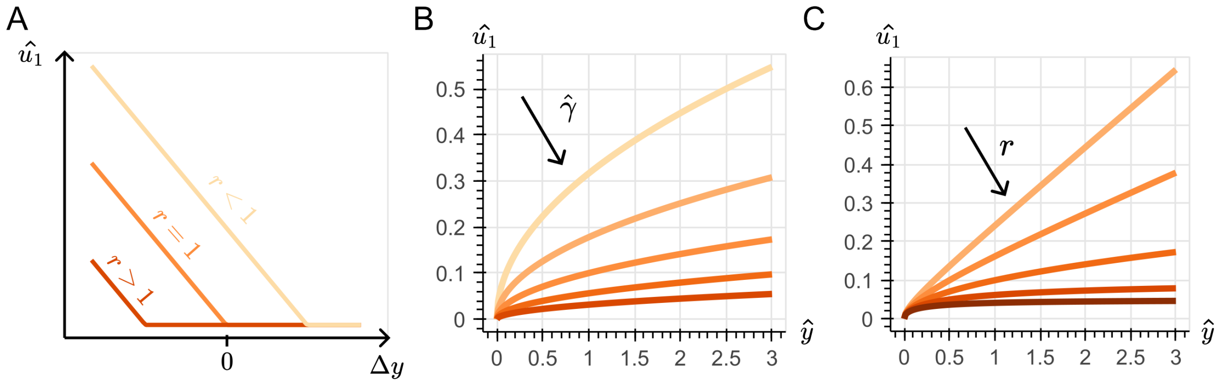

***Figure 2: Deviations from the ideal adaptive metric.*** *(A) With the ideal adaptive metric ($r = 1$), the adaptive controller responds to negative gradients ($-\frac{\hat{\xi}}{\hat{k}\hat{\theta}}s$). Any deviations to this metric result in an offset. (B) Input/output map of the open-loop system, highlighting the magnitude of $\hat{u}_1$ as a function of sequestration rate ($\hat{\gamma}$) (C) Input/output map for a fixed, high sequestration rate ($\hat{\gamma} = 1000$), and varying $\hat{\xi}$ within a $20\%$ of its nominal value, resulting in the metric between 0.8 and 1.2 times its nominal value 1.*

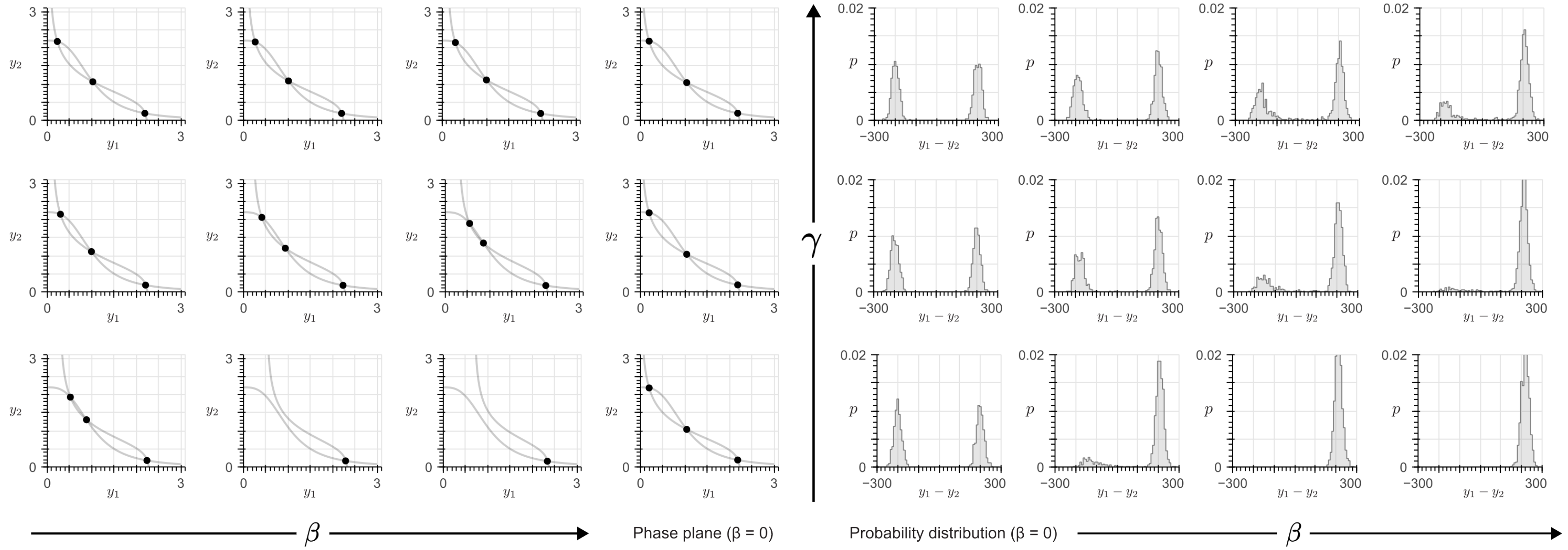

***Figure 3: Trade–off between the sequestration rate ($\gamma$) and the control gain ($\beta$)***. *From left to right, increasing $\beta$ corresponds to a larger deviation to the location of the equilibrium points, as well as an increase in the probability distribution favoring the production of the $Y_1$ species. A sufficiently high gain value ($\beta \geq 4$ for this specific example) results in a monostable system (i.e, single, stable equilibrium point, and a corresponding single "peak" in the probability distribution). From bottom to top, increasing the value of sequestration rate ($\gamma$) decreases the perturbation to the equilibrium, at the expense of the biased output, yielding a more symmetric, bimodal probability distribution, akin the one corresponding to the isolated toggle switch. Simulations were done by numerically solving Eqs.* (24)–(28) *for the nominal values indicated in Table 1. From left to right, $\beta$ corresponds to $1X$, $2X$ and $3X$ the nominal value of $\beta = 1$. From bottom to top, $\gamma$ corresponds to $0.1X$, $1X$ and $10X$ the nominal value of $\gamma = 100$.*

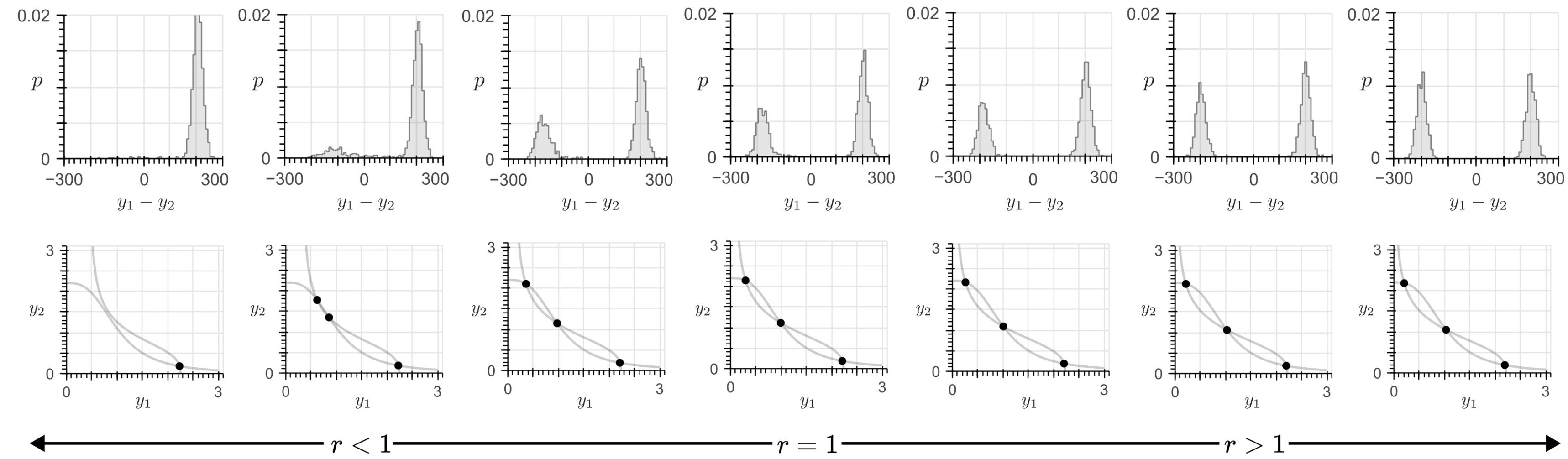

***Figure 4:* *The controller must satisfy the adaptive metric requirement (*$r = \xi\delta/k\theta$*).*** *A positive deviation (*$r > 1$*) decreases the probability of the biased cell fate, but minimizes the perturbation to the equilibrium. Inversely, a negative deviation (*$r < 1$*) increases the probability of the biased output, but with a corresponding higher deviation to the equilibrium. Simulations were done by numerically solving Eqs.* (24)–(28) *for the nominal values indicated in Table 1. The adaptive metric was varied within a* $20\%$ *of its nominal value of* $r = 1$ *by either increasing the value of the production rate* $k$ *(see Eq.* (27)*, yielding* $r < 1$*, or decreasing it, yielding* $r > 1$.

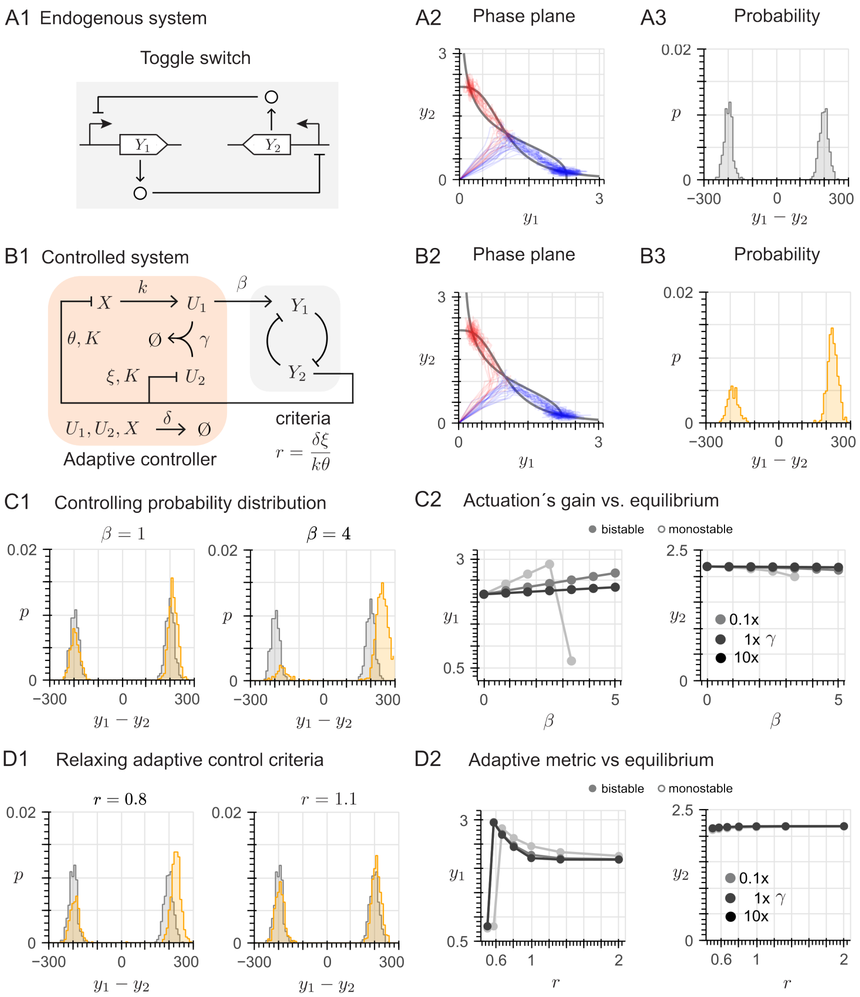

***Figure 5: The adaptive controller with a positive feedback loop enables a biased output.*** *From the main text, we include the following results for comparison: (A1) Architecture of the toggle switch, formed by two mutually inhibiting transcription factors* $Y_1$ *and* $Y_2$*. (A2) For the toggle switch, in the phase plane, the nullclines (black lines) are shown along with* $1000$ *trajectories starting from different initial conditions; each trajectory converges either to the stable equilibrium point with high expression of* $Y_1$ *and low expression of* $Y_2$ *(blue trajectories) or to the stable equilibrium point with high expression of* $Y_2$ *and low expression of* $Y_1$ *(red trajectories). (A3) The trajectories in (A2) converge essentially with equal probability to either of the stable equilibria, resulting in an unbiased bimodal probability distribution. (B1) Architecture of the toggle switch with the adaptive controller* ***with a positive feedback architecture****. (B2) For the controlled system, in the phase plane, the nullclines (black lines) are shown along with* $1000$ *trajectories starting from different initial conditions; more trajectories converge to the stable equilibrium point with high expression of* $Y_1$ *(blue) and less to the stable equilibrium point with high expression of* $Y_2$ *(red). (B3) The controller yields a biased cell fate: the trajectories in (B2) converge with higher probability to the equilibrium where the production of* $Y_1$ *is favored over that of* $Y_2$*. (C1) Increasing the control gain* $\beta$ *increases the bias in the cell fate, leading to a larger imbalance in the probability distribution. (C2) Equilibrium values for increasing control gain* $\beta$*, for different values of the sequestration rate* $\gamma$*. (D1) Effect of a* $10\%$ *increase and a* $20\%$ *decrease of the adaptive metric* $r = \frac{\xi\delta}{k\theta}$ *from its nominal value* $1$*: a biased cell fate is still generated at the price of a small alteration of the equilibrium values. (D2) Equilibrium values for varying adaptive metric* $r = \frac{\xi\delta}{k\theta}$*, for different values of the sequestration rate* $\gamma$*.*

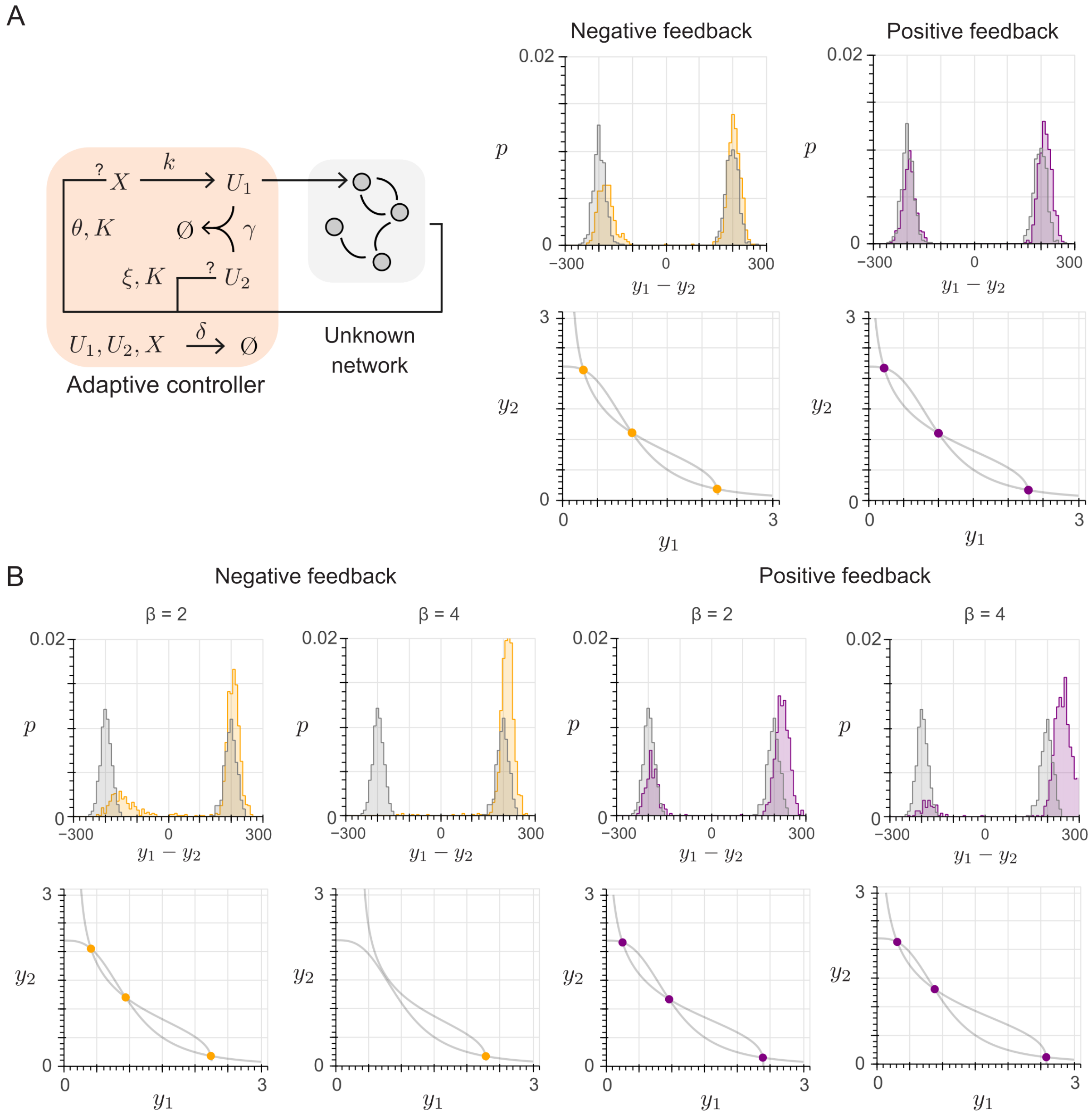

***Figure 6: The adaptive controller is effective in both negative and positive feedback architectures.*** *(A) By satisfying the high sequestration rate and ideal adaptive metric requirements, we numerically validate that the effectiveness of the proposed adaptive controller is independent of the type of feedback provided by the endogenous gene regulatory network. From low control gain values ($\beta \leq 1$, for the toggle switch example), the performance of the adaptive controller (i.e, a biased cell fate and the minimal equilibrium alteration) is similar for both architectures. (B) For higher control gain values ($\beta > 1$ for the toggle switch example), the negative feedback architecture enables a higher probability for the production of the target species $Y_1$, at the price of a higher perturbation to the equilibrium. On the other hand, the positive feedback architecture minimizes the deviation to the equilibrium, but requires a higher $\beta$ value to achieve the same probability distribution as the negative feedback architecture.*

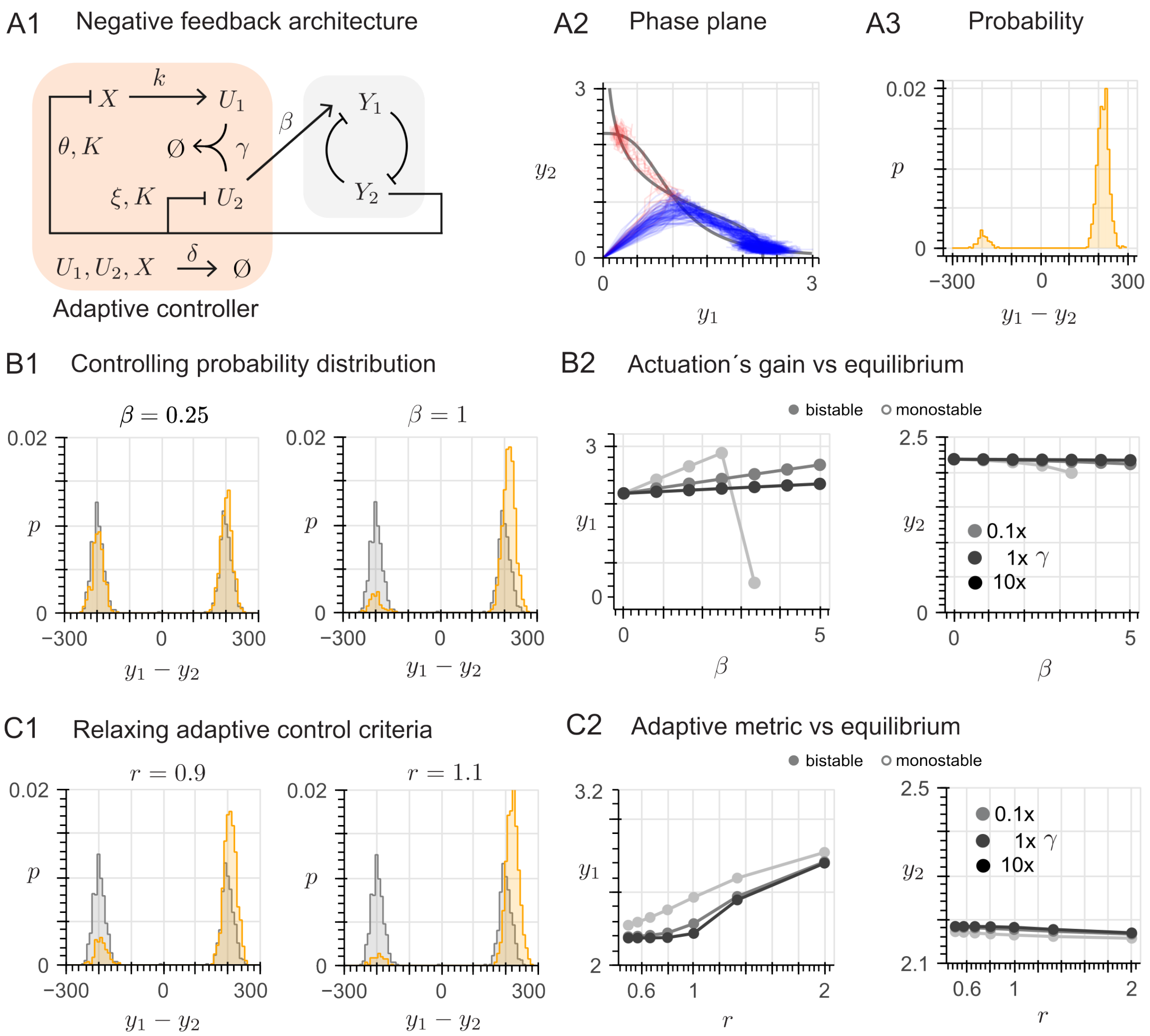

***Figure 7: The adaptive controller is symmetric with respect to its actuation.*** *(A1) Architecture of the toggle switch with the adaptive controller, where* $Y_2$ *is the feedback species, and the controller's actuation is enforced via the* $U_2$ *species; overall corresponding to a negative feedback architecture. (A2) For the controlled system, in the phase plane, the nullclines (black lines) are shown along with* $1000$ *trajectories starting from different initial conditions; more trajectories converge to the stable equilibrium point with high expression of* $Y_1$ *(blue) and less to the stable equilibrium point with high expression of* $Y_2$ *(red). (A3) The controller yields a biased cell fate: the trajectories in (A2) converge with higher probability to the equilibrium where the production of* $Y_1$ *is favored over that of* $Y_2$*. (B1) Increasing the control gain* $\beta$ *increases the bias in the cell fate, leading to a larger imbalance in the probability distribution. Note that a higher* $\beta$ *value is needed to achieve a similar probability distribution, compared to the negative feedback architecture. (B2) Equilibrium values for increasing control gain* $\beta$*, for different values of the sequestration rate* $\gamma$*. (C1) Effect of a* $10\%$ *deviation of the adaptive metric* $r = \frac{\xi\delta}{k\theta}$ *from its nominal value* $1$*: a biased cell fate is still generated at the price of a small alteration of the equilibrium values. (C2) Equilibrium values for varying adaptive metric* $r = \frac{\xi\delta}{k\theta}$*, for different values of the sequestration rate* $\gamma$*.*

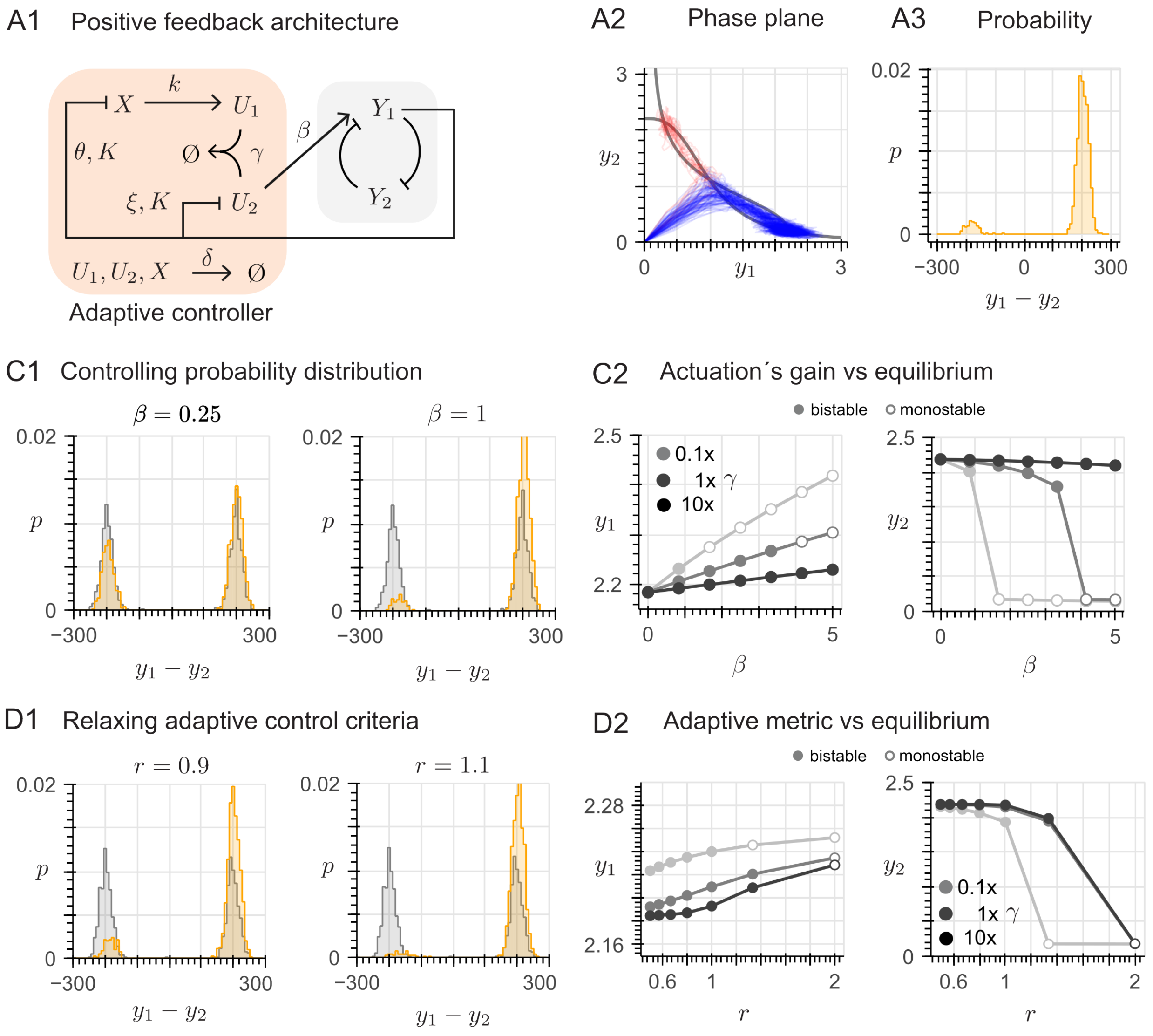

***Figure 8: The adaptive controller is symmetric with respect to its actuation.*** *(A1) Architecture of the toggle switch with the adaptive controller, where $Y_1$ is the feedback species, and the controller's actuation is enforced via the $U_2$ species; overall corresponding to a positive feedback architecture. (A2) For the controlled system, in the phase plane, the nullclines (black lines) are shown along with $1000$ trajectories starting from different initial conditions; more trajectories converge to the stable equilibrium point with high expression of $Y_1$ (blue) and less to the stable equilibrium point with high expression of $Y_2$ (red). (A3) The controller yields a biased cell fate: the trajectories in (A2) converge with higher probability to the equilibrium where the production of $Y_1$ is favored over that of $Y_2$. (B1) Increasing the control gain $\beta$ increases the bias in the cell fate, leading to a larger imbalance in the probability distribution. Note that a higher $\beta$ value is needed to achieve a similar probability distribution, compared to the negative feedback architecture. (B2) Equilibrium values for increasing control gain $\beta$, for different values of the sequestration rate $\gamma$. (C1) Effect of a $10\%$ deviation of the adaptive metric $r = \frac{\xi\delta}{k\theta}$ from its nominal value $1$: a biased cell fate is still generated at the price of a small alteration of the equilibrium values. (C2) Equilibrium values for varying adaptive metric $r = \frac{\xi\delta}{k\theta}$, for different values of the sequestration rate $\gamma$.*

### 3 Robustness analysis under kinetic mutations

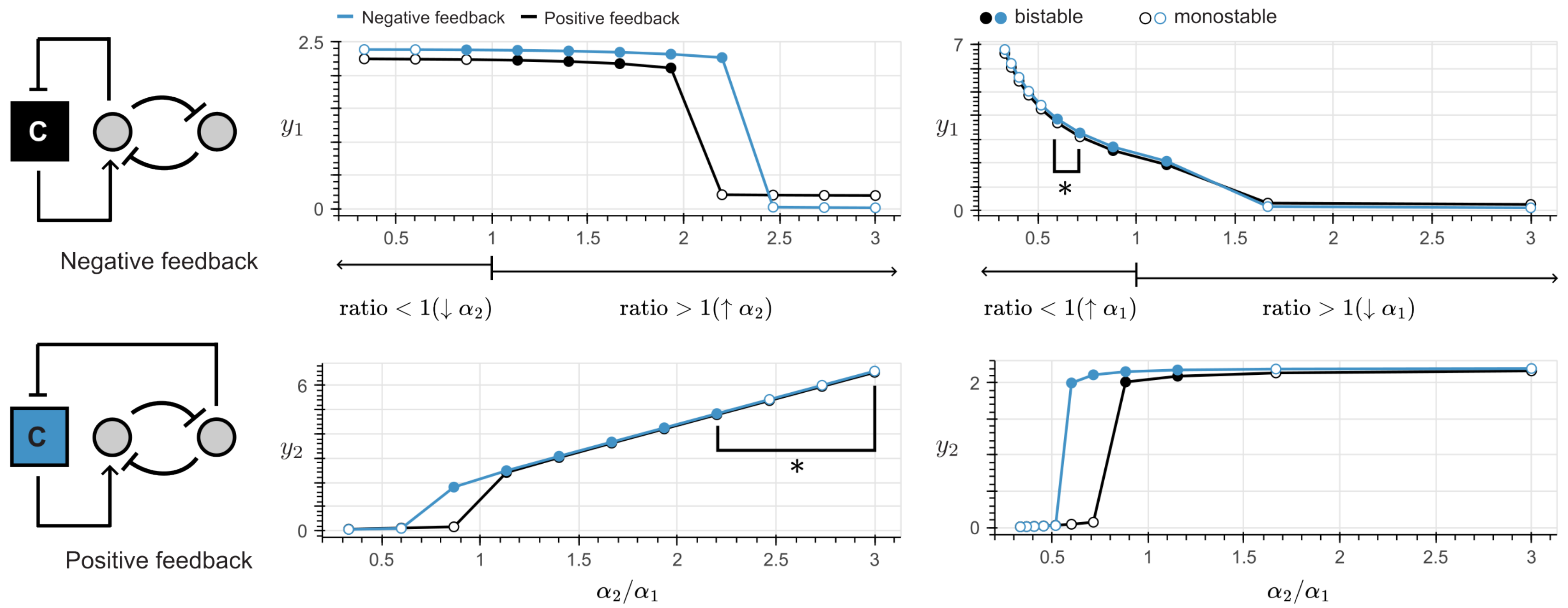

***Figure 9: The positive feedback configuration (blue line) is more robust than the negative feedback one (black line)**. For small values of control gain ($\beta \leq 1$), both configurations exhibit similar dynamic properties (Fig. S6–A), so their robustness to parametric perturbations is similar (not shown here). For higher values of actuation gain (for this example, $\beta = 2$), the positive feedback exhibits a wider range of $\alpha_2/\alpha_1$ values for which the bistable behavior is maintained. For the top right and bottom left panels, the points highlighted with the (*) symbol correspond to monostability of the black line, overlapped by the blue line, since deviations to the equilibrium are within a similar magnitude.*

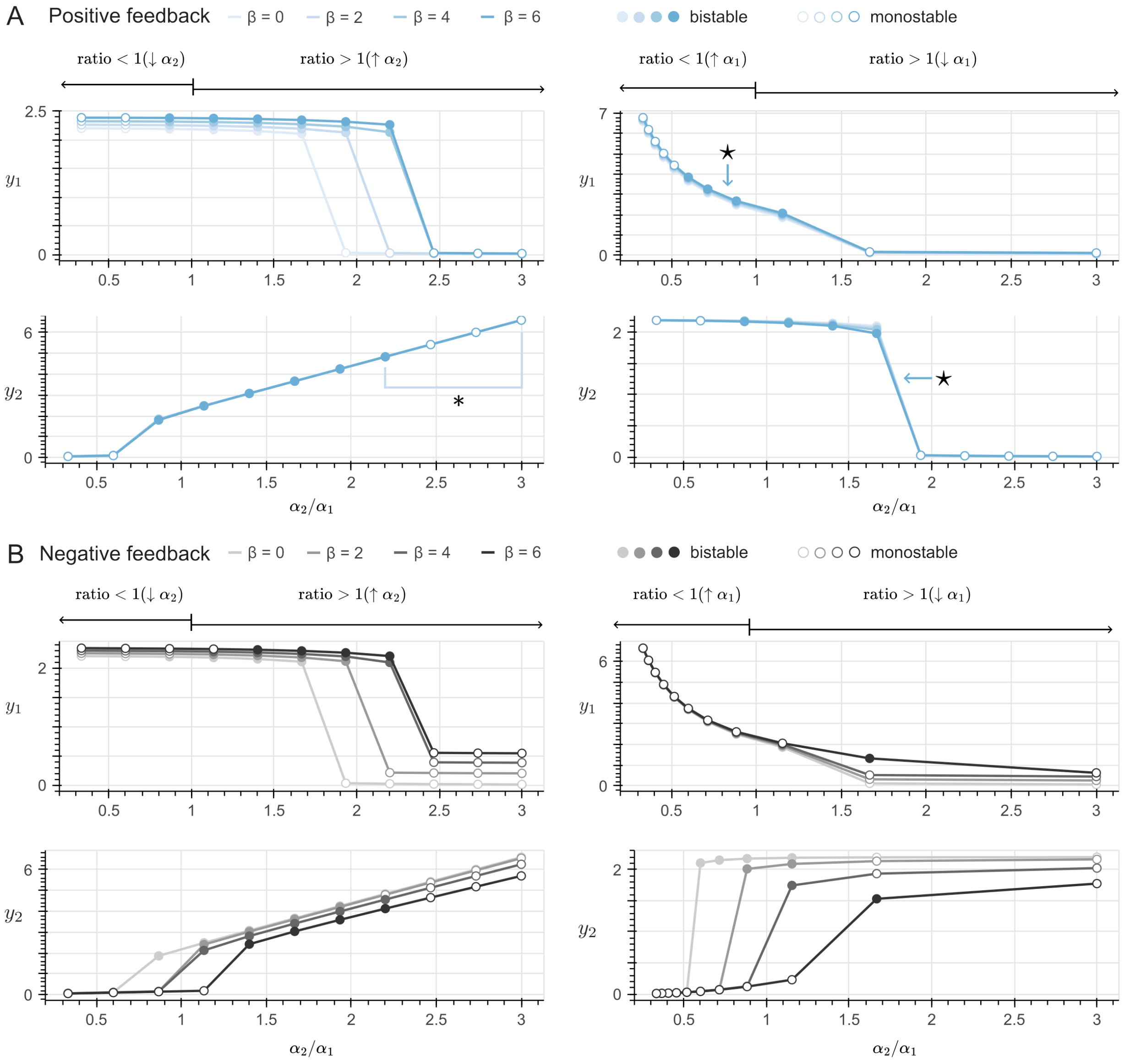

***Figure 10: Increasing the robustness of the negative and positive feedback through the control gain ($\beta$).*** *(A) For mutations in the controlled system with the positive feedback, where ratio > 1 by changing $\alpha_2$ and where ratio < 1 by changes in $\alpha_1$, increasing the $\beta$ correlates with a wider range of $\alpha_2/\alpha_1$ values for which the bistable behavior is maintained. For the bottom left panel, the points highlighted with the (*) symbol correspond to monostability of the light blue lines corresponding to $\beta = 0$ and $\beta = 2$, overlapped by the darker blue lines, since deviations to the equilibrium are within a similar magnitude. Likewise, for the top and bottom right panels, all lines highlighted with the ($\star$) symbol overlap each other, as they exhibit similar stability for mutations where $\alpha_1$ changes (B) For the negative feedback architecture, for mutations where ratio > 1 by changing $\alpha_2$, the strategy of increasing $\beta$ is valid. However, for mutations where ratio < 1 by changes in $\alpha_1$, increasing $\beta$ is counterproductive. In particular, there is single gain value for each $\alpha_2/\alpha_1$ where the system exhibits bistability.*

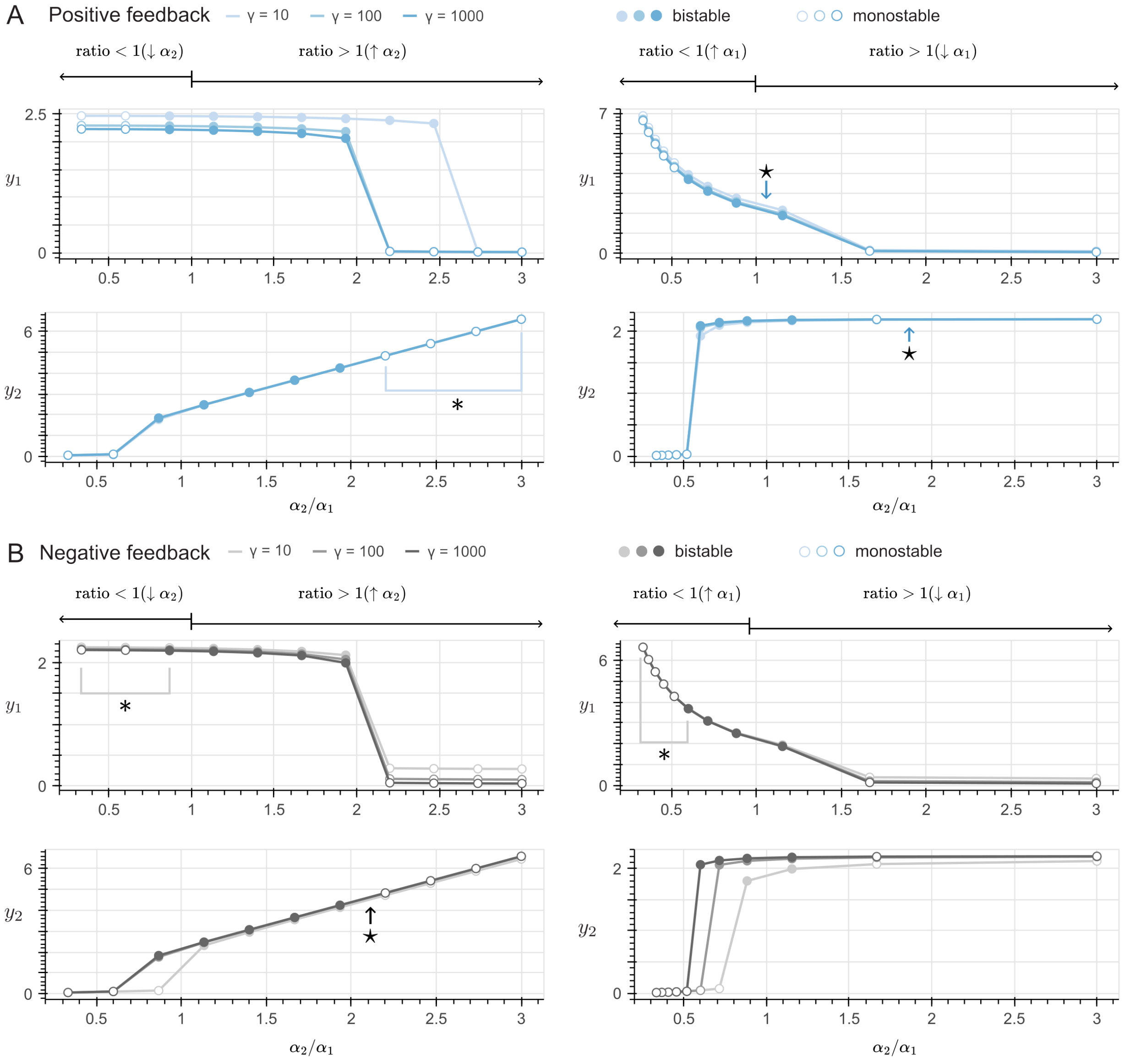

***Figure 11: Increasing the robustness of the negative and positive feedback through the sequestration rate ($\gamma$).*** *(A) For the controlled system with the positive feedback architecture, for mutations where ratio > 1 by changing $\alpha_1$ and ratio < 1 by changes in $\alpha_2$, the range of $\alpha_2/\alpha_1$ values for which the system exhibits bistability can be extended by increasing the sequestration rate (ten-fold for a noticeable change). For the bottom left panel, the points highlighted with the (*) symbol correspond to bistability of the light blue lines corresponding to $\gamma = 10$, overlapped by the darker blue lines, since deviations to the equilibrium are within a similar magnitude. Likewise, for the top and bottom right panels, all line highlighted with the ($\star$) symbol overlap each other, as they exhibit similar stability for mutations where $\alpha_1$ changes (B) For the negative feedback architecture in a very high sequestration regime ($10 \times \gamma$), the range of $\alpha_2/\alpha_1$ values for which the system exhibits bistability is similar to the positive feedback architecture. For the top left and right panels, the points highlighted with the (*) symbol correspond to monostability of the light grey line corresponding to $\gamma = 10$ (top left) and to $\gamma = 10$ and $\gamma = 100$ (top right panel), respectively; overlapped by the darker grey lines, since deviations to the equilibrium are within a similar magnitude. Likewise, for the bottom left panel, all lines highlighted with the ($\star$) symbol overlap each other, as they exhibit similar stability for mutations where $\alpha_2$ changes*

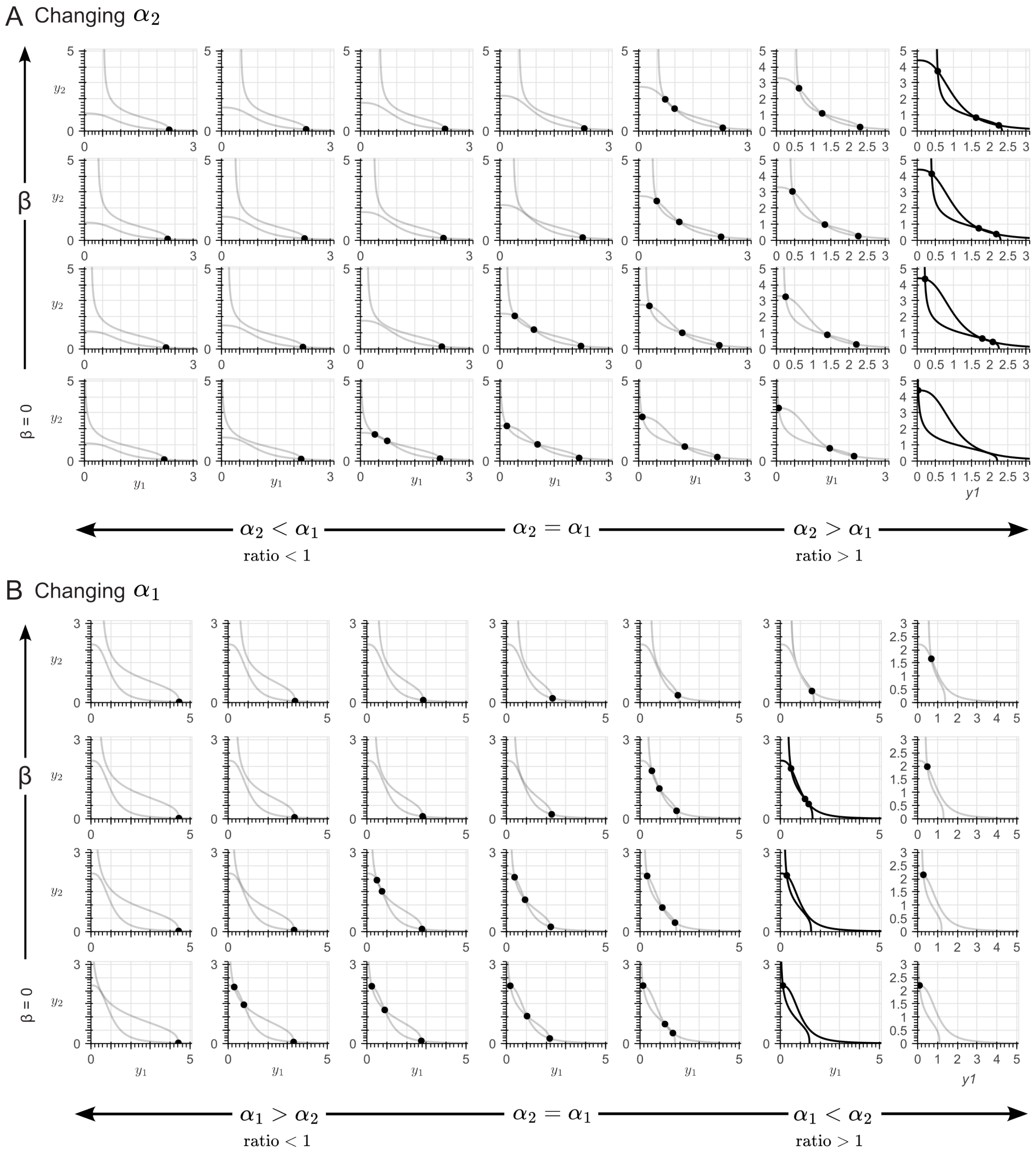

***Figure 12: Phase plane of the controlled system with the negative feedback architecture for increasing control gain ($\beta$) values**. The highlighted cases (from left to right: the last column in Panel A, and the penultimate column in Panel B) illustrate how increasing $\beta$ results in an increased range of $\alpha_2/\alpha_1$ values where bistability is preserved. From bottom to top, the control gain values were $1X$, $2X$ and $3X$ the nominal value of $\beta = 1$.*

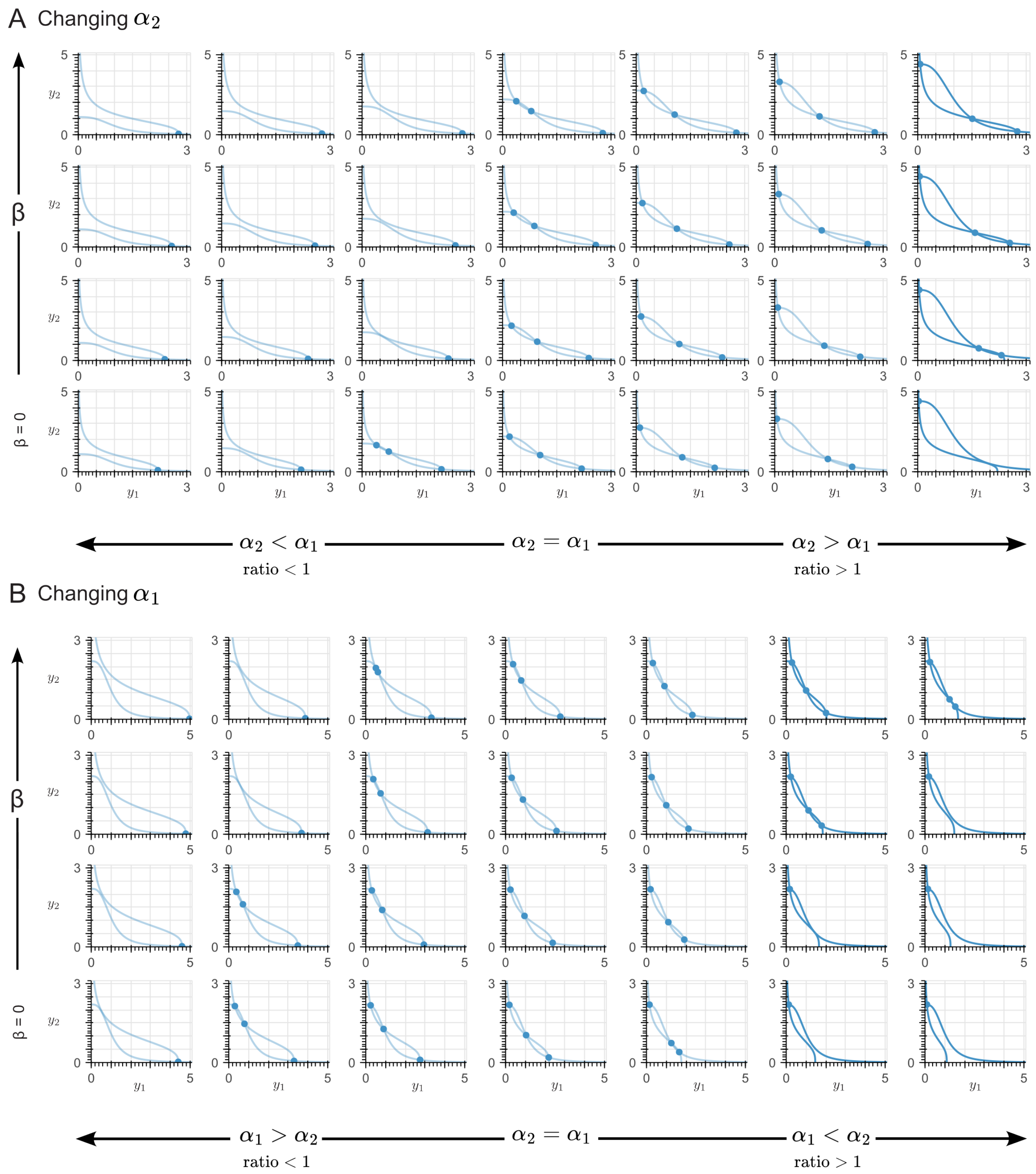

***Figure 13: Phase plane of the controlled system with the positive feedback architecture for increasing control gain ($\beta$) values****. The highlighted cases (from left to right: the last column in Panel A and the last two columns in Panel B) illustrate how increasing the actuation gain ($\beta$) results in an increased range of $\alpha_2/\alpha_1$ values where bistability is maintained. From bottom to top, the control gain values were $1X$, $2X$ and $3X$ the nominal value of $\beta = 1$.*

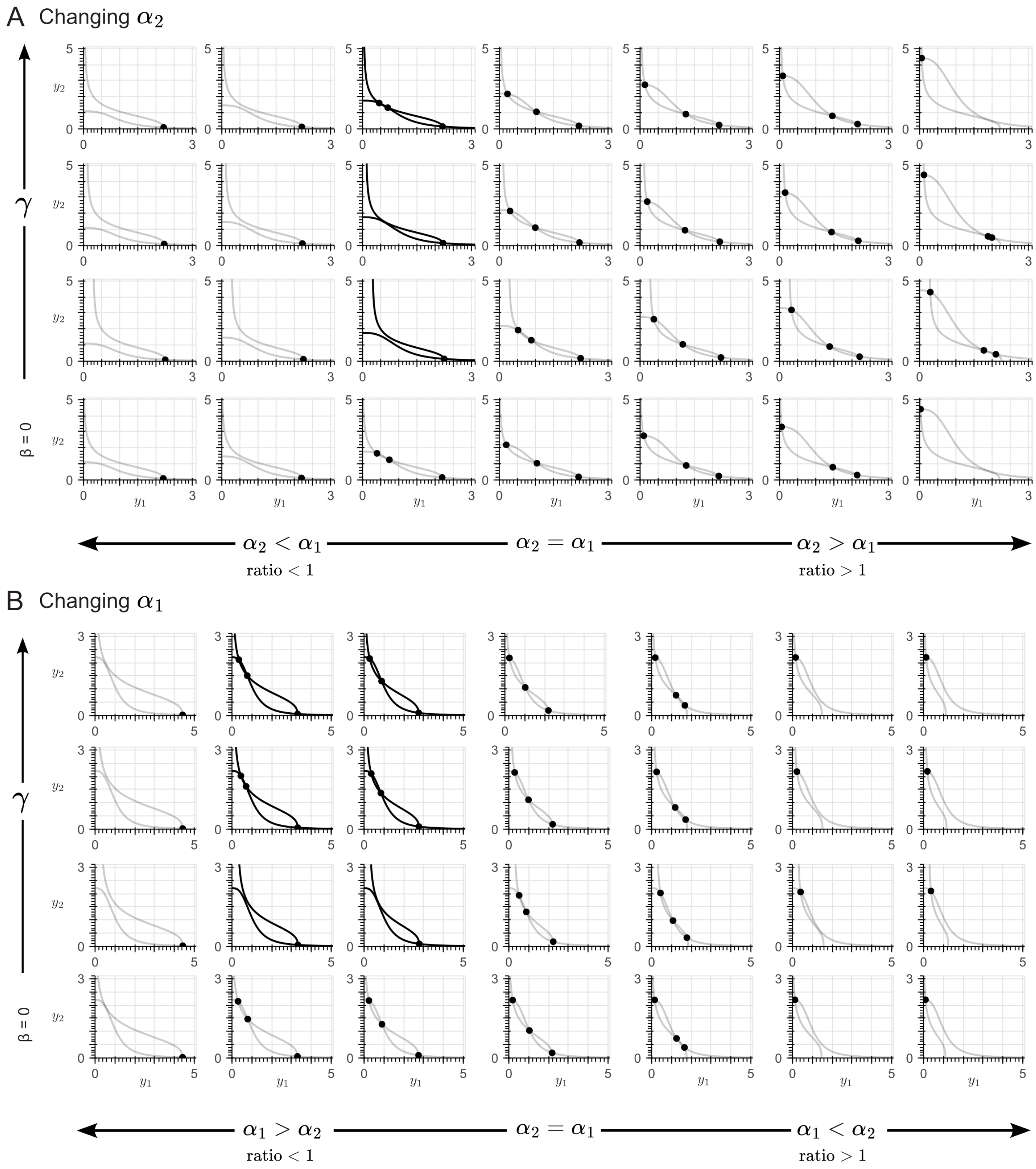

***Figure 14: Phase plane of the controlled system with the negative feedback architecture for increasing sequestration rate ($\gamma$) values.*** *The highlighted cases (from left to right: the third column in Panel A, and the second and third columns in Panel B) illustrates how a significantly faster sequestration rate ($\gamma$) results in an increased range of $\alpha_2/\alpha_1$ values where bistability is maintained. From top to bottom, the sequestration rate values were $0.1X$, $1X$ and $10X$ the nominal value of $\gamma = 100$.*

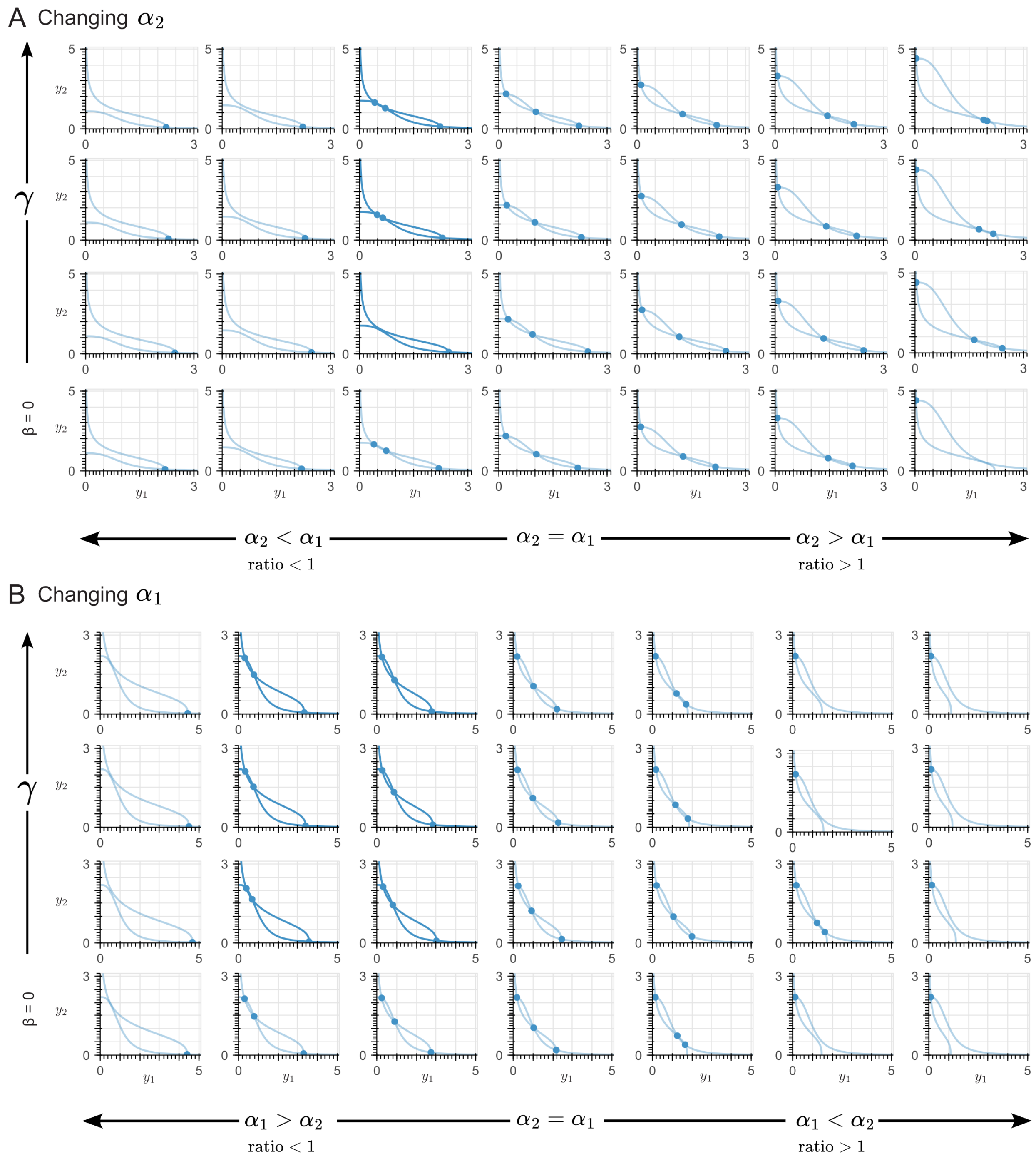

***Figure 15: Phase plane of the controlled system with the positive feedback architecture for increasing sequestration rate ($\gamma$) values.*** *The highlighted cases (from left to right: the third column in Panel A, and the second and third columns in Panel B) illustrates how a significantly faster sequestration rate results in an increased range of $\alpha_2/\alpha_1$ values where bistability is maintained. From top to bottom, the sequestration rate values were $0.1X$, $1X$ and $10X$ the nominal value of $\gamma = 100$.*

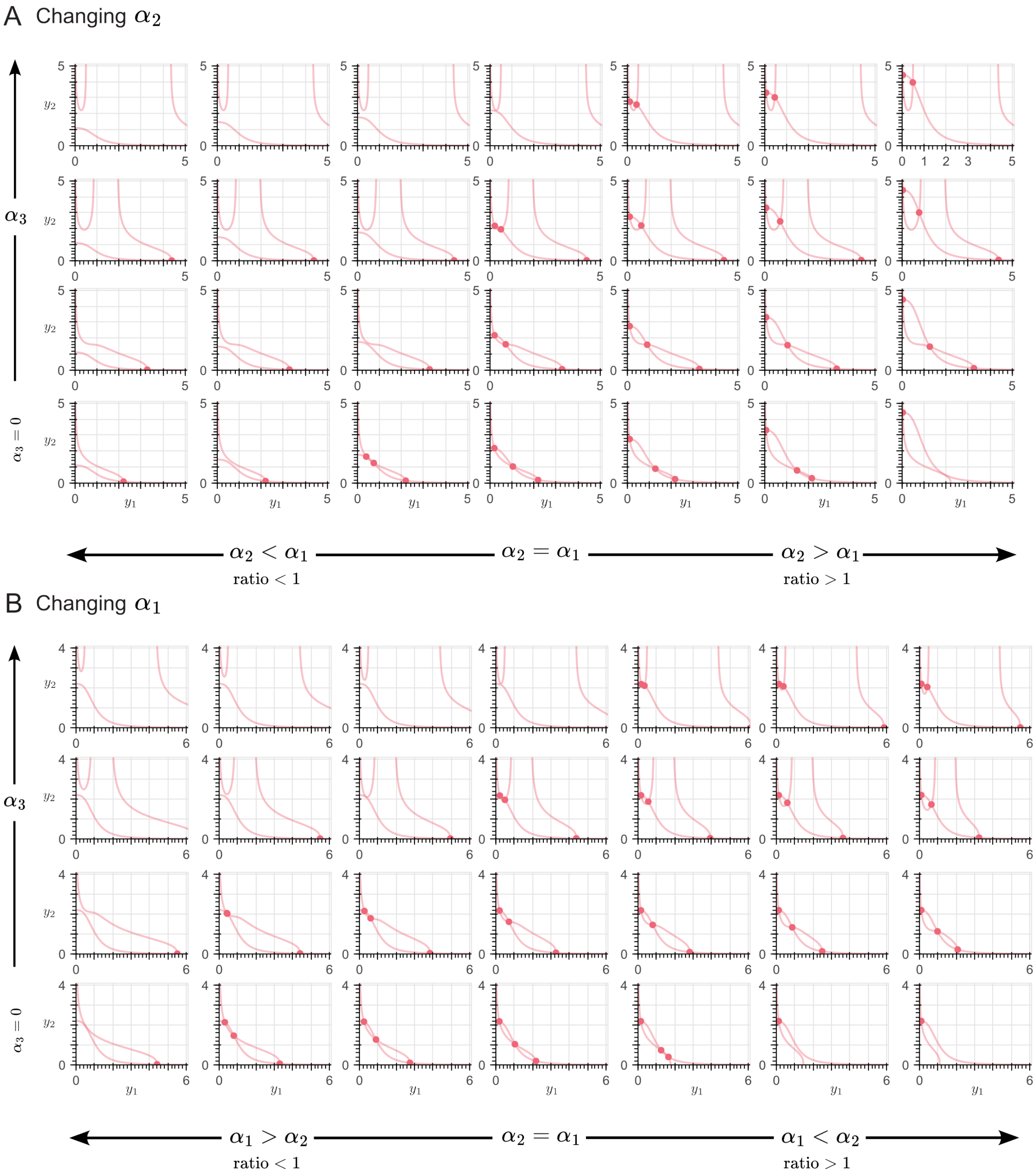

***Figure 16: Phase plane of the toggle switch with a self-activation motif in the $Y_1$ species for increasing values of activation strength ($\alpha_3$).*** *The increased robustness for mutations where ratio > 1 correlates with the modification of the nullclines' shape, observed for high values of $\alpha_3$. From top to bottom, the activation strength values were $1X$, $2X$ and $3X$ the nominal value of $\alpha_3 = 2.2$.*

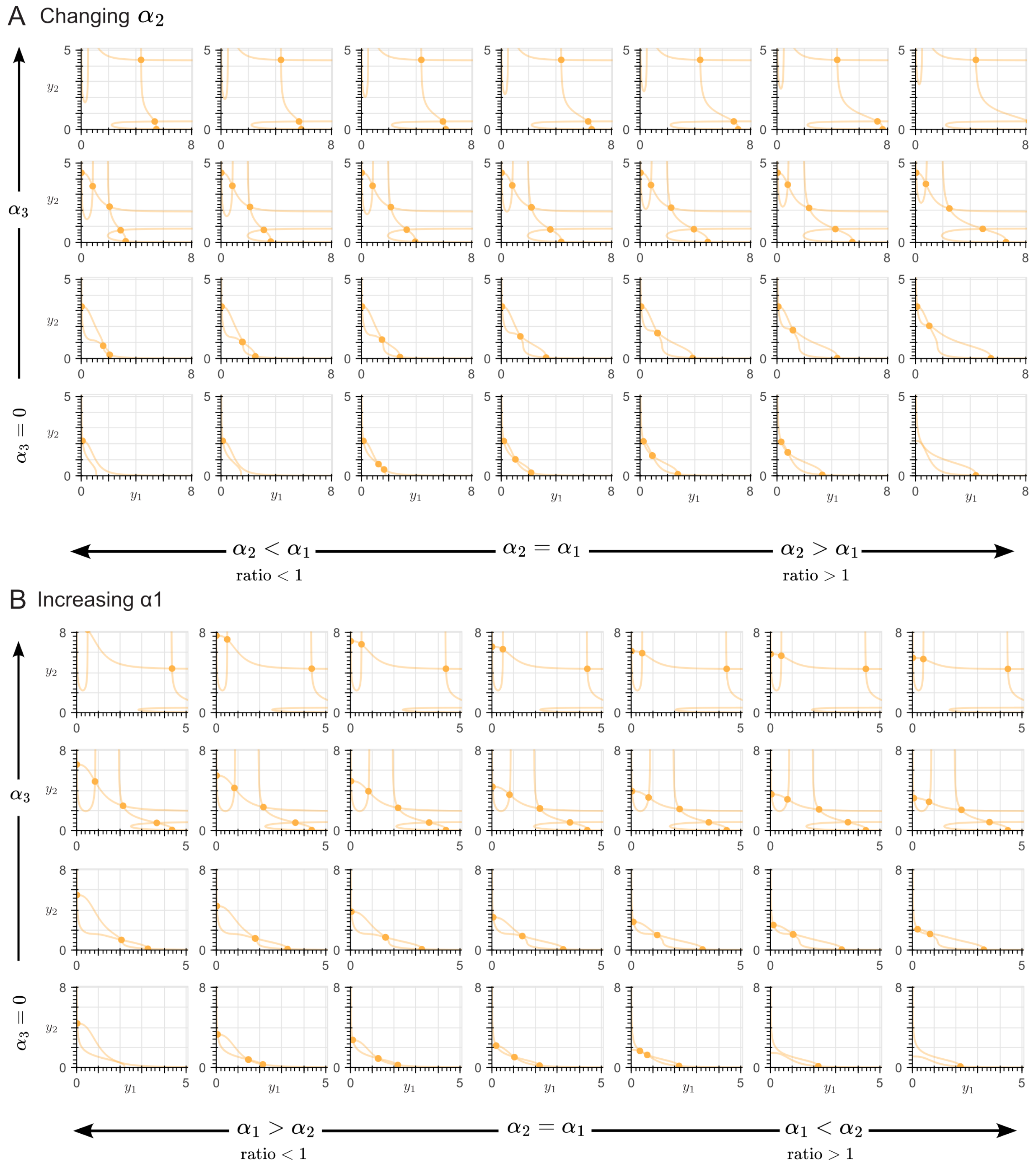

***Figure 17: Phase plane of the toggle switch with a self-activation motif for each species $Y_1$ and $Y_2$, showing the effect of increasing values of activation strength ($\alpha_3$).*** *The dynamics of the network can be divided in two regimes: bistability, observed for low $\alpha_3$ values ($\alpha_3 = 1.1$, for this example) and tristability, in which the self-induction motif induces the generation of a third (intermediate) equilibrium point (for $alpha_3 > 1.1$, for this example). From top to bottom, the activation strength values were $1X$, $2X$ and $3X$ the nominal value of $\alpha_3 = 2.2$.*

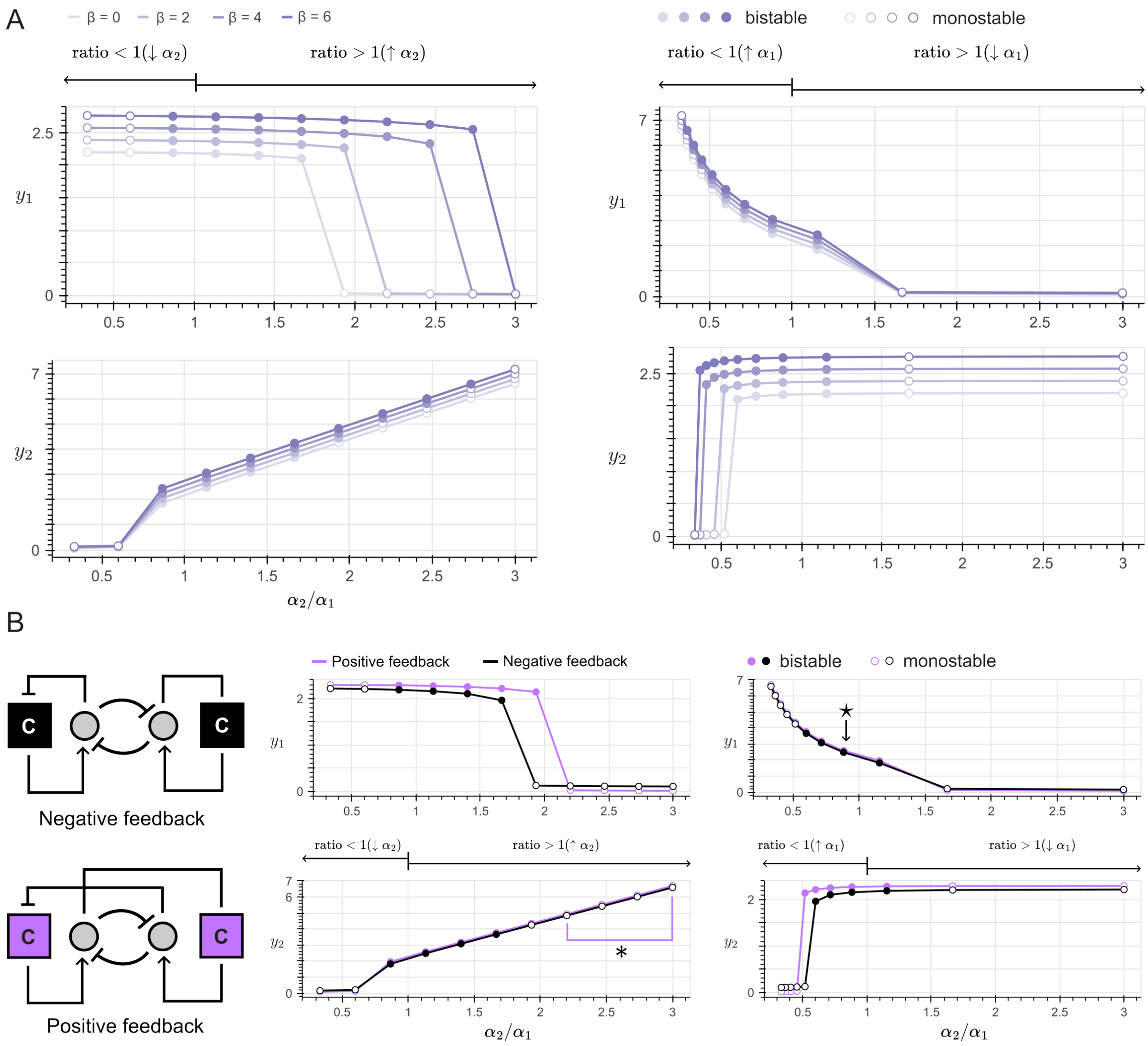

***Figure 18: Increasing robustness through the addition of another adaptive controller*** *(A) The controlled system with two adaptive controllers enforced on each species* $Y_1$ *and* $Y_2$*, with the positive feedback architecture and with a high control gain* ($\beta = \beta' = 6$ *for this example) exhibit robustness to virtually any type of mutation. (B) Nevertheless, the same strategy is not suited for the negative feedback architecture, as the addition of another adaptive controller reduces the* $\alpha_2/\alpha_1$ *range where bistability is maintained, compared to the application of a single adaptive controller. For the bottom left panel, the points highlighted with the (*) symbol correspond to monostability of the purple line, overlapped by the black line, since deviations to the equilibrium are within a similar magnitude. Likewise, for the top right panel, both lines highlighted with the* ($\star$) *symbol overlap each other, as they exhibit similar stability for mutations where* $\alpha_1$ *changes*

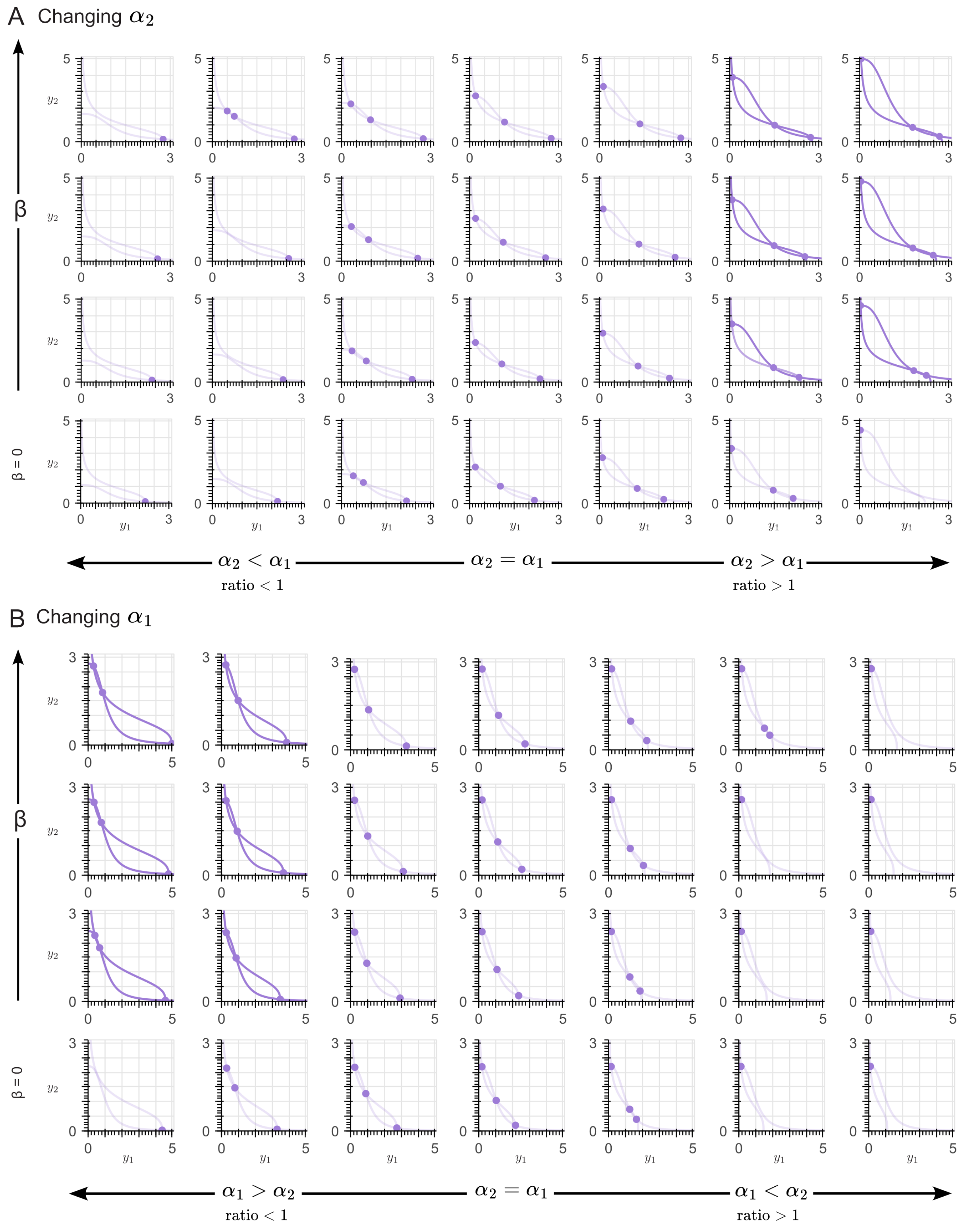

***Figure 19: Phase plane of the controlled system with two adaptive controllers, each with the positive feedback architecture, showing the effect of increasing values of control gain (assumed $\beta = \beta'$ for simplicity).*** *The highlighted cases (from left to right: the last two columns in Panel A, and the first three columns in Panel B) illustrate an overall increase in the system's robustness for both mutations where ratio $< 1$ for increasing values in $\alpha_1$, and where ratio $> 1$ for increasing values of $\alpha_2$. From top to bottom, the control gain values were $1X$, $2X$ and $3X$ the nominal value of $\beta = \beta' = 1$*

### 4 Extending the applications of the adaptive controller

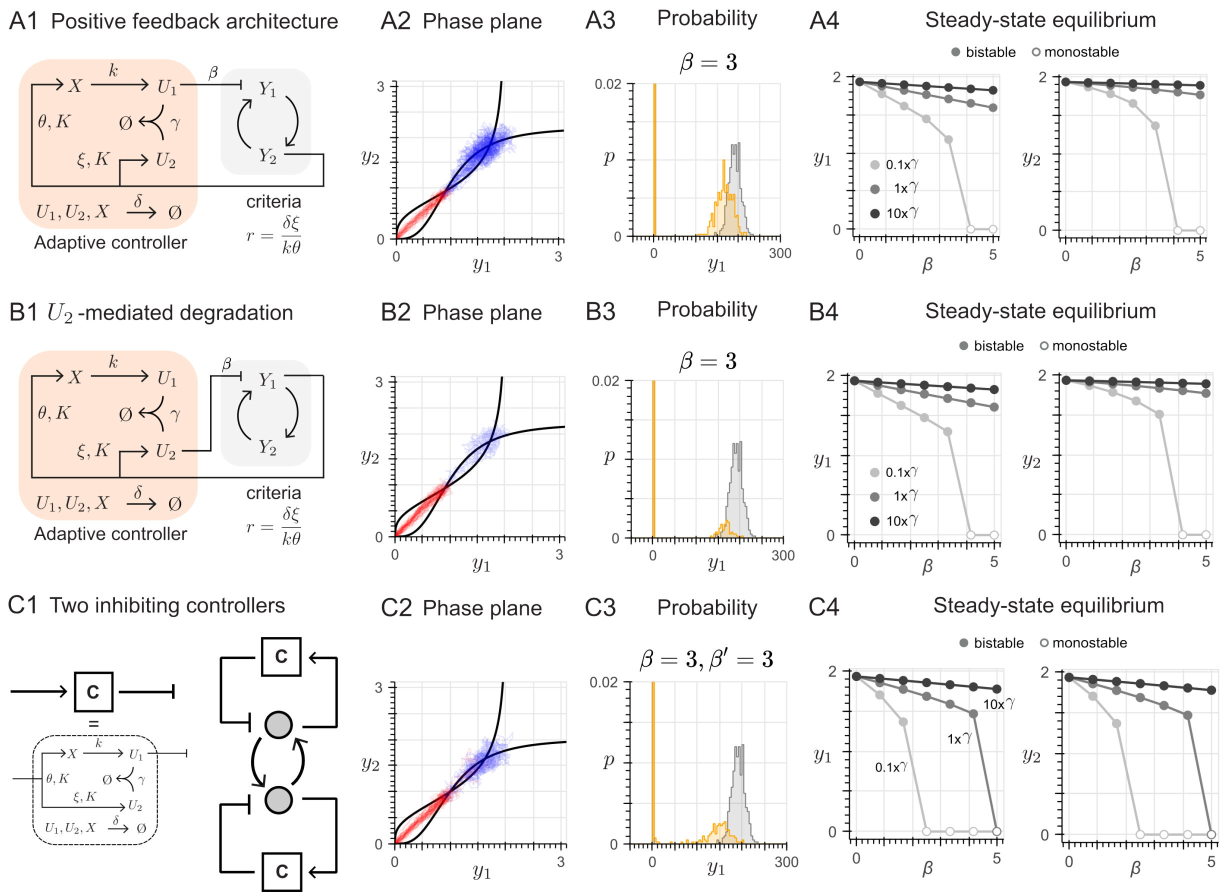

***Figure 20: Alternative control strategies for the mutual induction network*** *(A1) Architecture of the adaptive controller with the positive feedback architecture. (A2) For the controlled system, nullclines (black lines) and trajectories in the phase plane. (A3) Generation of a biased cell fate, which favors the equilibrium with no expression: the probability of converging to the equilibrium with high expression significantly decreases for both species. (A4) Characterization of the effect on the equilibrium value of increasing inhibition strength* $\beta$*, for different choices of the sequestration rate* $\gamma$*. (B1) Architecture of the adaptive controller with an inhibiting effect through species* $U_2$*. (B2)–(B4) Same analysis as in (A2)–(A4) (C1) Architecture of two adaptive controllers with the negative feedback architecture, simultaneously exerting an inhibting effect through species* $U_1$ *and* $U_1'$ *to both species* $Y1$ *and* $Y_2$*, respectively. (C2)–(C4) Same analysis as in (A2)–(A4)*

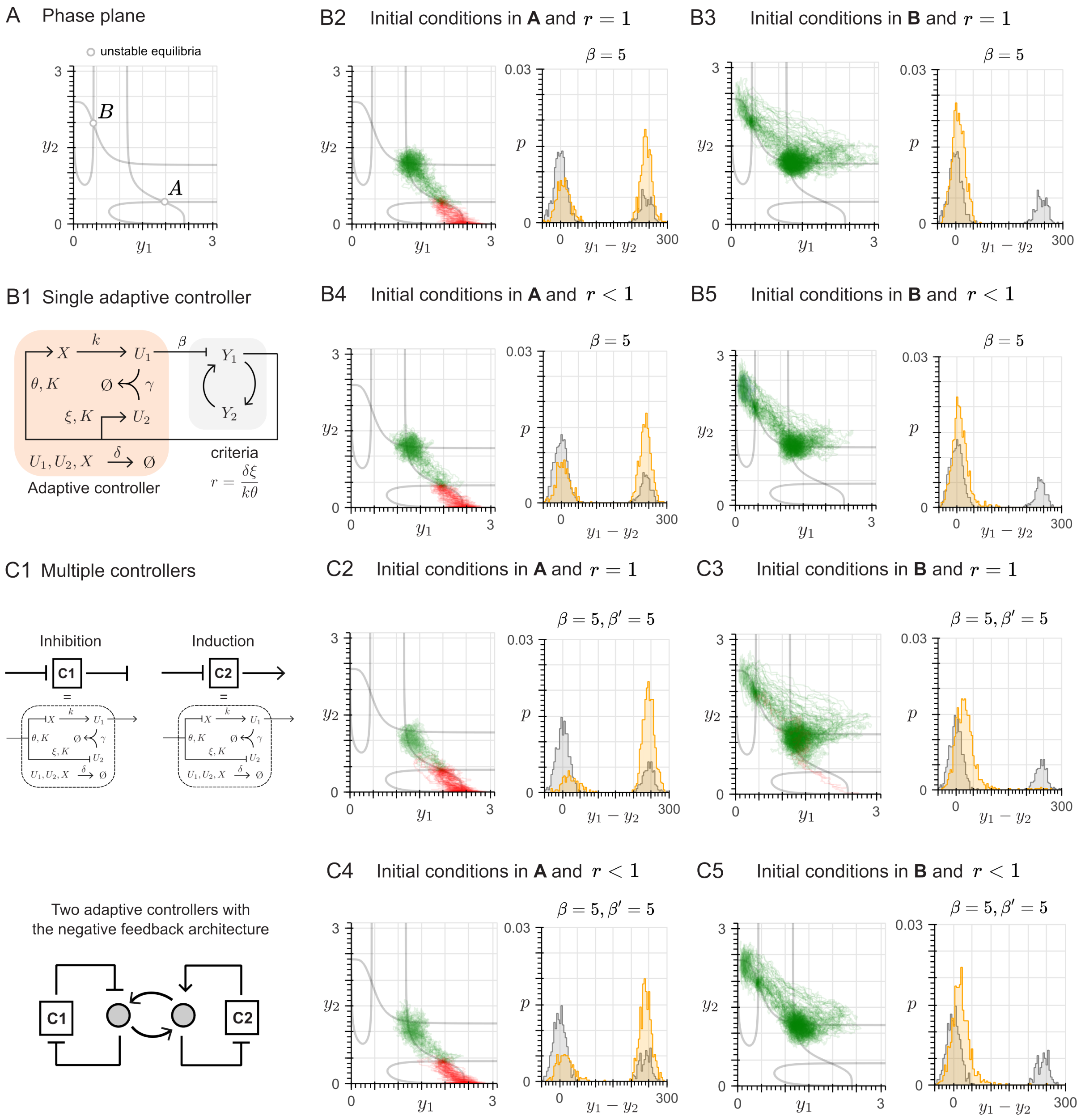

***Figure 21: Design challenge: how can we favor the high production state of either $Y_1$ or $Y_2$?*** *(A1) Initial conditions that comprise the study cases (see main text). (A2) Architecture of the adaptive controller in negative feedback, based in the toggle switch example. (A3) Architecture of the inhibition-induction pair, designed to overcome the limitations of the single controller approach. (B) When the initial conditions are located in the A unstable equilibrium, (B1) a local bistable system emerges, with a bimodal distribution that can be biased through the application of either a single controller or (B2) the inhibition-induction pair. (C) When the initial conditions are located in the coordinate´s origin, (C1) the single controller favors instead the intermediate state, while (C2) the inhibition-induction pair is able to equate the probability of adopting the intermediate state and the equilibrium of high production of the $Y_1$ specie. Nevertheless, a biased probability distribution can´t be generated in either case. (D) When the initial conditions are located in the C unstable equilibrium, neither the (D1) single controller or the (D2) induction-inhibition pair are capable of biasing, resulting in the waterbed effect. All simulations were done for an actuation gain $\beta = \beta' = 5$.*

### 5 Models

#### 5.1 Adaptive controller with negative feedback and $U_1$ actuation for the toggle switch case study

This is the network introduced in the section *2.1.2 Toggle switch: model description* in the main text. Chemical reactions and ordinary differential equations are repeated here for ease of reference.

##### Chemical reactions

| | | |
|---|---|---|
| $\emptyset \xrightarrow{\alpha p_1} Y_1,$ | $\emptyset \xrightarrow{\alpha p_2} Y_2,$ | Production/Production, |
| $Y_1 \xrightarrow{\delta} \emptyset,$ | $Y_2 \xrightarrow{\delta} \emptyset,$ | Decay/Decay |
| $Y_1 \xrightarrow{\theta p_2} Y_1 + X$ | $Y_1 \xrightarrow{\xi p_2} Y_1 + U_2$ | Production/Production |
| $U_1 \xrightarrow{\beta} U_1 + Y_1$ | $U_1 + U_2 \xrightarrow{\gamma} \emptyset$ | Production/Sequestration |
| $U_1, U_2 \xrightarrow{\delta} \emptyset$ | $X \xrightarrow{\delta} \emptyset$ | Decay/Decay |

where production rate constants are defined as $p_1 = \frac{K^m}{y_2^m + K^m}$, and $p_2 = \frac{K^m}{y_1^m + K^m}$.

##### Ordinary Differential Equations

$$\dot{y}_1 = \alpha \frac{K^m}{y_2^m + K^m} - \delta y_1 + \beta u_1 \tag{24}$$

$$\dot{y}_2 = \alpha \frac{K^m}{y_1^m + K^m} - \delta y_2 \tag{25}$$

$$\dot{u}_1 = kx - \delta u_1 - \gamma u_1 u_2 \tag{26}$$

$$\dot{u}_2 = \xi \frac{K^m}{K^m + y_1^m} - \delta u_2 - \gamma u_1 u_2 \tag{27}$$

$$\dot{x} = \theta \frac{K^m}{K^m + y_1^m} - \delta x \tag{28}$$

##### Steady-state analysis

To determine the system´s phase plane, we calculate the nullclines by equating $\dot{y}_1 = 0$ and $\dot{y}_2 = 0$, obtaining the following expressions:

$$\bar{y}_1 = \sqrt[m]{\frac{\alpha K^m}{\delta \bar{y}_2} - K^m}$$

$$\bar{y}_2 = \sqrt[m]{\frac{\alpha K^m}{\delta \bar{y}_1 - \beta \bar{u}_1} - K^m}$$

We can further find $\bar{u}_1$ as a function of $\bar{y}_1$.

$$\bar{u}_2 = \frac{f_1 - \delta \bar{u}_1}{\gamma \bar{u}_1} = \frac{f_2}{\gamma \bar{u}_1 + \delta},$$

where $f_1 = k\bar{x}$, $f_2 = \xi\frac{K^m}{K^m+\bar{y}_1^m}$, and $\bar{x} = \frac{\theta}{\delta}\frac{K^m}{K^m+\bar{y}_1^m}$. This leads to a second order polynomial, $P(\bar{u}_1) = \bar{u}_1^2 + A\bar{u}_1 + B = 0$, and to a single positive real solution

$$\bar{u}_1 = \frac{-A + \sqrt{A^2 - 4B}}{2}$$

where $A = \frac{f_2 - f_1}{\delta} + \frac{\delta}{\gamma}$, and $B = -\frac{f_1}{\gamma}$.

#### 5.2 Adaptive controller with positive feedback and $U_1$ actuation for the toggle switch case study

This is a modification of the previous network regarding the feedback specie, detailed in the main text and characterized in Fig. S5.

##### Chemical reactions

| | | |
|---|---|---|
| $\emptyset \xrightarrow{\alpha p_1} Y_1,$ | $\emptyset \xrightarrow{\alpha p_2} Y_2,$ | Production, |
| $Y_1 \xrightarrow{\delta} \emptyset,$ | $Y_2 \xrightarrow{\delta} \emptyset,$ | Decay |
| $Y_2 \xrightarrow{\theta p_1} Y_2 + X$ | $Y_2 \xrightarrow{\xi p_1} Y_2 + U_2$ | Production |
| $U_1 \xrightarrow{\beta} U_1 + Y_1$ | $U_1 + U_2 \xrightarrow{\gamma} \emptyset$ | Production/Sequestration |
| $U_1, U_2 \xrightarrow{\delta} \emptyset$ | $X \xrightarrow{\delta} \emptyset$ | Decay |

where production rate constants are defined as $p_1 = \frac{K^m}{y_2^m + K^m}$, and $p_2 = \frac{K^m}{y_1^m + K^m}$.

##### Ordinary Differential Equations

$$\dot{y}_1 = \alpha\frac{K^m}{y_2^m + K^m} - \delta y_1 + \beta u_1 \quad (29)$$

$$\dot{y}_2 = \alpha\frac{K^m}{y_1^m + K^m} - \delta y_2 \quad (30)$$

$$\dot{u}_1 = kx - \delta u_1 - \gamma u_1 u_2 \quad (31)$$

$$\dot{u}_2 = \xi\frac{K^m}{K^m + y_2^m} - \delta u_2 - \gamma u_1 u_2 \quad (32)$$

$$\dot{x} = \theta\frac{K^m}{K^m + y_2^m} - \delta x \quad (33)$$

##### Steady-state analysis

To determine the system´s phase plane, we calculate the nullclines by equating $\dot{y}_1 = 0$ and $\dot{y}_2 = 0$, obtaining the following expressions:

$$\bar{y}_2 = \frac{\alpha}{\delta}\left(\frac{K^m}{K^m + \bar{y}_1^m}\right)$$

$$\bar{y}_1 = \frac{\alpha}{\delta}\frac{K^m}{K^m + \bar{y_2}^m} + \frac{\beta}{\delta}u_1$$

We can further find $\bar{u}_1$ as a function of $\bar{y}_2$.

$$\bar{u}_2 = \frac{f_1 - \delta \bar{u}_1}{\gamma \bar{u}_1} = \frac{f_2}{\gamma \bar{u}_1 + \delta},$$

where $f_1 = k\bar{x}$, $f_2 = \xi \frac{K^m}{K^m + \bar{y}_2^m}$, and $\bar{x} = \frac{\theta}{\delta} \frac{K^m}{K^m + \bar{y}_2^m}$. This leads to a second order polynomial, $P(\bar{u}_1) = \bar{u}_1^2 + A\bar{u}_1 + B = 0$, and leads to a single positive real solution

$$\bar{u}_1 = \frac{-A + \sqrt{A^2 - 4B}}{2}$$

where $A = \frac{f_2 - f_1}{\delta} + \frac{\delta}{\gamma}$, and $B = -\frac{f_1}{\gamma}$.

#### 5.3 Adaptive controller with positive feedback and $U_2$ actuation for the toggle switch case study

This is a modification of the negative feedback architecture regarding the actuation specie (switch from $u_1$ to $u_2$), detailed in the main text and characterized in Fig. S7.

##### Chemical reactions

| | | |
|---|---|---|
| $\emptyset \xrightarrow{\alpha p_1} Y_1,$ | $\emptyset \xrightarrow{\alpha p_2} Y_2,$ | Production, |
| $Y_1 \xrightarrow{\delta} \emptyset,$ | $Y_2 \xrightarrow{\delta} \emptyset,$ | Decay |
| $Y_2 \xrightarrow{\theta p_1} Y_2 + X$ | $Y_2 \xrightarrow{\xi p_1} Y_2 + U_2$ | Production |
| $U_2 \xrightarrow{\beta} U_2 + Y_1$ | $U_1 + U_2 \xrightarrow{\gamma} \emptyset$ | Production/Sequestration |
| $U_1, U_2 \xrightarrow{\delta} \emptyset$ | $X \xrightarrow{\delta} \emptyset$ | Decay |

where production rate constants are defined as $p_1 = \frac{K^m}{y_2^m + K^m}$, and $p_2 = \frac{K^m}{y_1^m + K^m}$.

##### Ordinary Differential Equations

$$\dot{y}_1 = \alpha \frac{K^m}{y_2^m + K^m} - \delta y_1 + \beta u_2 \quad (34)$$

$$\dot{y}_2 = \alpha \frac{K^m}{y_1^m + K^m} - \delta y_2 \quad (35)$$

$$\dot{u}_1 = kx - \delta u_1 - \gamma u_1 u_2 \quad (36)$$

$$\dot{u}_2 = \xi \frac{K^m}{K^m + y_2^m} - \delta u_2 - \gamma u_1 u_2 \quad (37)$$

$$\dot{x} = \theta \frac{K^m}{K^m + y_2^m} - \delta x \quad (38)$$

##### Steady-state analysis

To determine the system´s phase plane, we calculate the nullclines by equating $\dot{y}_1 = 0$ and $\dot{y}_2 = 0$, obtaining the following expressions:

$$\begin{aligned}\bar{y}_2 &= \frac{\alpha}{\delta}\left(\frac{K^m}{K^m+\bar{y}_1^m}\right)\\ \bar{y}_1 &= \frac{\alpha}{\delta}\frac{K^m}{K^m+\bar{y_2}^m}+\frac{\beta}{\delta}u_2\end{aligned}$$

We can further find $\bar{u}_2$ as a function of $\bar{y}_2$.

$$\bar{u}_1 = \frac{f_2-\delta\bar{u}_2}{\gamma\bar{u}_2} = \frac{f_1}{\gamma\bar{u}_2+\delta},$$

where $f_1 = k\bar{x}$, $f_2 = \xi\frac{K^m}{K^m+\bar{y}_2^m}$, and $\bar{x} = \frac{\theta}{\delta}\frac{K^m}{K^m+\bar{y}_2^m}$. This leads to a second order polynomial, $P(\bar{u}_2) = \bar{u}_2^2 + A\bar{u}_2 + B = 0$, and leads to a single positive real solution

$$\bar{u}_2 = \frac{-A+\sqrt{A^2-4B}}{2}$$

where $A = \frac{f_1-f_2}{\delta} + \frac{\delta}{\gamma}$, and $B = -\frac{f_2}{\gamma}$.

#### 5.4 Adaptive controller with negative feedback and $U_2$ actuation for the toggle switch case study

This is a modification of the positive feedback architecture regarding the actuation specie (switch from $u_1$ to $u_2$), detailed in the main text and characterized in Fig. S8.

##### Chemical reactions

| | | |
|---|---|---|
| $\emptyset \xrightarrow{\alpha p_1} Y_1,$ | $\emptyset \xrightarrow{\alpha p_2} Y_2,$ | Production, |
| $Y_1 \xrightarrow{\delta} \emptyset,$ | $Y_2 \xrightarrow{\delta} \emptyset,$ | Decay |
| $Y_1 \xrightarrow{\theta p_2} Y_1 + X$ | $Y_1 \xrightarrow{\xi p_2} Y_1 + U_2$ | Production |
| $U_2 \xrightarrow{\beta} U_2 + Y_1$ | $U_1 + U_2 \xrightarrow{\gamma} \emptyset$ | Production/Sequestration |
| $U_1, U_2 \xrightarrow{\delta} \emptyset$ | $X \xrightarrow{\delta} \emptyset$ | Decay |

where production rate constants are defined as $p_1 = \frac{K^m}{y_2^m+K^m}$, and $p_2 = \frac{K^m}{y_1^m+K^m}$.

###### Ordinary Differential Equations

$$\begin{aligned}
\dot{y}_1 &= \alpha\frac{K^m}{y_2^m + K^m} - \delta y_1 + \beta u_2 && (39)\\
\dot{y}_2 &= \alpha\frac{K^m}{y_1^m + K^m} - \delta y_2 && (40)\\
\dot{u}_1 &= kx - \delta u_1 - \gamma u_1 u_2 && (41)\\
\dot{u}_2 &= \xi\frac{K^m}{K^m + y_1^m} - \delta u_2 - \gamma u_1 u_2 && (42)\\
\dot{x} &= \theta\frac{K^m}{K^m + y_1^m} - \delta x && (43)
\end{aligned}$$

###### Steady-state analysis

To determine the system´s phase plane, we calculate the nullclines by equating $\dot{y}_1 = 0$ and $\dot{y}_2 = 0$, obtaining the following expressions:

$$\begin{aligned}
\bar{y}_1 &= \sqrt[m]{\frac{\alpha K^m}{\delta \bar{y}_2} - K^m}\\
\bar{y}_2 &= \sqrt[m]{\frac{\alpha K^m}{\delta \bar{y}_1 - \beta \bar{u}_2} - K^m}
\end{aligned}$$

We can further find $\bar{u}_2$ as a function of $\bar{y}_1$.

$$\bar{u}_1 = \frac{f_2 - \delta \bar{u}_2}{\gamma \bar{u}_2} = \frac{f_1}{\gamma \bar{u}_2 + \delta},$$

where $f_1 = k\bar{x}$, $f_2 = \xi\frac{K^m}{K^m + \bar{y}_1^m}$, and $\bar{x} = \frac{\theta}{\delta}\frac{K^m}{K^m + \bar{y}_1^m}$. This leads to a second order polynomial, $P(\bar{u}_2) = \bar{u}_2^2 + A\bar{u}_2 + B = 0$, and leads to a single positive real solution

$$\bar{u}_2 = \frac{-A + \sqrt{A^2 - 4B}}{2}$$

where $A = \frac{f_1 - f_2}{\delta} + \frac{\delta}{\gamma}$, and $B = -\frac{f_2}{\gamma}$.

#### 5.5 Adaptive controller for the mutual induction case study

These are the networks studied in the subsection *2.3.1 Applying the adaptive control principle to a different system* in the main text, and characterized in Fig. S20.

##### 5.5.1 Single inhibiting controller in a negative feedback with actuation through $U_1$ specie

###### Chemical reactions

| | | |
|---|---|---|
| $\emptyset \xrightarrow{\alpha p_1} Y_1,$ | $\emptyset \xrightarrow{\alpha p_2} Y_2,$ | Production, |
| $Y_1 \xrightarrow{\delta} \emptyset,$ | $Y_2 \xrightarrow{\delta} \emptyset,$ | Decay |
| $Y_1 \xrightarrow{\theta p_2} Y_1 + X$ | $Y_1 \xrightarrow{\xi p_2} Y_1 + U_2$ | Production |
| $U_1 + Y_1 \xrightarrow{\beta} \emptyset$ | $U_1 + U_2 \xrightarrow{\gamma} \emptyset$ | Decay/Sequestration |
| $U_1, U_2 \xrightarrow{\delta} \emptyset$ | $X \xrightarrow{\delta} \emptyset$ | Decay |

where production rate constants are defined as $p_1 = \frac{y_2^m}{y_2^m + K^m}$, and $p_2 = \frac{y_1^m}{y_1^m + K^m}$.

###### Ordinary Differential Equations

$$\dot{y}_1 = \alpha \frac{y_2^m}{y_2^m + K^m} - \delta y_1 - \beta u_1 \frac{y_1}{y_1 + K} \tag{44}$$

$$\dot{y}_2 = \alpha \frac{y_1^m}{y_1^m + K^m} - \delta y_2 \tag{45}$$

$$\dot{u}_1 = kx - \delta u_1 - \gamma u_1 u_2 \tag{46}$$

$$\dot{u}_2 = \xi \frac{y_1^m}{K^m + y_1^m} - \delta u_2 - \gamma u_1 u_2 \tag{47}$$

$$\dot{x} = \theta \frac{y_1^m}{K^m + y_1^m} - \delta x \tag{48}$$

###### Steady-state analysis

To determine the system´s phase plane, we calculate the nullclines by equating $\dot{y}_1 = 0$ and $\dot{y}_2 = 0$, obtaining the following expressions:

$$\bar{y}_1 = \sqrt[m]{\frac{\delta \bar{y}_2 K^m}{\alpha - \delta \bar{y}_2}}$$

$$\bar{y}_2 = \sqrt[m]{\frac{h K^m}{\alpha - h}}$$

where $h = \delta \bar{y}_1 + \beta \bar{u}_1 \frac{\bar{y}_1}{\bar{y}_1 + K}$. We can further find $\bar{u}_1$ as a function of $\bar{y}_1$

$$\bar{u}_2 = \frac{f_1 - \delta \bar{u}_1}{\gamma \bar{u}_1} = \frac{f_2}{\gamma \bar{u}_1 + \delta},$$

where $f_1 = k\bar{x}$, $f_2 = \xi \frac{y_1^m}{K^m + \bar{y}_1^m}$, and $\bar{x} = \frac{\theta}{\delta} \frac{y_1^m}{K^m + \bar{y}_1^m}$. This leads to a second order polynomial, $P(\bar{u}_1) = \bar{u}_1^2 + A\bar{u}_1 + B = 0$, and leads to a single positive real solution

$$\bar{u}_1 = \frac{-A + \sqrt{A^2 - 4B}}{2}$$

where $A = \frac{f_2 - f_1}{\delta} + \frac{\delta}{\gamma}$, and $B = -\frac{f_1}{\gamma}$.

##### 5.5.2 Single inhibiting controller in a positive feedback with actuation through $U_1$ specie

###### Chemical reactions

| | | |
|---|---|---|
| $\emptyset \xrightarrow{\alpha p_1} Y_1,$ | $\emptyset \xrightarrow{\alpha p_2} Y_2,$ | Production, |
| $Y_1 \xrightarrow{\delta} \emptyset,$ | $Y_2 \xrightarrow{\delta} \emptyset,$ | Decay |
| $Y_2 \xrightarrow{\theta p_1} Y_2 + X$ | $Y_1 \xrightarrow{\xi p_1} Y_2 + U_2$ | Production |
| $U_1 + Y_1 \xrightarrow{\beta} \emptyset$ | $U_1 + U_2 \xrightarrow{\gamma} \emptyset$ | Decay/Sequestration |
| $U_1, U_2 \xrightarrow{\delta} \emptyset$ | $X \xrightarrow{\delta} \emptyset$ | Decay |

where production rate constants are defined as $p_1 = \frac{y_2^m}{y_2^m + K^m}$, and $p_2 = \frac{y_1^m}{y_1^m + K^m}$.

###### Ordinary Differential Equations

$$\begin{aligned}
\dot{y}_1 &= \alpha \frac{y_2^m}{y_2^m + K^m} - \delta y_1 - \beta u_1 \frac{y_1}{y_1 + K} \\
\dot{y}_2 &= \alpha \frac{y_1^m}{y_1^m + K^m} - \delta y_2 \\
\dot{u}_1 &= kx - \delta u_1 - \gamma u_1 u_2 \\
\dot{u}_2 &= \xi \frac{y_2^m}{K^m + y_2^m} - \delta u_2 - \gamma u_1 u_2 \\
\dot{x} &= \theta \frac{y_2^m}{K^m + y_2^m} - \delta x
\end{aligned}$$

###### Steady-state analysis

To determine the system´s phase plane, we calculate the nullclines by equating $\dot{y}_1 = 0$ and $\dot{y}_2 = 0$, obtaining the following expressions:

$$\begin{aligned}
\dot{y}_1 &= \alpha \frac{\bar{y}_2^m}{\bar{y}_2^m + K^m} - \delta \bar{y}_1 - \beta \bar{u}_1 \frac{\bar{y}_1}{\bar{y}_1 + K} = 0 \\
\bar{y}_2 &= \frac{\alpha}{\delta} \frac{\bar{y}_1^m}{\bar{y}_1^m + K^m}
\end{aligned}$$

The solution of $\dot{y}_1 = 0$ in terms of $\bar{y}_1$ has a complex closed-form analytical expression, so we applied numerical methods. Since the adaptive controller is essentially an exogenous circuit, we can still find $\bar{u}_1$ as a function of $\bar{y}_2$,

$$\bar{u}_2 = \frac{f_1 - \delta \bar{u}_1}{\gamma \bar{u}_1} = \frac{f_2}{\gamma \bar{u}_1 + \delta},$$

where $f_1 = k\bar{x}$, $f_2 = \xi \frac{y_2^m}{K^m + \bar{y}_2^m}$, and $\bar{x} = \frac{\theta}{\delta} \frac{y_2^m}{K^m + \bar{y}_2^m}$. This leads to a second order polynomial, $P(\bar{u}_1) = \bar{u}_1^2 + A\bar{u}_1 + B = 0$, and leads to a single positive real solution

$$\bar{u}_1 = \frac{-A + \sqrt{A^2 - 4B}}{2}$$

where $A = \frac{f_2 - f_1}{\delta} + \frac{\delta}{\gamma}$, and $B = -\frac{f_1}{\gamma}$.

##### 5.5.3 Single inhibiting controller in a negative feedback with actuation through $U_2$ specie

###### Chemical reactions

$$
\begin{array}{lll}
\emptyset \xrightarrow{\alpha p_1} Y_1, & \emptyset \xrightarrow{\alpha p_2} Y_2, & \text{Production,} \\
Y_1 \xrightarrow{\delta} \emptyset, & Y_2 \xrightarrow{\delta} \emptyset, & \text{Decay} \\
Y_1 \xrightarrow{\theta p_2} Y_1 + X & Y_1 \xrightarrow{\xi p_2} Y_1 + U_2 & \text{Production} \\
U_2 + Y_1 \xrightarrow{\beta} \emptyset & U_1 + U_2 \xrightarrow{\gamma} \emptyset & \text{Decay/Sequestration} \\
U_1, U_2 \xrightarrow{\delta} \emptyset & X \xrightarrow{\delta} \emptyset & \text{Decay}
\end{array}
$$

where production rate constants are defined as $p_1 = \frac{y_2^m}{y_2^m + K^m}$, and $p_2 = \frac{y_1^m}{y_1^m + K^m}$.

###### Ordinary Differential Equations

$$
\begin{aligned}
\dot{y}_1 &= \alpha \frac{y_2^m}{y_2^m + K^m} - \delta y_1 - \beta u_2 \frac{y_1}{y_1 + K} && (49) \\
\dot{y}_2 &= \alpha \frac{y_1^m}{y_1^m + K^m} - \delta y_2 && (50) \\
\dot{u}_1 &= kx - \delta u_1 - \gamma u_1 u_2 && (51) \\
\dot{u}_2 &= \xi \frac{y_1^m}{K^m + y_1^m} - \delta u_2 - \gamma u_1 u_2 && (52) \\
\dot{x} &= \theta \frac{y_1^m}{K^m + y_1^m} - \delta x && (53)
\end{aligned}
$$

###### Steady-state analysis

To determine the system´s phase plane, we calculate the nullclines by equating $\dot{y}_1 = 0$ and $\dot{y}_2 = 0$, obtaining the following expressions:

$$
\begin{aligned}
\bar{y}_1 &= \sqrt[m]{\frac{\delta \bar{y}_2 K^m}{\alpha - \delta \bar{y}_2}} \\
\bar{y}_2 &= \sqrt[m]{\frac{h K^m}{\alpha - h}}
\end{aligned}
$$

where $h = \delta \bar{y}_1 + \beta \bar{u}_2 \frac{\bar{y}_1}{\bar{y}_1 + K}$. We can further find $\bar{u}_2$ as a function of $\bar{y}_1$,

$$
\bar{u}_1 = \frac{f_2 - \delta \bar{u}_2}{\gamma \bar{u}_2} = \frac{f_1}{\gamma \bar{u}_2 + \delta},
$$

where $f_1 = k\bar{x}$, $f_2 = \xi \frac{y_1^m}{K^m + \bar{y}_1^m}$, and $\bar{x} = \frac{\theta}{\delta} \frac{y_1^m}{K^m + \bar{y}_1^m}$. This leads to a second order polynomial, $P(\bar{u}_2) = \bar{u}_2^2 + A\bar{u}_2 + B = 0$, and leads to a single positive real solution

$$
\bar{u}_2 = \frac{-A + \sqrt{A^2 - 4B}}{2}
$$

where $A = \frac{f_1 - f_2}{\delta} + \frac{\delta}{\gamma}$, and $B = -\frac{f_2}{\gamma}$.

##### 5.5.4 Double inhibiting controller with actuation through $U_1$ and $U_1'$ species

The species that comprise the controller that actuates over the $Y_1$ specie are named $U_1$, $U_2$ and $X$, while the species that comprise the controller that actuates over the $Y_2$ specie are named $U_1'$, $U_2'$ and $X'$. We assumed the same kinetic constants ($\delta$, $k$, $\xi$, $\theta$ and $\gamma$) for both controllers, except for the actuation gain, which has the same notation as detailed before ($\beta$ and $\beta'$).

###### Chemical reactions

| | | |
|---|---|---|
| $\emptyset \xrightarrow{\alpha p_1} Y_1,$ | $\emptyset \xrightarrow{\alpha p_2} Y_2,$ | Production, |
| $Y_1 \xrightarrow{\delta} \emptyset,$ | $Y_2 \xrightarrow{\delta} \emptyset,$ | Decay |
| $Y_1 \xrightarrow{\theta p_2} Y_1 + X$ | $Y_1 \xrightarrow{\xi p_2} Y_1 + U_2$ | Production |
| $U_1 + Y_1 \xrightarrow{\beta} \emptyset$ | $U_1 + U_2 \xrightarrow{\gamma} \emptyset$ | Decay/Sequestration |
| $U_1, U_2 \xrightarrow{\delta} \emptyset$ | $X \xrightarrow{\delta} \emptyset$ | Decay |
| $Y_2 \xrightarrow{\theta p_1} Y_2 + X'$ | $Y_2 \xrightarrow{\xi p_1} Y_2 + U_2'$ | Production |
| $U_1' + Y_2 \xrightarrow{\beta'} \emptyset$ | $U_1' + U_2' \xrightarrow{\gamma} \emptyset$ | Decay/Sequestration |
| $U_1', U_2' \xrightarrow{\delta} \emptyset$ | $X' \xrightarrow{\delta} \emptyset$ | Decay |

where production rate constants are defined as $p_1 = \frac{y_2^m}{y_2^m + K^m}$, and $p_2 = \frac{y_1^m}{y_1^m + K^m}$.

###### Ordinary Differential Equations

$$\dot{y}_1 = \alpha \frac{y_2^m}{y_2^m + K^m} - \delta y_1 - \beta u_1 \frac{y_1}{y_1 + K} \quad (54)$$

$$\dot{y}_2 = \alpha \frac{y_1^m}{y_1^m + K^m} - \delta y_2 - \beta u_1' \frac{y_2}{y_2 + K} \quad (55)$$

$$\dot{u}_1 = kx - \delta u_1 - \gamma u_1 u_2 \quad (56)$$

$$\dot{u}_2 = \xi \frac{y_1^m}{K^m + y_2^m} - \delta u_2 - \gamma u_1 u_2 \quad (57)$$

$$\dot{x} = \theta \frac{y_1^m}{K^m + y_2^m} - \delta x \quad (58)$$

$$\dot{u}_1' = kx' - \delta u_1' - \gamma u_1' u_2' \quad (59)$$

$$\dot{u}_2' = \xi \frac{y_2^m}{K^m + y_2^m} - \delta u_2' - \gamma u_1' u_2' \quad (60)$$

$$\dot{x}' = \theta \frac{y_2^m}{K^m + y_2^m} - \delta x' \quad (61)$$

###### Steady-state analysis

To determine the system´s phase plane, we calculate the nullclines by equating $\dot{y}_1 = 0$ and $\dot{y}_2 = 0$, obtaining the following expressions:

$$
\begin{aligned}
\bar{y_1} &= \sqrt[m]{\frac{h_1 K^m}{\alpha - h_1}} \\
\bar{y_2} &= \sqrt[m]{\frac{h_2 K^m}{\alpha - h_2}}
\end{aligned}
$$

where $h_1 = \delta\bar{y_2} + \beta\bar{u_1}'\frac{\bar{y_2}}{\bar{y_2}+K^m}$ and $h_2 = \delta\bar{y_1} + \beta\bar{u_1}\frac{\bar{y_1}}{\bar{y_1}+K^m}$. As detailed before, $\bar{u_1}$ and $\bar{u_1}'$ can be expressed as functions of $\bar{y_1}$ and $\bar{y_2}$, respectively.

#### 5.6 Adaptive controller with negative feedback for the toggle switch with self-induction case study

These are the networks studied in the subsection *2.3.2 Applying the adaptive control principle to multi-stable systems* in the main text, and characterized in Fig. S21.

##### 5.6.1 Double inhibiting controller with actuation through $U_1$ and $U_1'$ specie

**Chemical reactions**

| | | |
|---|---|---|
| $\emptyset \xrightarrow{\alpha(p_1,p_3)} Y_1,$ | $\emptyset \xrightarrow{\alpha(p_2,p_4)} Y_2,$ | Production, |
| $Y_1 \xrightarrow{\delta} \emptyset,$ | $Y_2 \xrightarrow{\delta} \emptyset,$ | Decay |
| $Y_1 \xrightarrow{\theta p_2} Y_1 + X$ | $Y_1 \xrightarrow{\xi p_2} Y_1 + U_2$ | Production |
| $U_1 + Y_1 \xrightarrow{\beta} \emptyset$ | $U_1 + U_2 \xrightarrow{\gamma} \emptyset$ | Decay/Sequestration |
| $U_1, U_2 \xrightarrow{\delta} \emptyset$ | $X \xrightarrow{\delta} \emptyset$ | Decay |
| $Y_2 \xrightarrow{\theta p_1} Y_2 + X'$ | $Y_2 \xrightarrow{\xi p_1} Y_2 + U_2'$ | Production |
| $U_1' + Y_2 \xrightarrow{\beta'} \emptyset$ | $U_1' + U_2' \xrightarrow{\gamma} \emptyset$ | Decay/Sequestration |
| $U_1', U_2' \xrightarrow{\delta} \emptyset$ | $X' \xrightarrow{\delta} \emptyset$ | Decay |

where production rate constants are defined as $r_1 = \alpha\frac{K^m}{y_2^m+K^m}$, $r_2 = \alpha\frac{K^m}{y_1^m+K^m}$, $r_3 = \alpha\frac{y_1^m}{y_1^m+K^m}$ and $r_4 = \alpha\frac{y_2^m}{y_2^m+K^m}$

##### Ordinary Differential Equations

$$
\begin{aligned}
\dot{y}_1 &= \alpha\frac{K^m}{y_2^m+K^m}+\alpha\frac{y_1^m}{y_1^m+K^m}-\delta y_1-\beta u_1\frac{y_1}{y_1+K} && (62)\\
\dot{y}_2 &= \alpha\frac{K^m}{y_1^m+K^m}+\alpha\frac{y_2^m}{y_2^m+K^m}-\delta y_2-\beta u_1'\frac{y_2}{y_2+K} && (63)\\
\dot{u}_1 &= kx-\delta u_1-\gamma u_1u_2 && (64)\\
\dot{u}_2 &= \xi\frac{K^m}{K^m+y_1^m}-\delta u_2-\gamma u_1u_2 && (65)\\
\dot{x} &= \theta\frac{K^m}{K^m+y_1^m}-\delta x && (66)\\
\dot{u}_1' &= kx'-\delta u_1'-\gamma u_1'u_2' && (67)\\
\dot{u}_2' &= \xi\frac{K^m}{K^m+y_2^m}-\delta u_2'-\gamma u_1'u_2' && (68)\\
\dot{x}' &= \theta\frac{K^m}{K^m+y_2^m}-\delta x' && (69)
\end{aligned}
$$

##### Steady-state analysis

To determine the system´s phase plane, we calculate the nullclines by equating $\dot{y}_1 = 0$ and $\dot{y}_2 = 0$, obtaining the following expressions:

$$
\begin{aligned}
\bar{y_1} &= \sqrt[m]{\frac{\alpha K^m}{h_1}-K^m}\\
\bar{y_2} &= \sqrt[m]{\frac{\alpha K^m}{h_2}-K^m}
\end{aligned}
$$

where $h_1 = \delta\bar{y_2}+\frac{\bar{y_2}^m}{\bar{y_2}^m+K^m}(\beta\bar{u_1}'-\alpha)$ and $h_2 = \delta\bar{y_1}+\frac{\bar{y_1}^m}{\bar{y_1}^m+K^m}(\beta\bar{u_1}-\alpha)$. As detailed before, $\bar{u_1}$ and $\bar{u_1}'$ can be expressed as functions of $\bar{y_1}$ and $\bar{y_2}$, respectively.

##### 5.6.2 Inhibition-induction pair

##### Chemical reactions

| | | |
|---|---|---|
| $\emptyset \xrightarrow{\alpha(p_1,p_3)} Y_1,$ | $\emptyset \xrightarrow{\alpha(p_2,p_4)} Y_2,$ | Production, |
| $Y_1 \xrightarrow{\delta} \emptyset,$ | $Y_2 \xrightarrow{\delta} \emptyset,$ | Decay |
| $Y_1 \xrightarrow{\theta p_2} Y_1+X$ | $Y_1 \xrightarrow{\xi p_2} Y_1+U_2$ | Production |
| $U_1 \xrightarrow{\beta} U_1+Y_1$ | $U_1+U_2 \xrightarrow{\gamma} \emptyset$ | Production/Sequestration |
| $U_1, U_2 \xrightarrow{\delta} \emptyset$ | $X \xrightarrow{\delta} \emptyset$ | Decay |
| $Y_2 \xrightarrow{\theta p_1} Y_2+X'$ | $Y_2 \xrightarrow{\xi p_1} Y_2+U_2'$ | Production |
| $U_1'+Y_2 \xrightarrow{\beta'} \emptyset$ | $U_1'+U_2' \xrightarrow{\gamma} \emptyset$ | Decay/Sequestration |
| $U_1', U_2' \xrightarrow{\delta} \emptyset$ | $X' \xrightarrow{\delta} \emptyset$ | Decay |

where production rate constants are defined as $p_1 = \frac{K^m}{y_2^m+K^m}$, $p_2 = \frac{K^m}{y_1^m+K^m}$, $p_3 = \frac{y_1^m}{y_1^m+K^m}$ and $p_4 = \frac{y_2^m}{y_2^m+K^m}$

#### Ordinary Differential Equations

$$\begin{aligned}
\dot{y}_1 &= \alpha\frac{K^m}{y_2^m + K^m} + \alpha\frac{y_1^m}{y_1^m + K^m} - \delta y_1 + \beta u_1 && (70)\\
\dot{y}_2 &= \alpha\frac{K^m}{y_1^m + K^m} + \alpha\frac{y_2^m}{y_2^m + K^m} - \delta y_2 - \beta u_1'\frac{y_2}{y_2 + K} && (71)\\
\dot{u}_1 &= kx - \delta u_1 - \gamma u_1 u_2 && (72)\\
\dot{u}_2 &= \xi\frac{K^m}{K^m + y_1^m} - \delta u_2 - \gamma u_1 u_2 && (73)\\
\dot{x} &= \theta\frac{K^m}{K^m + y_1^m} - \delta x && (74)\\
\dot{u}_1' &= kx' - \delta u_1' - \gamma u_1' u_2' && (75)\\
\dot{u}_2' &= \xi\frac{K^m}{K^m + y_2^m} - \delta u_2' - \gamma u_1' u_2' && (76)\\
\dot{x}' &= \theta\frac{K^m}{K^m + y_2^m} - \delta x' && (77)
\end{aligned}$$

#### Steady-state analysis

To determine the system´s phase plane, we calculate the nullclines by equating $\dot{y}_1 = 0$ and $\dot{y}_2 = 0$, obtaining the following expressions:

$$\begin{aligned}
\bar{y}_1 &= \sqrt[m]{\frac{h_1 K^m}{\alpha - h_1}}\\
\bar{y}_2 &= \sqrt[m]{\frac{h_2 K^m}{\alpha - h_2}}
\end{aligned}$$

where $h_1 = \delta\bar{y}_2 + \beta\bar{u}_1'\frac{\bar{y}_2}{\bar{y}_2 + K} - \alpha\frac{\bar{y}_2^m}{\bar{y}_2^m + K^m}$ and $h_2 = \delta y_1 - \beta\bar{u}_1 - \alpha\frac{\bar{y}_1^m}{\bar{y}_1^m + K^m}$. As detailed before, $\bar{u}_1$ and $\bar{u_1}'$ can be expressed as functions of $\bar{y}_1$ and $\bar{y}_2$, respectively.
